# Evolution of dominance in a Mendelian trait is linked to the evolution of environmental plasticity

**DOI:** 10.1101/2025.05.30.657093

**Authors:** Yuichi Fukutomi, Alexandra Phillips-Garcia, Jingqi Liu, Ashley Chuang, Masayoshi Watada, Seema Ramniwas, Artyom Kopp

**Affiliations:** Department of Evolution and Ecology, University of California - Davis, Davis, CA, USA; Evergreen Valley High School, San Jose, CA, USA; Department of Biological Sciences, Tokyo Metropolitan University, Hachioji, Tokyo, Japan; Marwadi University Research Center, Faculty of Sciences, Marwadi University, Rajkot 360003, Gujarat, India

**Author notes:** The corresponding author (email address, mailing address: One Shields Ave, Davis, CA 95616, USA). Department of Biology, University of Virginia, Charlottesville, VA, USA.

**Keywords:** Evolution of dominance, evolution of phenotypic plasticity, sex-limited polymorphism, *Drosophila* pigmentation, developmental thresholds, genotype-phenotype mapping

## Abstract

Allelic dominance and phenotypic plasticity both influence how genetic variation is expressed in phenotypes, shaping evolutionary responses to selection. In both cases, changes in genotype or environment can cause sharp, nonlinear phenotypic shifts, hinting at shared underlying features of development that may link dominance and plasticity. Here, we investigate these links using a Mendelian, female-limited color dimorphism found in many species of the *Drosophila montium* lineage. In most species, the Dark allele is dominant, but two species—*D. jambulina* and *D.* cf. *bocqueti*—have been reported to have dominant Light alleles. We show that in both Dark-dominant and Light-dominant species, the color dimorphism is linked to the same locus: the *POU domain motif 3* (*pdm3*) transcription factor. We then demonstrate that the interspecific differences in dominance relationships between *pdm3* alleles are due to changes in phenotypic plasticity. In the Dark-dominant species *D. rufa* and *D. burlai*, the Dark allele is dominant across all developmental temperatures. In contrast, in both Light-dominant species, dominance is temperature-dependent, with the Light allele becoming increasingly dominant at higher temperatures. These results suggest a mechanistic connection between the evolution of dominance and phenotypic plasticity. We propose this connection may emerge from threshold-like properties of developmental systems.

**Teaser text:** This study reveals a direct relationship between the evolution of dominance and phenotypic plasticity in a Mendelian, female-limited color dimorphism. This trait is linked to the same transcription factor gene, *pdm3*, in multiple species of the *Drosophila montium* lineage. However in some species, the Light *pdm3* allele becomes increasingly dominant at higher temperature, while in others the Dark allele is fully dominant under all conditions. Based on these results, we propose that the evolution of dominance and plasticity may be connected by threshold-like properties of developmental systems.

## Introduction

Allelic dominance has a profound influence on evolution by shaping the genotype-phenotype relationship. By determining the phenotype of heterozygotes, dominance affects both the maintenance of genetic variation and the course of directional selection (Prout, 1971; Lewontin, 1974; Cook, 2003; Hedrick, 2011). Not surprisingly, the reasons why some alleles are dominant, and others are recessive have been debated since the earliest days of population genetics (Fisher, 1928; Wright, 1934; Kacser & Burns, 1981; Gilchrist & Nijhout, 2001; Porter et al., 2017). A key question is whether dominance is an inherent property of alleles or an outcome of natural selection. Fisher (1928) hypothesized that a wild-type allele will become dominant due to selection on modifier alleles at other loci in order to reduce the effects of deleterious mutations in heterozygotes. Other models suggest that dominance reflects intrinsic properties of biological systems such as metabolic flux, spatial diffusion of proteins, or DNA-protein binding, and therefore arises without the need for selection (Wright, 1934; Kacser & Burns, 1981; Gilchrist & Nijhout, 2001; Porter et al., 2017).

Whatever the mechanistic causes of dominance may be, dominance relationships between alleles are not constant; they can change over time and diverge between populations and species (Clarke & Sheppard, 1960; Kettlewell, 1965; Rosenblum et al., 2010; Le Poul et al., 2014). The evolution of dominance contributes, for example, to adaptation to the changing environment, predation avoidance in Mullerian mimics, and the resolution of sexual conflict (Haldane, 1956; Sheppard, 1975; Johnston et al., 2013; Barson et al., 2015; Arias et al., 2016). Theoretical models predict that differences in allelic dominance can evolve by selection acting either directly on the focal locus (Billiard et al., 2021), or on second-site modifier loci, e.g. via frequency-dependent selection or sexually antagonistic selection (Peischl & Bürger, 2008; Spencer & Priest, 2016).

Going beyond theoretical models and identifying the molecular mechanisms underlying the evolution of dominance has proven a difficult challenge. In one of the few examples, different mutations in the coding sequence of the melanocortin receptor *Mc1r* gene have similar phenotypic effects in homozygotes, but different dominance in heterozygotes, in two lizard species because they affect different biochemical processes: disrupting receptor signaling in one versus reducing membrane integration efficiency in the other (Rosenblum et al., 2010). Here, dominance appears to be an inherent property of the *Mc1r* alleles themselves. An example where dominance relationships between alleles depend on second-site modifiers and may be affected by selection is found in the evolution of self-incompatibility in *Brassica rapa*. In this case, polymorphic trans-acting small RNAs make some alleles at the self-incompatibility locus dominant by silencing the expression of recessive alleles (Tarutani et al., 2010; Yasuda et al., 2016). Even this limited comparison shows that the molecular mechanisms behind the evolution of dominance are clearly disparate in different systems, and it is likely that other mechanisms remain to be discovered (Billiard et al., 2021).

In addition to dominance, the genotype-phenotype relationship can be affected by phenotypic plasticity. Since the optimal phenotype often varies between environments, the ability of a single genotype to produce different phenotypes in response to environmental cues is an important aspect of adaptation (West-Eberhard, 2003; Whitman & Agrawal, 2009; Moczek, 2010; Sommer, 2020). As with allelic dominance, phenotypic plasticity has been proposed to be either a result of selection for adaptation to variable environments, or an intrinsic property of developmental pathways that, by default, have low robustness (West-Eberhard, 2003; de Jong, 2005; Pigliucci, 2005; Bhardwaj et al., 2020). The two explanations are not mutually exclusive, and regardless of the origin of plasticity, it can have significant costs as well as benefits. Most generally, the maintenance of regulatory mechanisms that enable plasticity may require energy and resources, while the expression of plasticity often involves fitness trade-offs (DeWitt, 1998; Auld et al., 2010; Kelly et al., 2012; Murren et al., 2015). For example, induction of defensive traits in response to predation can be negatively correlated with growth rates, and may increase the risks posed by functionally different predators (Hoverman & Relyea, 2009; Bennet et al., 2013; Brönmark et al., 2012). Plasticity may also limit adaptation to any single environment, and can produce phenotype-environment mismatches when environmental cues become less predictive (Ghalambor et al., 2007; Oostra et al., 2018; Rodriguez & Beldade, 2020; Schneider, 2022). Natural selection can therefore act on the magnitude of phenotypic plasticity, which can diverge between populations and species (West-Eberhard, 2003; Lind & Johansson, 2007; Davidson et al., 2011; Chevin & Hoffmann, 2017; Snell-Rood & Ehlman, 2021).

Since phenotypic plasticity is, by definition, a one (genotype) to many (phenotypes) relationship, it is mediated by genes whose expression depends on environmental conditions. What these genes are, how environmental cues regulate their expression, and what genetic changes explain the evolution of plasticity, are key questions in evolutionary biology (Moczek, 2010; Kelly et al., 2012; Sommer, 2020). Some of the best-studied examples involve hormone-mediated effects of temperature on development. For example, the tobacco hornworm, *Manduca sexta,* is normally monophenic, while its relative *M. quinquemaculata* shows color polyphenism in response to temperature. A mutation in *M. sexta* that reduces juvenile hormone (JH) production results in temperature-dependent polyphenism similar to that seen in *M. quinquemaculata,* and both the temperature reaction norms and JH titers can subsequently be changed by selection (Suzuki & Nijhout, 2006). However, neither the loci responsible for the difference in phenotypic plasticity between the two species, nor the loci responsible for the evolution of reaction norms in response to selection, have been identified. In the nymphalid butterfly *Bicyclus anynana,* lower temperatures lead to lower titers of 20-hydroxyecdysone (20E), which in turn cause the development of smaller wing eyespots (Bhardwaj et al., 2018). In other nymphalids, temperature does not affect eyespot size despite having similar effects on 20E titers, suggesting a recent origin of thermal plasticity (Bhardwaj et al., 2020) – however, the genetic changes responsible for this transition are also unknown.

A potential common thread linking dominance and plasticity is developmental robustness: in a sense, a fully dominant allele is robust against genetic variation, while a completely non-plastic genotype is robust against environmental variation. Both types of robustness can be caused by regulatory nonlinearities during development – that is, nonlinear mapping of the phenotypic outcome to its underlying determinants such as gene expression, protein levels, or hormone titers (Suzuki & Nijhout, 2006; Hallgrimsson et al., 2019). For example, environmentally plastic, hormone-dependent phenotypes often act as threshold traits: no phenotypic changes are observed unless changes in the hormone titer cross a genetically determined threshold, which may differ between populations or species (Suzuki & Nijhout, 2006; Bhardwaj et al., 2018). Nonlinear genotype-phenotype relationships can also stem from the properties of DNA-protein binding, regulatory feedback loops, or threshold responses to morphogen gradients (Hill, 1910; Steinacher et al., 2016; Bottani & Veitia, 2017; Hallgrimsson et al., 2019). Importantly, threshold-like behavior of biological systems may also explain dominance relationships between alleles and the evolution of dominance (Omholt et al., 2000; Gilchrist & Nijhout, 2001; Reid, 2022). Thus, there may be a deep mechanistic connection between dominance and phenotypic plasticity.

In this study, we examined interspecific differences in the dominance relationships between alleles at a Mendelian locus responsible for a female-limited color dimorphism in the *Drosophila montium* species subgroup (Fig. 1). In multiple species of this clade, two alternative female morphs, light and dark, are distinguished by the color of posterior abdominal segments, whereas males are either monomorphic light or monomorphic dark depending on the species. This female polymorphism is controlled by alternative alleles of the *POU domain motif 3* (*pdm3*) gene, which encodes a transcription factor that represses melanin production (Yassin et al., 2016). In most species, the Dark allele of *pdm3* is dominant (Ohnishi & Watanabe, 1985; Yassin et al., 2016). However, light pigmentation has been described as dominant in *D. jambulina* (Ohnishi & Watanabe, 1985), and in strains of *D. bocqueti* from Sao Tome (Prigent et al. 2020). Here, we show that the key difference between “light-dominant” and “dark-dominant” species is that the dominance relationships between *pdm3* alleles are temperature-dependent in the former, but invariant in the latter. These results suggest that the evolution of dominance can be directly related to the evolution of phenotypic plasticity, and vice versa.

**Figure 1.**
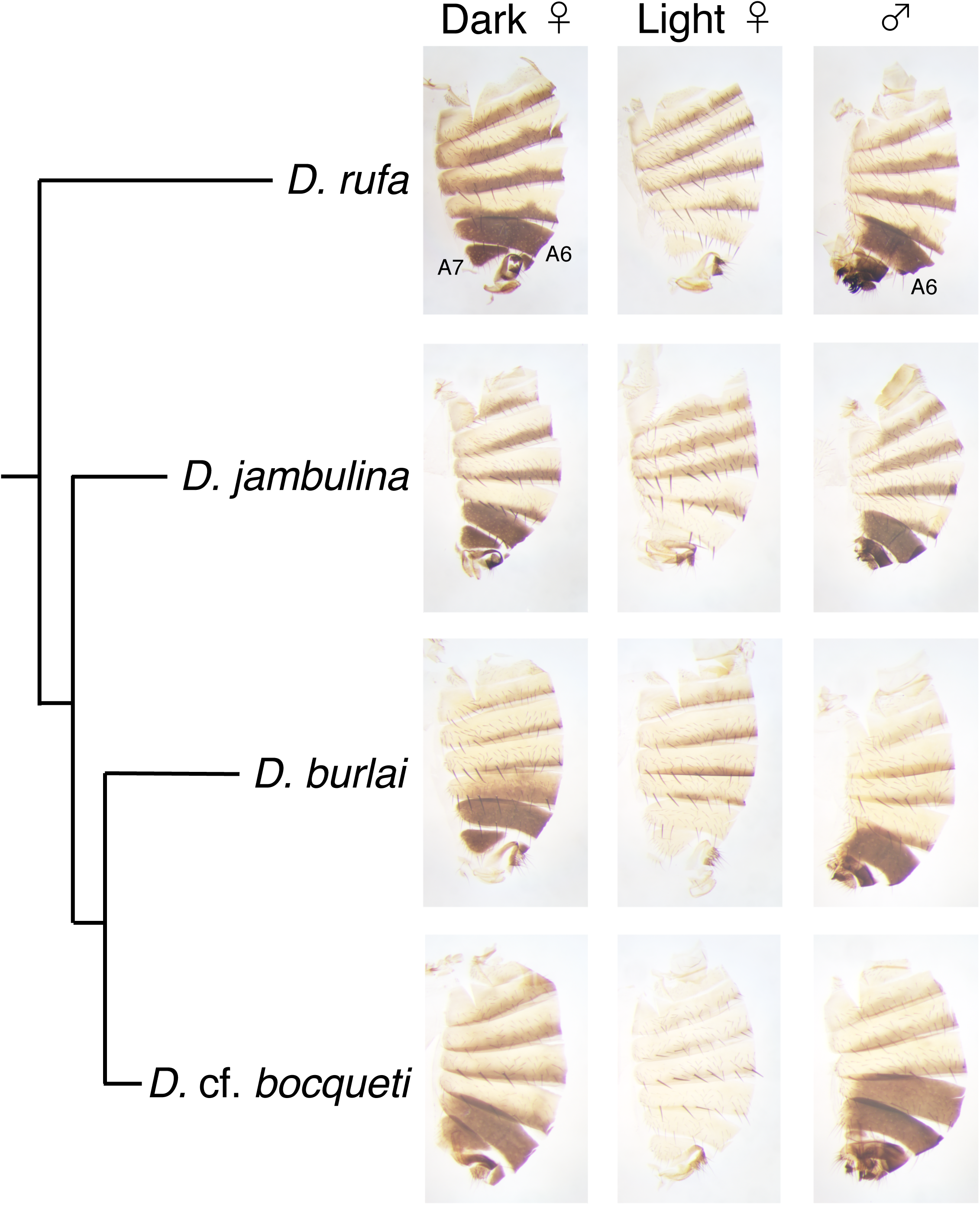
Abdominal pigmentation in four species of the *Drosophila montium* species subgroup (*melanogaster* species group) used in this study. In each species, females show a Mendelian color dimorphism in the A6 segment (Dark / fully pigmented vs Light / not pigmented) while males are monomorphic with dark A6. The phylogeny is based on Conner et al (2021), and branch lengths are not to scale.

## Materials and Methods

### Drosophila strains

The species examined in this study were *D. rufa*, *D. jambulina*, *D. burlai*, *D. chauvacae,* and *D.* cf. *bocqueti*, all belonging to the *montium* species subgroup (see Table 1 and Supplement Tables S1-S2 for strain information). *D.* cf. *bocqueti* is morphologically similar to *D. bocqueti*, but the taxonomic identity of *D. bocqueti* requires further investigation (A. Yassin, pers. comm.). *D.* cf. *bocqueti* is closely related to *D. burlai* and *D. chauvacae* (Conner et al. 2021). Flies were reared on standard *Drosophila* cornmeal media in controlled temperature incubators between 17°C and 26°C under 12:12 hour day/night cycle. Not all species were viable over the full range of temperatures (Supplement Table S3).

**Table 1.**
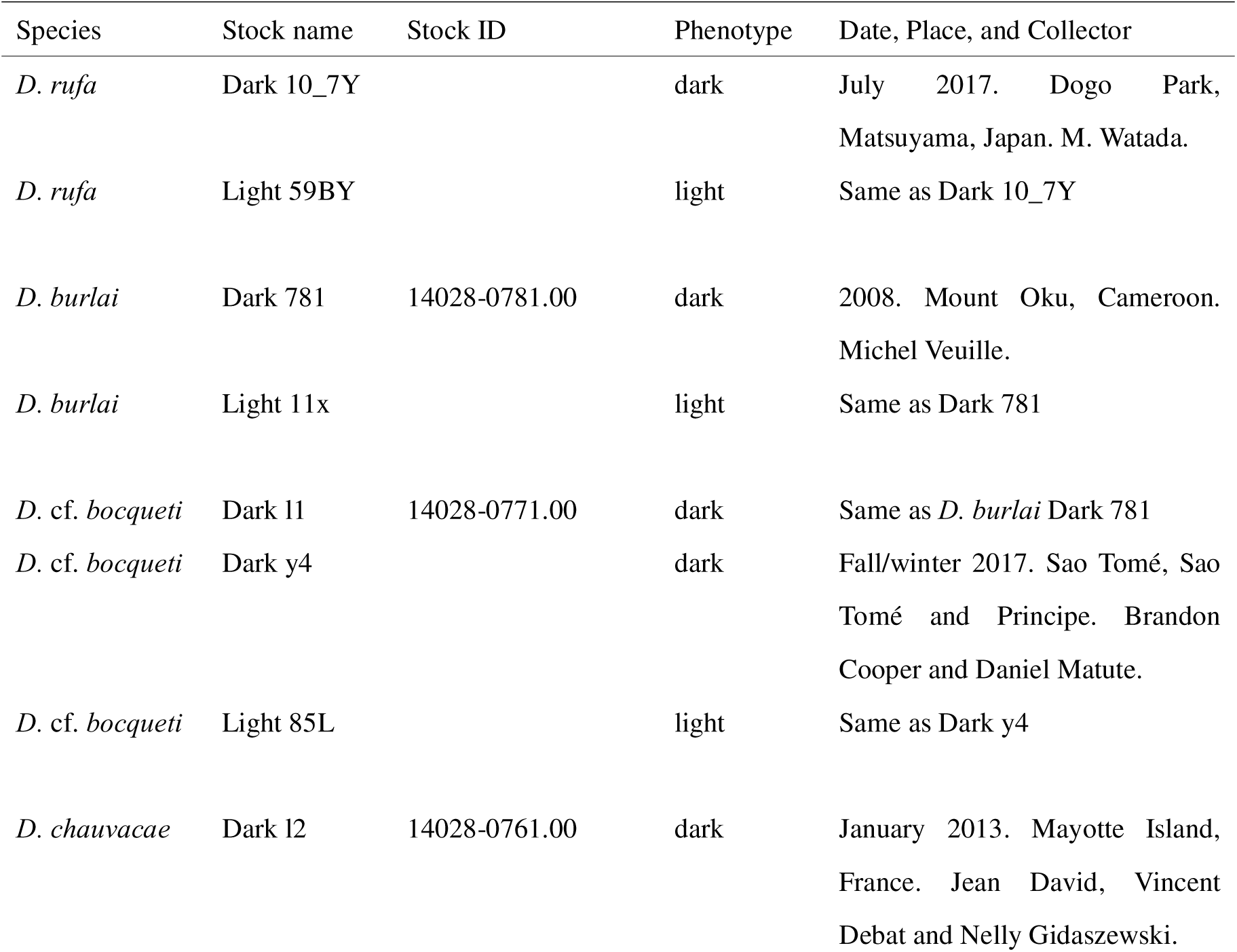
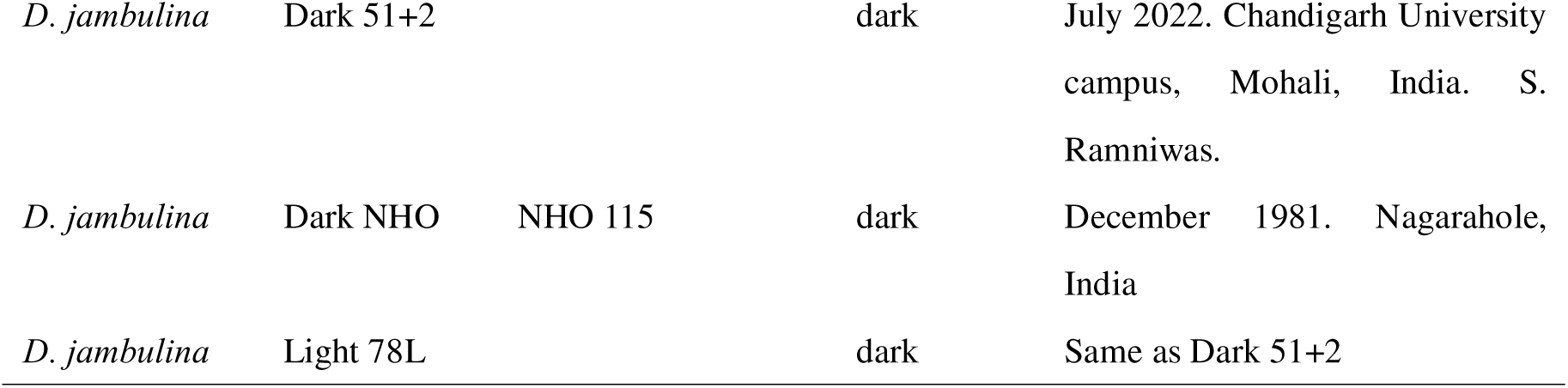
Strains used to examine dominance and plasticity. “Phenotype” column indicates the phenotype of females. For Dark 781 strain of *D. burlai*, Dark l1 strain of *D.* cf. *bocqueti*, and Dark l2 strain of *D. chauvacae,* the Stock ID column refers to the US National *Drosophila* Species Stock Center, where they were obtained. The Dark NHO strain of *D. jambulina* was provided by the Ehime *Drosophila* Stock Center (Japan). Additional strains used for genetic mapping are listed in Supplementary Tables S1-S2.

### Testing for linkage between female pigmentation and *pdm3*

As female-limited color dimorphism maps to the *pdm3* locus in multiple species of the *montium* species subgroup including *D. burlai* (Yassin et al., 2016), we tested whether *pdm3* was also associated with this trait in *D. rufa*, *D.* cf. *bocqueti,* and *D. jambulina*. In *D. rufa*, we used diagnostic PCR to genotype 43 isofemale strains collected from the same local population (Supplement Table S1). First, we used whole-genome sequences of a subset of these strains to identify diagnostic differences between light and dark strains in the first intron (21 kb) of *pdm3*. Based on these differences, we designed PCR primers that were specific to the Dark versus Light *pdm3* alleles. The forward primer CCATACCATACAAGCGCCATCTAG and the reverse primer TTATCATGCATCAATGCGACAGCAAC amplified a 1.5 kb fragment from the Dark allele, but not from the Light allele. The same forward primer in combination with the AGCGACACAGATAACCAGATTTCGA reverse primer amplified a 656 bp fragment from the Light allele, but not from the Dark allele (Supplement Fig. S1; Supplement Table S1). These two primer pairs, which allowed us to distinguish Dark and Light homozygotes and Dark/Light heterozygotes at the *pdm3* locus, were used to genotype 21 dark and 22 light females of *D. rufa* by diagnostic PCR.

In *D. jambulina*, we collected 19 isofemale strains from the same local population; in addition, the *Dark NHO* strain was obtained from the Ehime stock center (Supplement Table S2). Since most of these strains were polymorphic for female pigmentation, we genotyped dark virgin females from 13 different strains and light virgin females from 13 different strains by amplifying and sequencing a fragment of the *pdm3* locus. To maximize the proportion of *pdm3* homozygotes among the flies used for genotyping, dark females were isolated from cultures that were raised at 25°C (where light pigmentation is dominant), and light females were isolated from cultures raised at 17°C (where dark pigmentation is dominant). The intronic region of *pdm3* homologous to the structural variant that is responsible for female color polymorphism in *D. serrata* (Yassin et al., 2016) was amplified with the forward primer TAATTACACCCGCCAAAGGGCAC and the reverse primer CTCTCTACTTATCTGGGGTTGCAAAG and sequenced using the same forward primer or the alternative primer TGCAAGCTGAGATTACACTGAGGGA. For genotyping, the amplicon sequences were aligned with MEGA11 (Tamura et al., 2021). As we found an insertion specific to the Light *pdm3* allele within the sequenced region, we conducted diagnostic PCR on six additional strains of *D. jambulina*. The forward primer CAGTTTTTATGACTATGCAAATCAGGACTAG containing the Light allele specific insertion at the 3’ end and the reverse primer CTCTCTACTTATCTGGGGTTGCAAAG amplified 456 bp fragment from the Light allele, but not from the Dark allele (Supplement Fig. S2 A). The same reverse primer in combination with the forward primer ACCACCGTTGATTATGCCAGGTTAG containing Dark allele specific single nucleotide polymorphisms (SNPs) at the 3’ end, amplified a 625 bp fragment from the Dark allele, but not from the Light allele (Supplement Fig. S2 B).

In *D.* cf. *bocqueti*, where only a few strains (a gift from B. Cooper and D. Matute) are available, we generated an introgression strain by crossing the pure-breeding *Light 85L* strain to the pure-breeding *Dark l1* strain, selecting light hybrid females, and backcrossing them to males of the *Dark l1* strain. This backcrossing procedure was repeated for 10 generations, resulting in the introgression of the gene(s) responsible for light pigmentation into an otherwise dark genetic background. Four dark females from the *Dark l1* strain, four light females from the *Light 85L* strain, 19 dark virgin females from the introgression line, and 18 light virgin females from the introgression line were then genotyped for *pdm3*. To maximize the proportion of *pdm3* homozygotes, the introgression line was reared at 25°C, and virgin females with the darkest and lightest pigmentation were selected for genotyping. A 770 bp region containing Exon 11 of *pdm3* was amplified with the forward primer ACCGAGAGAGAGCCGCGTTTC and the reverse primer CTCGGGTTCTGTGCTTCGCTTG, and the amplicons were sequenced with the primer GCTGGACATCACACCAAAGA and aligned with MEGA11 (Tamura et al., 2021).

### Quantifying abdominal pigmentation

In each species, we compared the phenotypes of Light and Dark *pdm3* homozygotes and heterozygotes raised over a range of temperatures (Table 1; Supplement Table S3). For intraspecific crosses, we crossed five adult males and five adult virgin females per vial. For interspecific crosses, 30 males and 30 virgin females were crossed. The adults were removed after one week. Adult F1 progeny were collected and fixed in 100% ethanol when they were 6-8 days old. The pigmentation of each abdominal segment was scored on the scale from 0 (completely yellow) to 10 (completely black) as described by David et al. (1990). The scoring was blind, i.e., the scorers did not know the species, genotypes, and rearing temperatures of the flies they scored. Pigmentation scores were summarized in graph form using ggplot2 (Wickham, 2009). Statistical analyses were conducted with R version 4.2.1 (R Core Team, 2022). To test for significant differences between pigmentation scores at different temperatures in the same cross, we used one-way ANOVA and Tukey’s honest significant differences (HSD). To compare reaction norms between different crosses, two-way ANOVA was performed.

### Mounting adult abdominal cuticles

To mount the abdominal cuticles, adult flies were fixed in 100 % ethanol. The abdomens were separated, cut open along the dorsal midline with a razor blade, and incubated in 10 % KOH for 20 minutes to partly digest soft tissues. Most internal tissues were then removed using forceps, the samples were incubated in 10 % KOH for another 40 minutes, and the remaining soft tissues were removed with forceps in water. The resulting cuticles were mounted flat in Hoyer’s medium and imaged with an Excelis MPX-20C camera (ACCU-SCOPE) connected to a Stemi SV11 microscope (ZEISS).

## Results

### Female-limited color dimorphism is linked to *pdm3* in *D.* cf. *bocqueti*, *D. jambulina,* and *D. rufa*

The Mendelian, female-limited color dimorphism is controlled by alternative alleles at the *pdm3* locus in three distantly related species of the *montium* species subgroup – *D. serrata, D. kikkawai*, and *D. burlai* (Yassin et al., 2016). Since this trait also segregates in a Mendelian fashion in other *montium* subgroup species, we tested whether *pdm3* was also responsible for female-limited color dimorphism in *D. rufa*, *D.* cf. *bocqueti*, and *D. jambulina*.

In *D. rufa,* we genotyped 43 isofemale strains sampled from the same natural population; these included pure dark, pure light, and polymorphic strains (Supplement Table S1). Diagnostic PCR showed that all light females only carried the Light *pdm3* allele, whereas all dark females either carried only the Dark *pdm3* allele, or carried both alleles (Table 2; Supplement Table S1). Thus, *pdm3* shows a complete linkage with female pigmentation in this species. Moreover, confirming the original report of Oshima (1952), the dark phenotype, and presumably the Dark *pdm3* allele, is dominant.

**Table 2.**
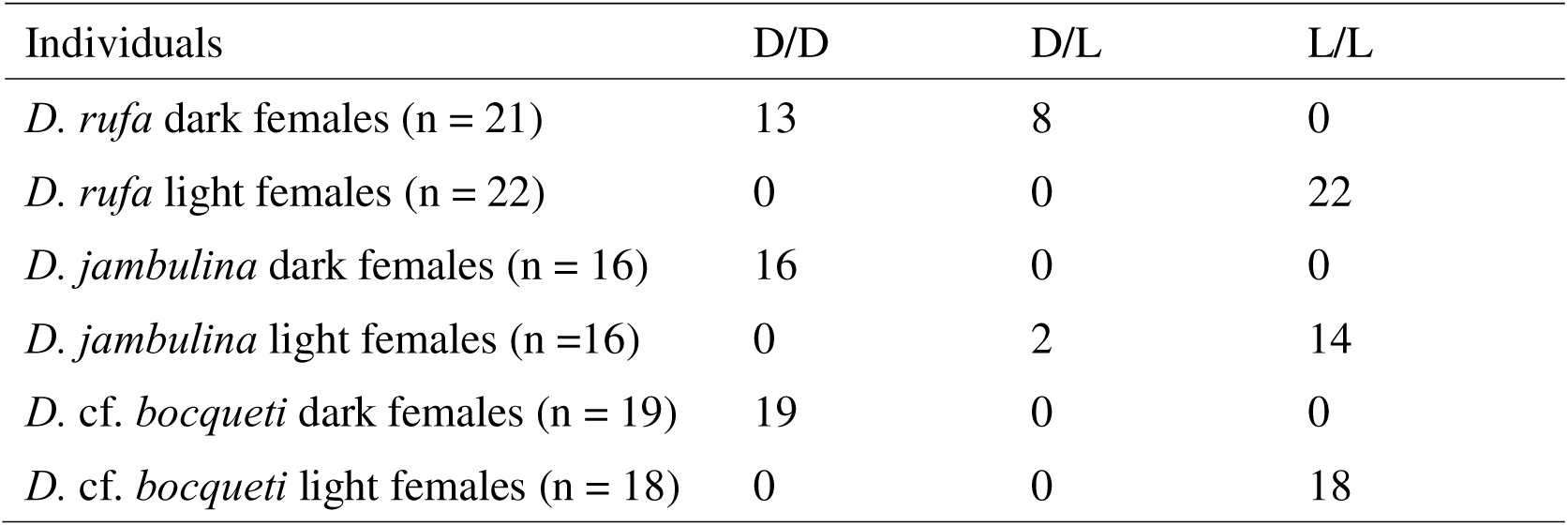
The results of genotyping in *D. rufa*, *D. jambulina*, and *D.* cf. *bocqueti*. The first column indicates the numbers of genotyped individuals for each species and phenotype. Columns 2-4 show the number of flies homozygous for the Dark *pdm3* allele (D/D), heterozygous for *pdm3* (D/L), and homozygous for the Light *pdm3* allele (L/L) among the genotyped individuals. For *D. jambulina* and *D.* cf. *bocqueti*, flies were reared at different temperatures to maximize the proportion of homozygotes among the females selected for genotyping (see Methods).

| Individuals | D/D | D/L | L/L |
| --- | --- | --- | --- |
| *D. rufa* dark females (n = 21) | 13 | 8 | 0 |
| *D. rufa* light females (n = 22) | 0 | 0 | 22 |
| *D. jambulina* dark females (n = 16) | 16 | 0 | 0 |
| *D. jambulina* light females (n = 16) | 0 | 2 | 14 |
| *D. cf. bocqueti* dark females (n = 19) | 19 | 0 | 0 |
| *D. cf. bocqueti* light females (n = 18) | 0 | 0 | 18 |

In *D. jambulina*, we genotyped light and dark females isolated from 19 mostly polymorphic isofemale strains collected from the same natural population, as well as the Dark NHO strain obtained separately (Supplement Table S2). Amplicon sequencing showed that a diagnostic 8-bp indel in the *pdm3* gene (GGACTAGA) was present in all light females, but in none of the dark females (Table 2; Supplement Table S4). Two additional indels and 14 SNPs in the *pdm3* amplicon were also completely linked to the pigmentation phenotype. Additionally, we tested the presence of the diagnostic 8-bp indel in light and dark females collected from another six strains by diagnostic PCR. The diagnostic 8-bp indel was present in all the light females, but in none of the dark females (Supplement Fig. S2). These results indicate that female pigmentation is associated with *pdm3* in *D. jambulina* (Table 2), as well, as that the Light *pdm3* allele is dominant in this species (under standard conditions – see below).

In *D.* cf. *bocqueti*, we genotyped 19 dark and 18 light females from an introgression line in which the light phenotype was introgressed into the dark genetic background. Amplicon sequencing showed that all dark females from the introgression line had the *pdm3* sequence that matched the dark parental allele, while all light introgression females matched the light parental allele of *pdm3* (Table 2; Supplement Table S5).

### The dark *pdm3* allele is fully dominant at all temperatures in *D. rufa* and *D. burlai*

*D. rufa* females homozygous for the dark *pdm3* allele were fully dark (pigmentation score ∼10 in the A6 segment), while those homozygous for the light *pdm3* allele were fully light (pigmentation score 0-1 in the A6 segment), at all rearing temperatures (Fig. 2A). F1 heterozygous females showed dark pigmentation, similar to the dark homozygotes, at all temperatures in both reciprocal crosses (Fig. 2B). Although slight differences between temperatures were detected by one-way ANOVA and Tukey’s HSD test, these differences were very slight in both homo- and heterozygotes (Fig. 2A, B).

**Figure 2.**
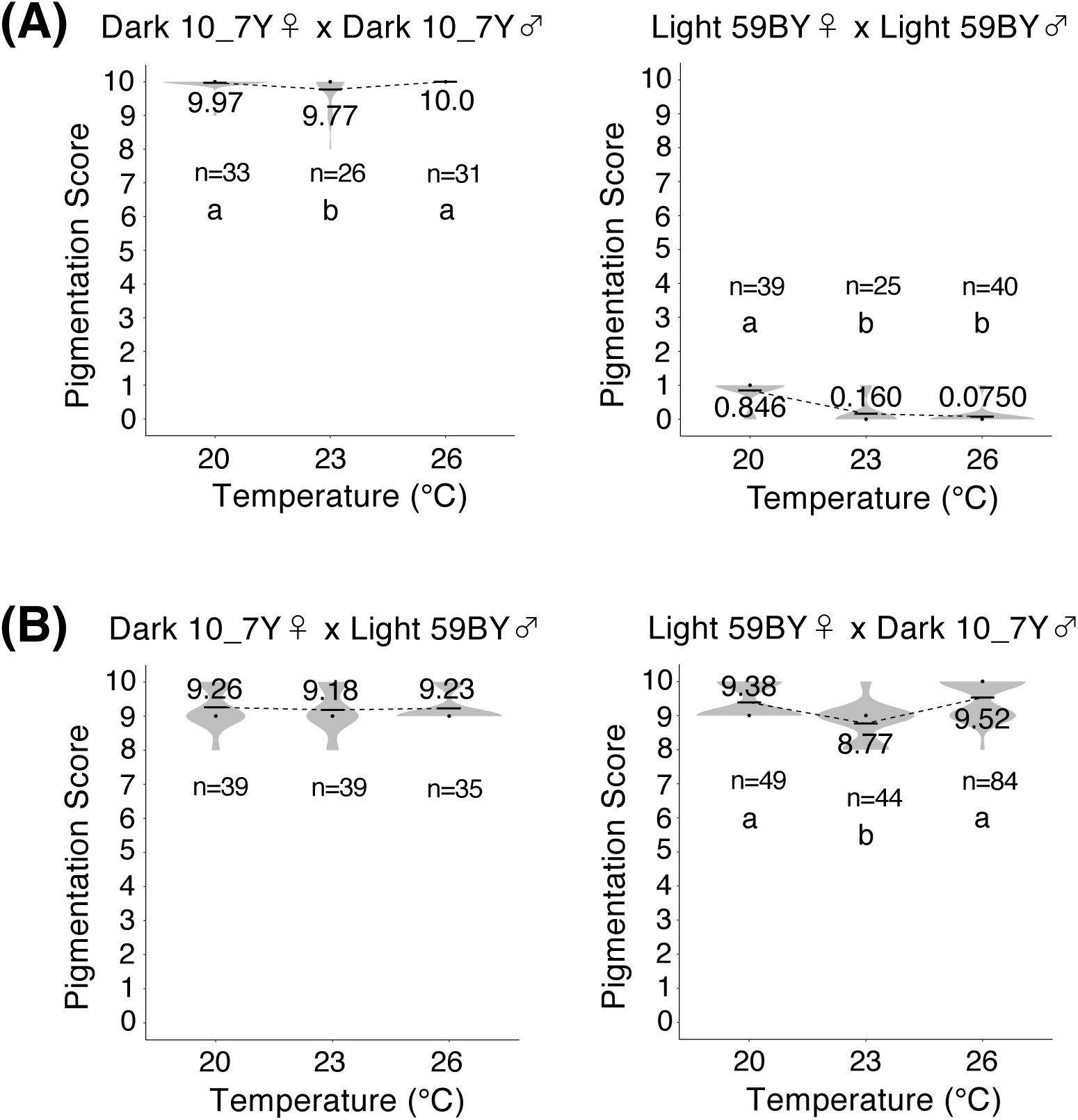
Female A6 pigmentation in *D. rufa* as a function of rearing temperature. (A) The homozygous parental strains Dark 10_7Y (left) and Light 59BY (right). A slight dependence on temperature is observed in each parent (one-way ANOVA, df = 2; *F* = 4.992, *p* = 0.0089 for Dark homozygotes; *F* = 60.37, *p* < 10^-10^ for Light homozygotes). (B) A slight dependence on temperature is observed in the F1 females from the cross between Light females and Dark males (right, *F* = 22.68, *p* < 10^-8^), but not in the reciprocal cross (left, *F* = 0.184). Black bars and black dots indicate mean and median values, respectively; the numbers indicate mean values. Different letters indicate significant differences (*p* < 0.05, Tukey’s HSD test).

Similarly, *D. burlai* females homozygous for the dark *pdm3* allele were fully dark (pigmentation score 10 in the A6 segment), while those homozygous for the light *pdm3* allele were fully light (pigmentation score 0-1 in the A6 segment), at all rearing temperatures (Fig. 3A). F1 heterozygous females showed dark pigmentation, identical to the dark homozygotes, at all temperatures in both reciprocal crosses (Fig. 3B). Thus, both *D. burlai* and *D. rufa* display little or no temperature plasticity, and the dark *pdm3* allele is fully dominant at all temperatures within these species’ survival range.

**Figure 3.**
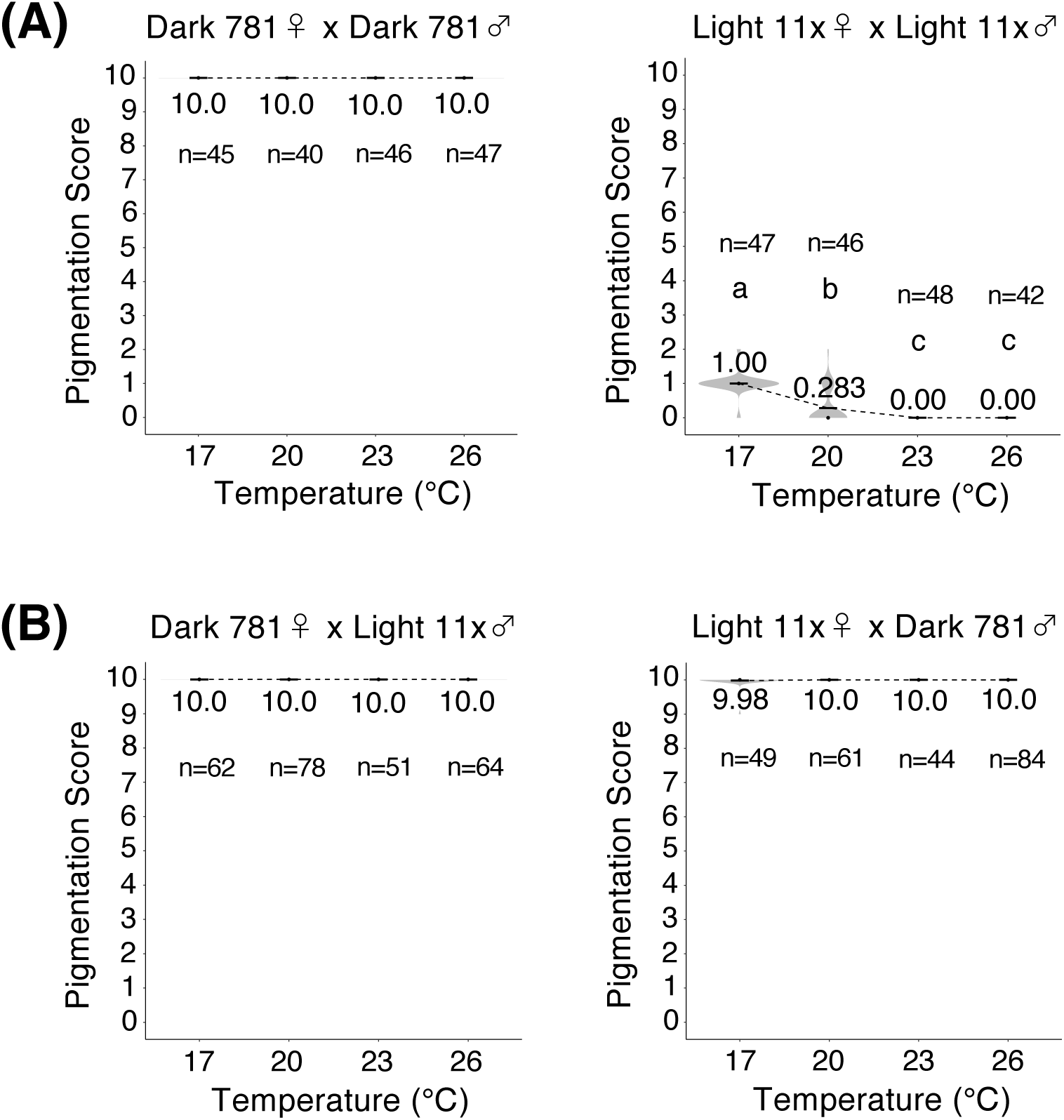
Female A6 pigmentation in *D. burlai* as a function of rearing temperature. Labeling as in Figure 2. (A) The homozygous parental strains Dark 781 (left) and Light 11x (right). A slight dependence on temperature is observed in the Light homozygotes (one-way ANOVA, df = 3; *F* = 120.7, *p* < 10^-10^), but not in the Dark homozygotes (*F* = 0.985). (B) No temperature plasticity is observed in the F1 females from reciprocal crosses between Dark and Light parents (*F* = 1.038 for the left cross, and *F* = 1.29 for the right cross). Black bars and black dots indicate mean and median values, respectively; the numbers indicate mean values. Different letters indicate significant differences (*p* < 0.05, Tukey’s HSD test).

### Temperature-dependent dominance in *D.* cf. *bocqueti* and *D. jambulina*

*D.* cf. *bocqueti* females homozygous for the dark *pdm3* allele were fully dark (pigmentation score ∼10 in the A6 segment), while those homozygous for the light *pdm3* allele were fully light (pigmentation score ∼0 in the A6 segment), at all temperatures (Fig. 4A). However, F1 heterozygous females showed strong temperature plasticity in all crosses (Fig. 4B-E; Supplement Table S6). At low rearing temperature, the heterozygotes had the same pigmentation as the dark homozygote, whereas at high temperature the heterozygotes had pigmentation scores similar to the light homozygote; at intermediate rearing temperatures, intermediate pigmentation was observed (Fig. 4B-E). In other words, the dark *pdm3* allele in *D.* cf. *bocqueti* is dominant at low temperature, the light allele is dominant at high temperature, and the two alleles are co-dominant at intermediate temperatures.

**Figure 4.**
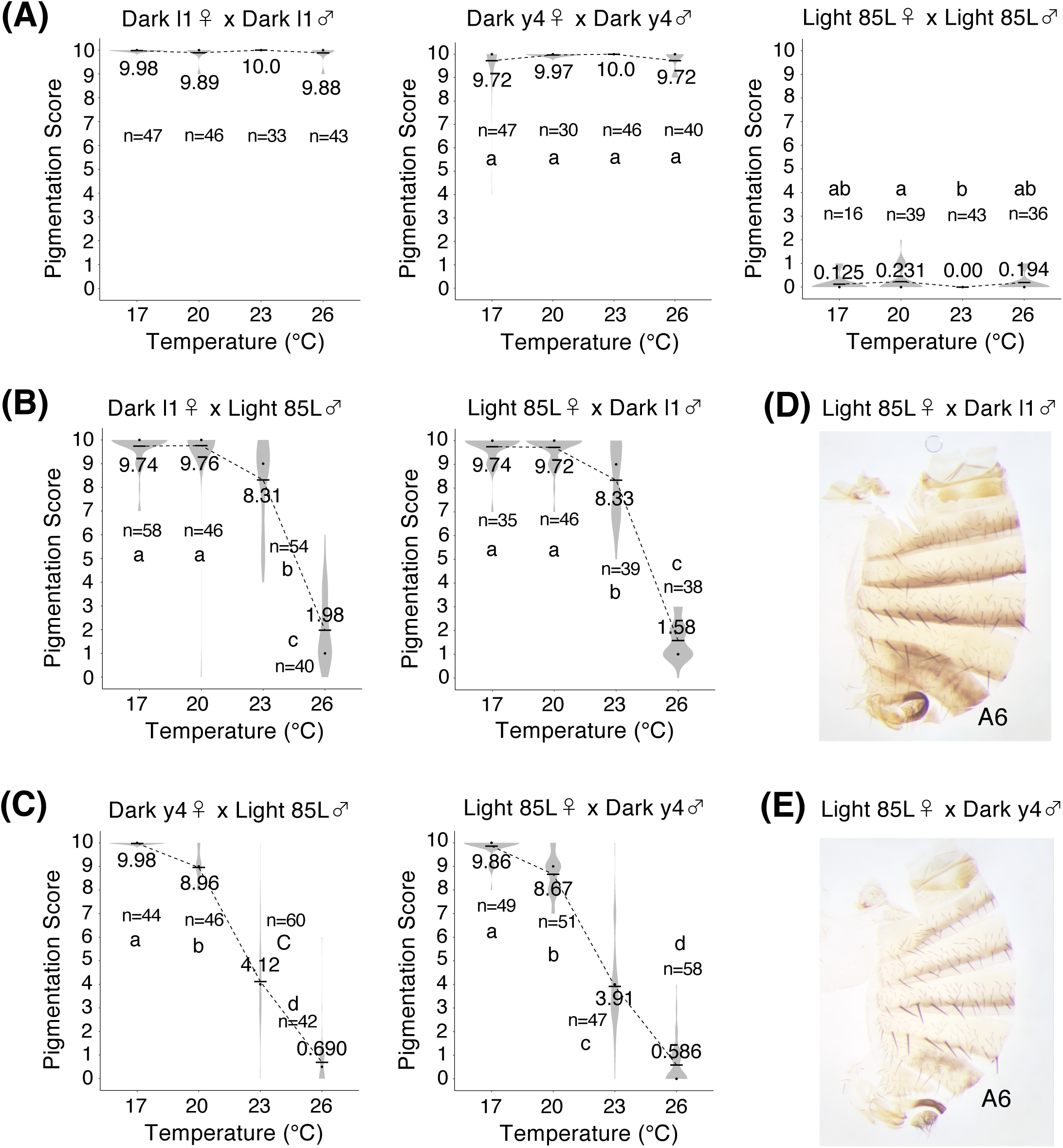
Female A6 pigmentation in *D.* cf. *bocqueti* as a function of rearing temperature. Labeling as in Figure 2. (A) The homozygous parental strains Dark l1 (left), Dark y4 (middle), and Light 85L (right). A slight dependence on temperature (one-way ANOVA, df = 3) is observed in the Light 85L (*F* = 3.374, *p* = 0.021) and Dark y4 homozygotes (*F* = 2.968, *p* = 0.034), but not in the Dark l1 homozygotes (*F* = 2.401). (B) F1 females from both reciprocal crosses between the Dark l1 and Light 85L parents show a strong temperature dependence (*F* = 300.6 for the left cross and *F* = 636.7 for the right, *p* < 10^-10^ for both). (C) F1 females from both reciprocal crosses between the Dark y4 and Light 85L parents show a strong temperature dependence (*F* = 517.0 for the left cross and *F* = 748.0 for the right, *p* < 10^-10^ for both). Note that the plasticity curves differ between the two heterozygous genotypes, depending on the Dark parent, but no differences are observed between reciprocal crosses (Supplement Table S6). (D) The abdominal pigmentation of an F1 female from the cross between Light 85L females and Dark l1 males, reared at 23 °C. (E) The abdominal pigmentation of an F1 female from the cross between Light 85L females and Dark y4 males, reared at 23 °C.

Similarly, strong temperature plasticity was observed in *D. jambulina* (Fig. 5; Supplement Table S7). *D. jambulina* females homozygous for the dark *pdm3* allele were fully dark (pigmentation score ∼10 in the A6 segment), while those homozygous for the light *pdm3* allele were fully light (pigmentation score ∼0 in the A6 segment), at all temperatures (Fig. 5A). At high rearing temperature, F1 heterozygous females had pigmentation similar to the light homozygote, whereas intermediate pigmentation was observed at low temperature (Fig. 5B-E). Thus, in this species, the light *pdm3* allele is dominant at high temperature, and the two alleles are co-dominant at lower temperatures.

**Figure 5.**
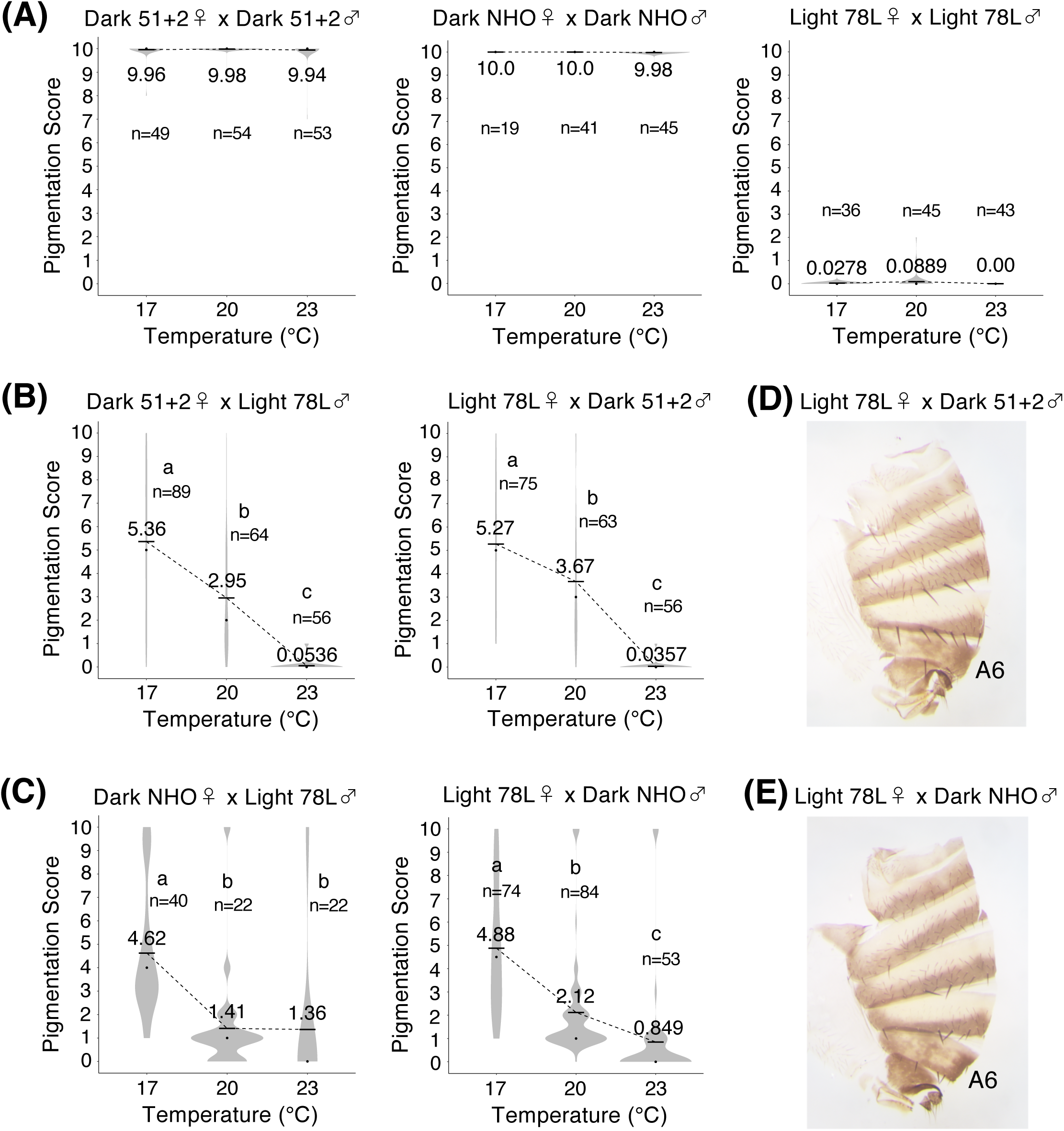
Female A6 pigmentation in *D. jambulina* as a function of rearing temperature. Labeling as in Figure 2. (A) The homozygous parental strains Dark 51+2 (left), Dark NHO (middle), and Light 78L (right). No significant differences between temperatures are observed in any of the three strains (one-way ANOVA, df = 2, *F* = 0.22 for the left cross, *F* = 0.66 for the cross in the middle, and *F* = 1.66 for the right cross). (B) F1 females from both reciprocal crosses between the Dark 51+2 and Light 78L parents show a strong temperature dependence (*F* = 84.38 for the left cross and *F* = 78.76 for the right, *p* < 10^-10^ for both). (C) F1 females from both reciprocal crosses between the Dark NHO and Light 78L parents show a strong temperature dependence (*F* = 13.66, *p* < 10^-5^ for the left cross and *F* = 47.38, *p* < 10^-10^ for the right). Note that the plasticity curves differ between the two heterozygous genotypes, depending on the Dark parent, but no differences are observed between reciprocal crosses (Supplement Table S7). (D) The abdominal pigmentation of an F1 female from the cross between Light 78L females and Dark 51+2 males, reared at 17 °C. (E) The abdominal pigmentation of an F1 female from the cross between Light 78L females and Dark NHO males, reared at 17 °C.

### Plasticity/dominance curves differ between species and genotypes

While allelic dominance is temperature-dependent in both *D.* cf. *bocqueti* and *D. jambulina*, with higher temperatures leading to a higher dominance of the light *pdm3* allele, the shape of plasticity curves is clearly different between these species. At low rearing temperatures, the dark *pdm3* allele is fully dominant in *D.* cf. *bocqueti,* but shows only partial dominance in *D. jambulina* (Fig. 4 vs Fig. 5). The dependence of the plasticity/dominance curves on the genotype is also observed within each of these species. In *D.* cf. *bocqueti,* crosses using the same light parental strain, but different dark parents, result in different phenotypes of the F1 heterozygotes; these differences are especially pronounced at intermediate rearing temperatures (Fig. 4B vs Fig. 4C). Two-way ANOVA shows that the effects of genotype are highly significant in both directions of the cross, and the genotype*temperature interaction is significant in all comparisons (Table 3).

**Table 3.**
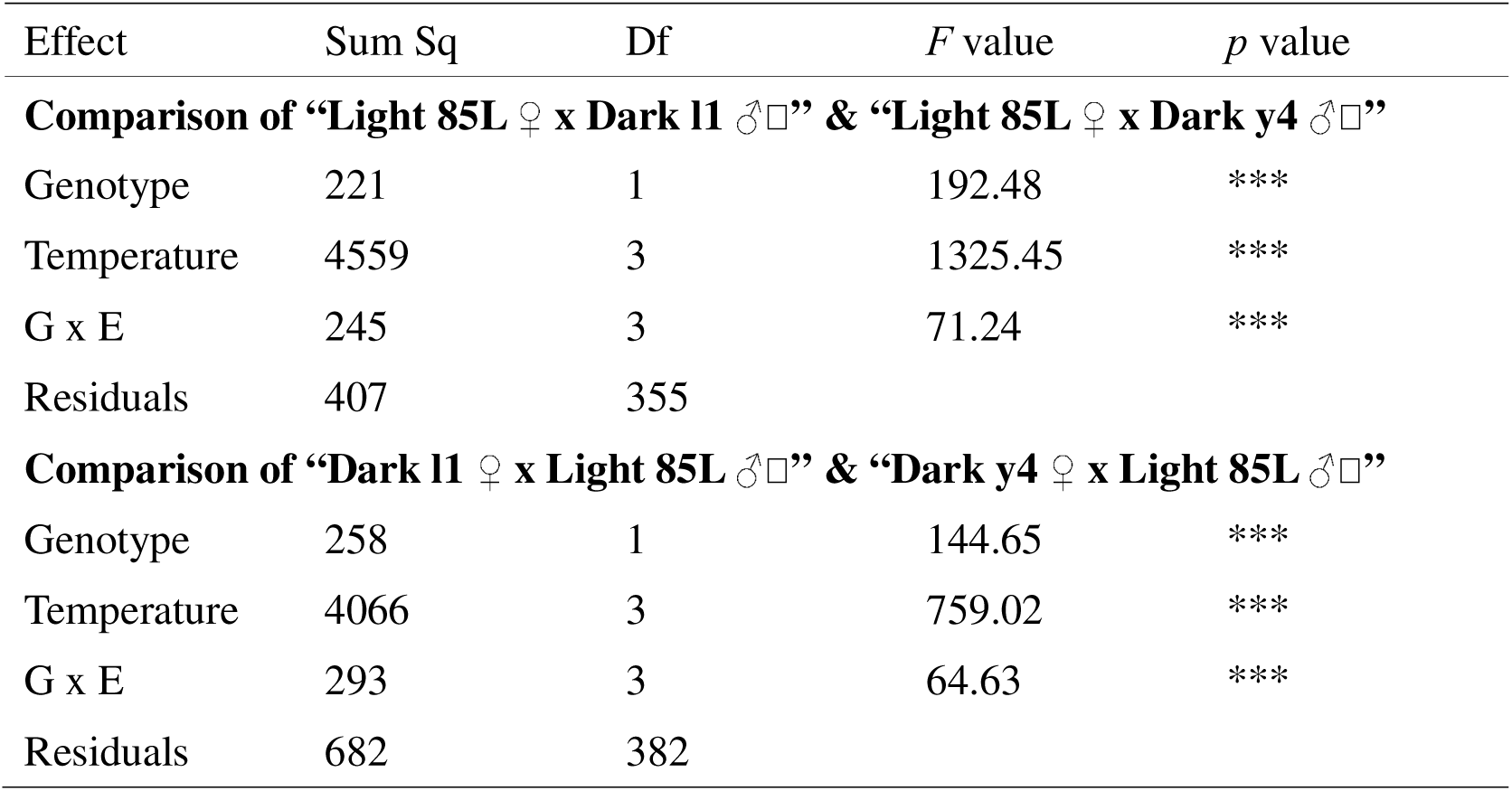
The results of two-way ANOVA on A6 pigmentation scores in heterozygous females of *D.* cf. *bocqueti*. Four “Genotypes” were used in the analyses: F1 females from the crosses between Light 85L females and Dark l1 males, between Light 85L females and Dark y4 males, between Dark l1 females and Light 85L males, and between Dark y4 females and Light 85L males. Two “Genotypes” (same direction of the cross, but using different Dark parents) were compared in each contrast. “G x E” indicates an interaction between “Genotype” and “Temperature”. Sum Sq: sum of squares, Df: degrees of freedom, ***: *p* < 10^-10^.

| Effect | Sum Sq | Df | F value | p value |
| --- | --- | --- | --- | --- |
| **Comparison of “Light 85L ♀ x Dark l1 ♂□” & “Light 85L ♀ x Dark y4 ♂□”** |  |  |  |  |
| Genotype | 221 | 1 | 192.48 | *** |
| Temperature | 4559 | 3 | 1325.45 | *** |
| G x E | 245 | 3 | 71.24 | *** |
| Residuals | 407 | 355 |  |  |
| **Comparison of “Dark l1 ♀ x Light 85L ♂□” & “Dark y4 ♀ x Light 85L ♂□”** |  |  |  |  |
| Genotype | 258 | 1 | 144.65 | *** |
| Temperature | 4066 | 3 | 759.02 | *** |
| G x E | 293 | 3 | 64.63 | *** |
| Residuals | 682 | 382 |  |  |

Similarly, in *D. jambulina*, crosses using the same light parental strain, but different dark parents, result in somewhat different F1 phenotypes, especially at intermediate temperatures (Fig. 5B vs Fig. 5C). In the ANOVA analysis, there is a significant effect of genotype in crosses using Light parental females, but not in the reciprocal crosses using Dark parental females. The genotype*temperature interaction is significant in all comparisons (Table 4).

**Table 4.** The results of two-way ANOVA on A6 pigmentation scores in heterozygous females of *D. jambulina*. Four “Genotypes” were used in the analyses: F1 females from the crosses between Light 78L females and Dark 51+2 males, between Light 78L females and Dark NHO males, between Dark 51+2 females and Light 78L males, and between Dark NHO females and Light 78L males. Two “Genotypes” (same direction of the cross, but using different Dark parents) were compared in each contrast. “G x E” indicates an interaction between “Genotype” and “Temperature”. Sum Sq: sum of squares, Df: degrees of freedom, ***: *p* <10^-10^, **: *p* < 10^-2^, *: *p* < 0.05, NS: not significant.

| Effect | Sum Sq | Df | *F* value | *p* value |
| --- | --- | --- | --- | --- |
| **Comparison of “Light 78L ♀ x Dark 51+2 ♂□” & “Light 78L ♀ x Dark NHO ♂□”** |  |  |  |  |
| Genotype | 23.0 | 1 | 3.96 | * |
| Temperature | 1367.0 | 2 | 117.85 | *** |
| G x E | 86.9 | 2 | 7.49 | ** |
| Residuals | 2314.1 | 399 |  |  |
| **Comparison of “Dark 51+2 ♀ x Light 78L ♂□” & “Dark NHO ♀ x Light 78L ♂□”** |  |  |  |  |
| Genotype | 10.3 | 1 | 1.61 | NS |
| Temperature | 1116.6 | 2 | 87.08 | *** |
| G x E | 70.7 | 2 | 5.51 | ** |
| Residuals | 1840.0 | 287 |  |  |

To further explore the dependence of dominance and plasticity on the genotype, we crossed the light strains of the plastic *D.* cf. *bocqueti* and the non-plastic *D. burlai* to the closely related species *D. chauvacae*, which, to our knowledge, is monomorphically Dark (Fig. 6A). These crosses could only be performed in one direction due to the reproductive isolation between these species. *D. chauvacae* females were fully dark at all temperatures (pigmentation score 10 in the A6 segment; Fig. 6B). Unlike the fully dark and non-plastic heterozygotes of *D. burlai* (Fig. 3B), F1 hybrid females between *D. chauvacae* and *D. burlai* Light 11x showed a weak temperature plasticity and were slightly lighter at high rearing temperatures (Fig. 6C; Fig. S3). F1 hybrid females between *D. chauvacae* and *D.* cf. *bocqueti* Light 85L showed a high degree of plasticity, with a dark phenotype at low temperature, a light phenotype at high temperature, and intermediate pigmentation at intermediate temperatures (Fig. 6D, E). The shape of the plasticity curve was within the range of those observed in *D.* cf. *bocqueti* heterozygotes (Fig. 4B, C). Overall, much greater plasticity is observed in hybrids between *D.* cf. *bocqueti* and *D. chauvacae* than between *D. burlai* and *D. chauvacae*; this is consistent with the temperature-independent dominance of the dark *pdm3* allele in *D. burlai* and temperature-dependent dominance in *D.* cf. *bocqueti*.

**Figure 6.**
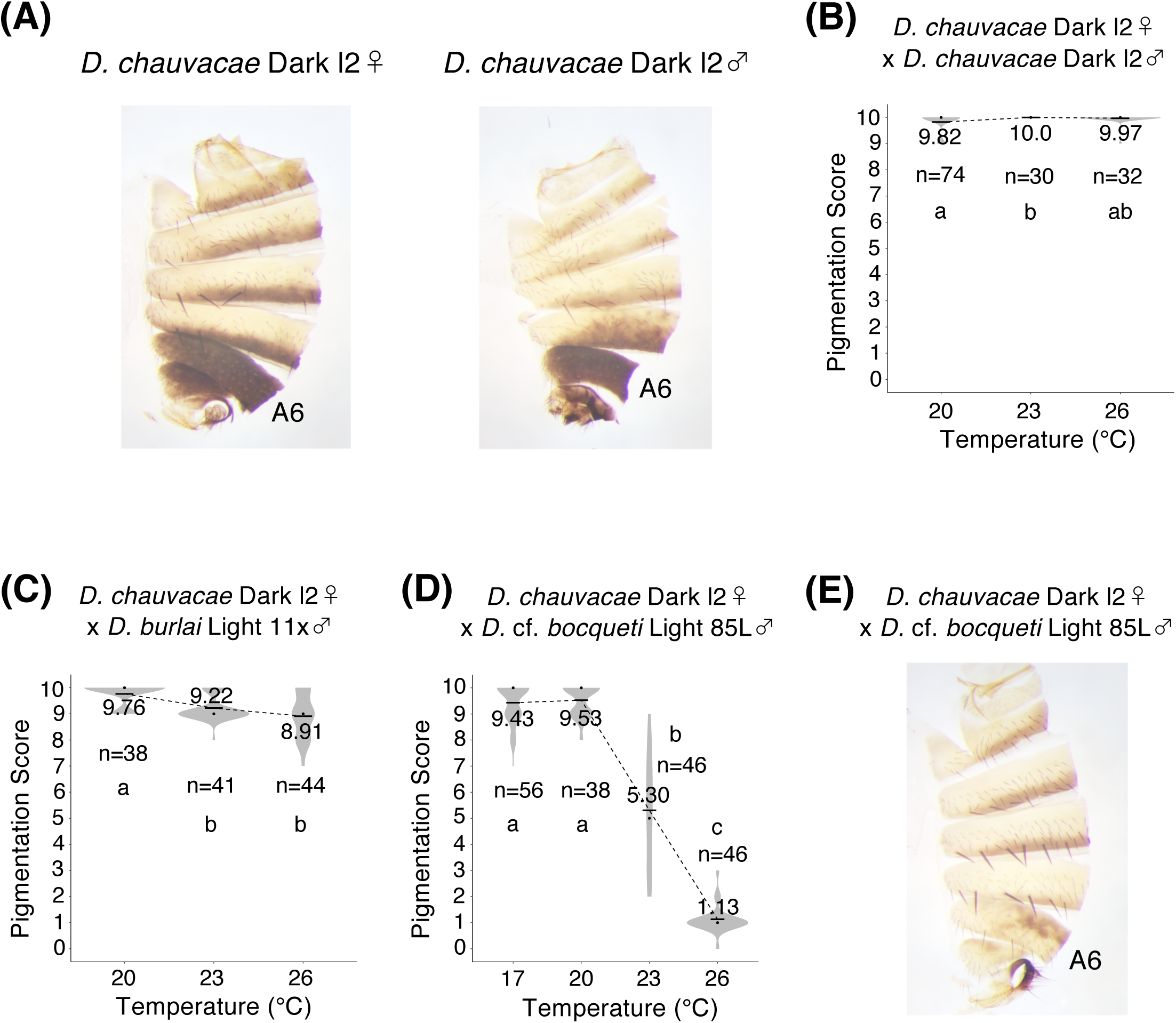
Female A6 pigmentation in the interspecific crosses between *D. chauvacae*, *D. burlai*, and *D.* cf. *bocqueti,* as a function of rearing temperature. Labeling as in Figure 2. (A) Female and male abdominal pigmentation in the *D. chauvacae* Dark l2 strain. (B) *D. chauvacae* Dark l2 homozygotes show only slight temperature plasticity (one-way ANOVA, df = 2; *F* = 4.973, *p* = 0.0083). (C) F1 females from the cross between *D. chauvacae* Dark l2 females and *D. burlai* Light 11x males show somewhat greater temperature plasticity than either the *D. chauvacae* parent (Fig 6B) or the *D. burlai* heterozygotes (Fig 3B) (one-way ANOVA, df = 2; *F* = 19.03, *p* < 10^-7^). (D) F1 females from the cross between *D. chauvacae* Dark l2 females and *D.* cf. *bocqueti* Light 85L males show greater temperature plasticity than the *D. chauvacae* parent, and comparable to *D.* cf. *bocqueti* heterozygotes (Fig 4B, C) (one-way ANOVA, df = 3; *F* = 513.2, *p* < 10^-10^). (E) The abdominal pigmentation of an F1 female from the cross between *D. chauvacae* Dark l2 females and *D.* cf. *bocqueti* Light 85L males, reared at 23 °C.

### Dominance and plasticity differ between sexes and segments

The patterns of dominance and plasticity in the female A7 segment are very similar to the A6, with only slight deviations (Fig.7, Fig. S4-S8). The dark *pdm3* allele is fully dominant across all rearing temperatures in *D. rufa* and *D. burlai* (Fig. S4, S5), whereas *D.* cf. *bocqueti*, *D. jambulina*, and interspecific hybrids between *D. chauvacae* and *D.* cf. *bocqueti* or *D. burlai* show temperature-dependent dominance, with darker pigmentation at low temperature and lighter pigmentation at high temperature (Fig. S3, S6-S8). In *D.* cf. *bocqueti*, the differences in plasticity curves between genotypes that are seen in the A6 (Fig. 4) are also reproduced in the A7 (Fig. S6). However, there are also some differences between segments. For example, in the *Dark y4* / *Light 85L* genotype of *D.* cf. *bocqueti*, the greatest change in pigmentation occurs between 20°C and 23°C in the A6 (Fig. 4), but between 17°C and 20°C in the A7 (Fig. S6). Minor differences in plasticity curves between A6 and A7 are also observed in *D. jambulina* (Fig. 5 vs Fig. S7), but the overall trends of dominance and plasticity are similar between these segments. In interspecific hybrids between *D. chauvacae* and *D. burlai*, plasticity curve in the A7 was much steeper than that in the A6. At a higher temperature, almost light phenotype was observed in the A7, whereas the dark phenotype was observed in the A6 (Fig. 6 vs Fig. S8). However, the tendency of darker pigmentation at low temperature and lighter pigmentation at high temperature is shared between A6 and A7.

**Figure 7.**
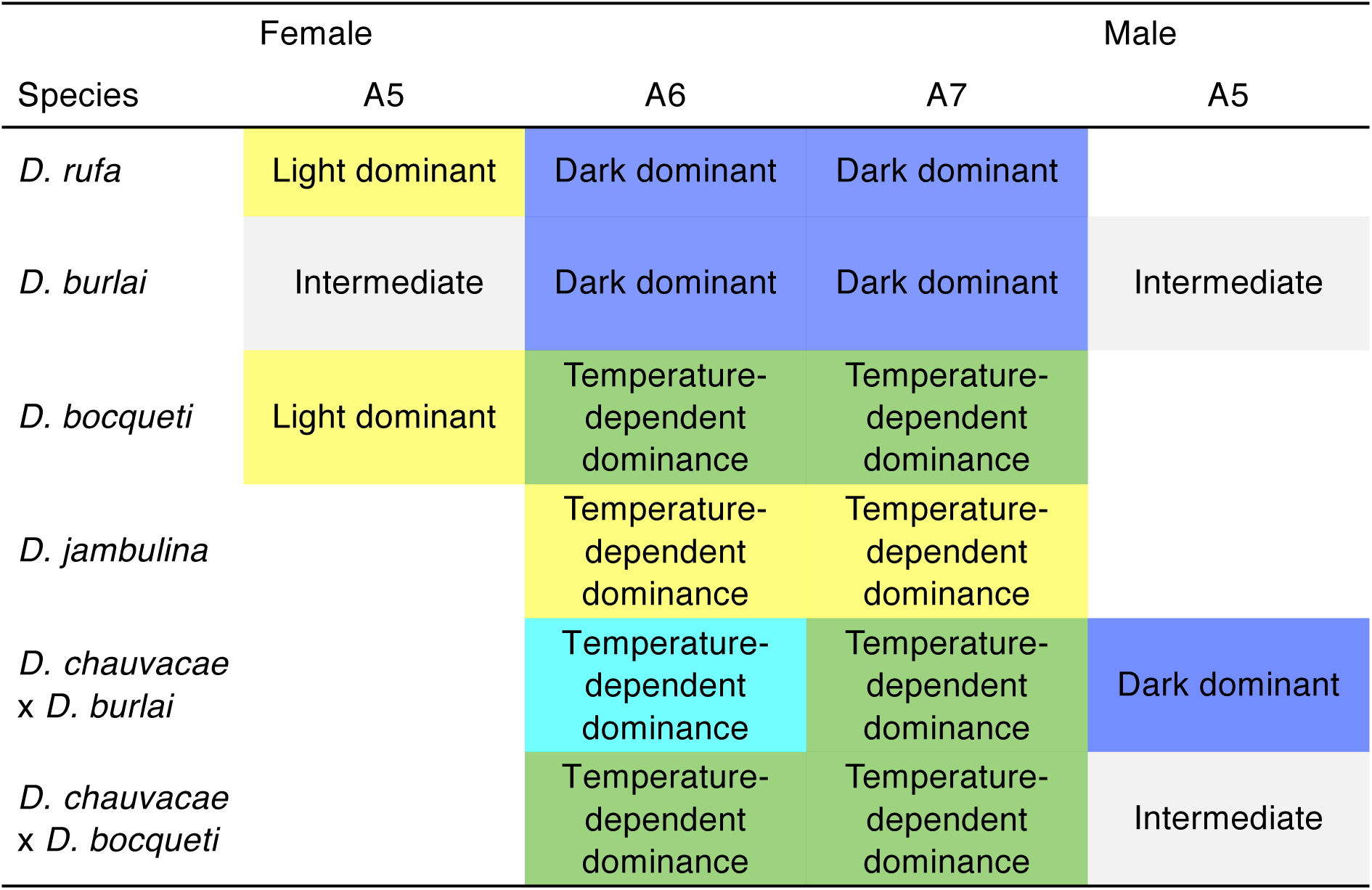
Dominance relationships between the Light and Dark *pdm3* alleles vary across species, segments, and sexes, with multiple examples of temperature-dependent dominance reversal. Abdominal segments 5-7 are shown for females, and abdominal segment 5 for males. All other segments have invariant pigmentation. Segments where the pigmentation of heterozygotes is similar to Light homozygotes across all temperatures are designated as “Light dominant” and highlighted in yellow. Segments where the pigmentation of heterozygotes is similar to Dark homozygotes across all temperatures are designated as “Dark dominant” and highlighted in blue. Segments where the dominance relationship between the Light and Dark alleles varies across temperatures are designated as “Temperature-dependent dominance”. Of these, cases where the Light allele is fully dominant at high temperature and the Dark allele is fully dominant at low temperature are highlighted in green. Cases where the Light allele is fully dominant at high temperature and intermediate dominance is observed at low temperature are highlighted in yellow. A case where the Dark allele is fully dominant at low temperature and incomplete dominance of the Dark allele is observed at high temperature is highlighted in cyan. Segments where the pigmentation of heterozygotes is intermediate between Dark and Light homozygotes across all temperatures are designated as “Intermediate” and highlighted in light gray. Segments where no difference is observed between Light and Dark homozygotes are left blank.

Unlike A6 and A7, the A5 segment in females shows little difference between the light and dark genotypes except in *D. burlai* (Fig. 1). Interspecific differences in the patterns of dominance and plasticity in the A5 are not as clear as in the A6. In *D.* cf. *bocqueti* and *D. rufa*, the phenotype of heterozygotes was similar to the light homozygotes across all rearing temperatures (Fig. 7, Fig. S9, S10), although the differences between light and dark homozygotes are generally slight. In *D. burlai*, the alleles responsible for the phenotypic difference in A5 are co-dominant, with heterozygotes somewhat closer to the light homozygotes, at all temperatures (Fig. 7, Fig. S11), while in *D. jambulina* all three genotypes have similar pigmentation (Fig. 7, Fig. S12). Light homozygotes of *D. burlai*, dark homozygotes of *D. chauvacae*, and their hybrids did not show obvious difference in phenotype, but in the interspecific crosses between *D. chauvacae* and *D.* cf. *bocqueti*, the pigmentation score was closer to the light *D.* cf. *bocqueti* parents (Fig. 7, Fig. S13). In contrast to the A6 segment, which shows a monotonous decrease in pigmentation with increasing temperature, plasticity is less predictable in the A5: while *D. jambulina* does show a similar trend (Fig. S12), *D.* cf. *bocqueti* has the lightest pigmentation at intermediate temperatures (Fig. S10), *D. burlai* has the darkest pigmentation at intermediate temperatures (Fig. S11), and *D. rufa* shows very little temperature plasticity (Fig. S9), as do the interspecific hybrids with *D. chauvacae* (Fig. S13).

In males, the A6 segment is completely dark at all temperatures, regardless of species and genotype (Fig. 1, Fig. S14-S18). The A5 segment in males is light at all temperatures in *D. rufa* and *D. jambulina*, and dark at all temperatures in *D.* cf. *bocqueti*, with a slight trend toward lighter pigmentation at higher temperatures in the latter two species (Fig. S19-21). In *D. burlai*, however, there is a clear difference in the A5 between light and dark homozygotes; heterozygotes show intermediate pigmentation, though closer to the dark homozygotes (Fig. 7, Fig. S22). In this species, the trend toward lighter pigmentation at higher temperatures is more pronounced in heterozygotes than in either light or dark homozygotes, indicating a potential temperature-dependent dominance reversal, although this effect is weak (Fig. 7, Fig. S22). Finally, *D. chauvacae* shows an unusual trend of darker pigmentation at higher temperatures in the male A5, which is also observed in hybrids between *D. chauvacae* and *D.* cf. *bocqueti* (Fig. S23).

## Discussion

The Mendelian, female-limited color dimorphism widespread in the *montium* species subgroup is controlled by the *pdm3* transcription factor in at least six different species (Yassin et al., 2016, and this report). Our results show that the key difference between the species that were described as “dark-dominant” and “light-dominant” is related to phenotypic plasticity. In *D. kikkawai* (see David et al. 2025), *D. rufa, D. burlai*, and probably most other female-dimorphic species, the Dark *pdm3* allele is dominant at all temperatures, whereas in both known “light-dominant” species, *D. jambulina* and *D.* cf. *bocqueti*, dominance is temperature-dependent, with the Light allele becoming increasingly dominant at higher temperatures. Thus, the evolution of dominance relationships between *pdm3* alleles is intimately connected to the evolution of thermal plasticity. Below, we consider potential mechanisms that could be responsible for this connection.

### A threshold model linking the evolution of dominance and phenotypic plasticity

The molecular mechanisms that link dominance and plasticity remain to be elucidated. Given that dominance is an inherently non-linear relationship, this link is likely to lie in the threshold behavior characteristic of many plastic traits (Nijhout, 2003; Suzuki & Nijhout, 2006; Bhardwaj et al., 2018; Grieshop et al., 2024). Based on this hypothesis, we propose that interspecific differences in the dominance relationships between *pdm3* alleles reflect different threshold responses of the pigment synthesis pathway to gradually changing levels of *pdm3* expression (Fig. 8). This model makes three assumptions. First, we propose that Light alleles drive higher levels of *pdm3* expression compared to Dark alleles. Second, we propose that the expression level of *pdm3* increases with temperature (Fig. 8A). Since Pdm3 is a repressor of pigmentation (Yassin et al., 2016), higher temperatures lead to lighter pigmentation. Third, we propose that melanin synthesis is repressed when *pdm3* level rises above a certain threshold, and enabled below that threshold; intermediate pigmentation is produced within a narrow intermediate range of *pdm3* levels (Fig. 8A).

**Figure 8.**
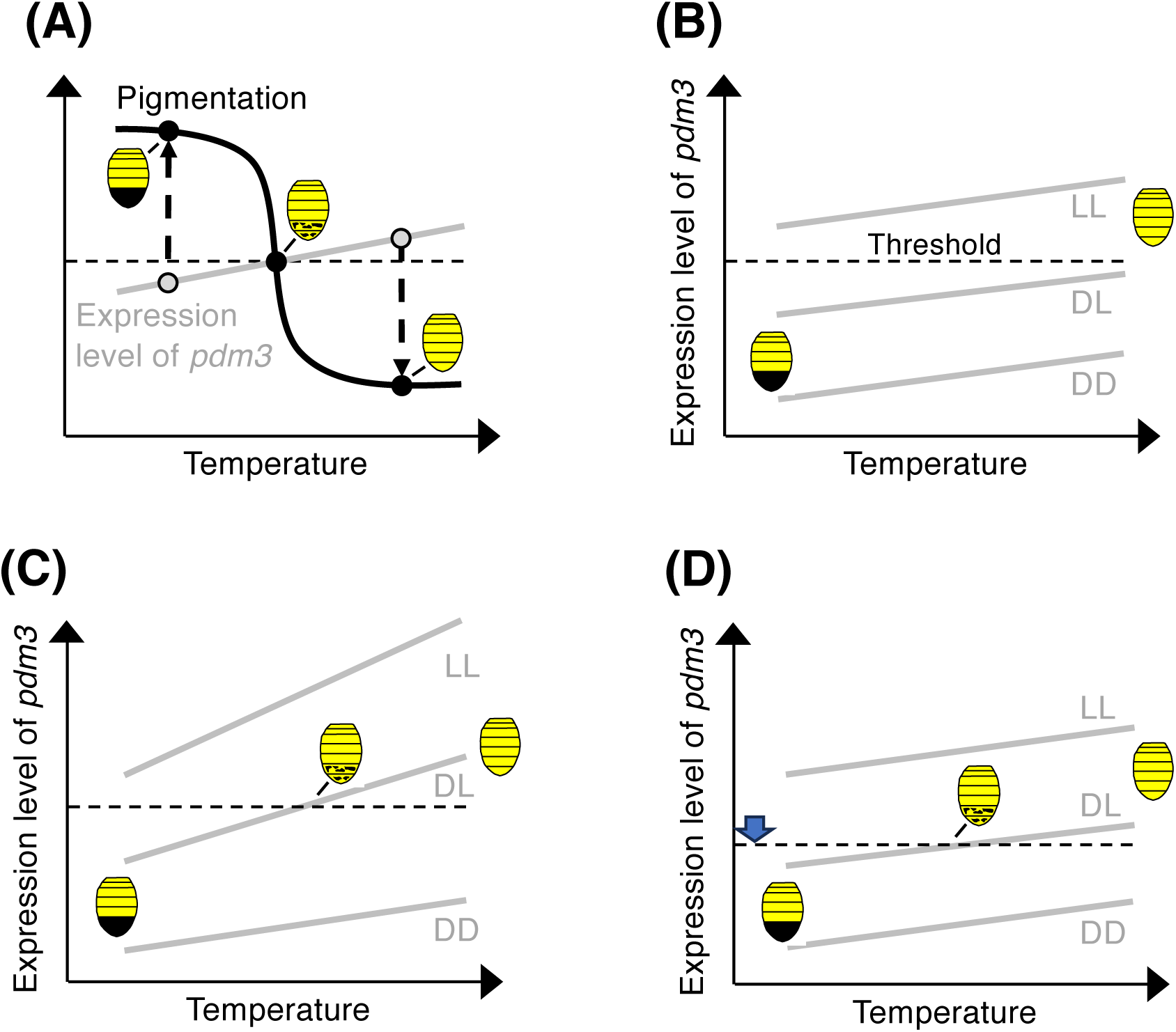
A putative model of the relationship between environmental plasticity and allelic dominance. (A) Hypothesized relationship between temperature, *pdm3* expression (grey curve), and pigmentation (black curve). *pdm3* is a repressor of dark pigmentation (Yassin et al. 2016). We propose that *pdm3* expression increases at higher temperatures, leading to lighter pigmentation in a threshold-dependent manner. When *pdm3* expression is above the threshold (horizontal dashed line), the result is a Light phenotype (downward arrow); when *pdm3* is expressed below the threshold, the result is a Dark phenotype (upward arrow); intermediate temperatures may lead to intermediate *pdm3* expression and intermediate pigmentation. (B) Hypothesized relationship between temperature and *pdm3* expression in species where the dark *pdm3* allele is fully dominant. Light *pdm3* alleles cause higher *pdm3* expression than Dark alleles; all *pdm3* alleles respond to temperature in a similar manner; and the threshold is such that *pdm3* expression is below the threshold in DD homozygotes and DL heterozygotes, and above the threshold in LL homozygotes, at all temperatures. (C) Hypothesized relationship between temperature and *pdm3* expression in species with temperature-dependent dominance. In this model, the threshold is the same, but the Light *pdm3* allele has a steeper temperature response than the Dark allele; as a result, the DL heterozygotes cross the pigmentation threshold (dashed horizontal line) at intermediate temperatures. (D) An alternative hypothesis for the relationship between plasticity and dominance. In this model, all *pdm3* alleles respond to temperature in a similar manner, but the pigmentation threshold is lower than in species with full dominance (downward arrow). As a result, DL heterozygotes cross the pigmentation threshold at intermediate temperatures. Models (C) and (D) are not mutually exclusive; different mechanisms may operate in different species, or both mechanisms may operate in the same species.

Collectively, these three assumptions are sufficient to explain the full range of phenotypes and dominance relationships observed in the *montium* subgroup. In species such as *D. rufa* and *D. burlai*, where the Dark *pdm3* allele is fully dominant, we hypothesize that *pdm3* expression is above the threshold in Light homozygotes and below the threshold in Dark homozygotes and heterozygotes at all temperatures. In the simplest case, the Light and Dark alleles have similar responses to temperature changes (Fig. 8B), but this assumption is not strictly required. In species with dominance reversal or temperature-dependent dominance, such as *D.* cf. *bocqueti* and *D. jambulina*, we propose that *pdm3* expression levels in heterozygotes cross the phenotypic threshold. This can occur for two, not mutually exclusive reasons. One possibility is that the Light *pdm3* allele has a steeper response to temperature compared to the Dark allele (Fig. 8C). Alternatively, the Dark and Light alleles may respond to temperature in a similar manner, both to each other and across species, but the phenotypic threshold may be lower in species with dominance reversal compared to species with constant dominance (Fig. 8D). These two alternatives may apply to different species, or both mechanisms could operate in the same species.

This descriptive model raises several questions about its mechanistic basis, namely: (1) why does *pdm3* expression increase with temperature? (2) why does a linear change in *pdm3* expression lead to a threshold-like phenotypic transition? (3) what explains expression differences between the Light and Dark *pdm3* alleles? (4) why does the phenotypic threshold differ between species? (5) why does allelic dominance differ between segments or sexes? We do not know the answers to any of these questions, but outline some potential explanations and avenues for future investigation below.

### Why could *pdm3* expression increase with temperature?

Across the entire genome, many genes show either increased or decreased transcript abundance at higher temperatures (Chen et al., 2015; Gómez-Orte et al., 2017; Voigt & Kost, 2021; Voigt & Froschauer, 2023; Pathak et al., 2025). The reasons for this are not fully understood, and are likely to be different for different loci. Some genes may be regulated by systemic hormones. In many cases where developmental traits show environmental plasticity, the titers of hormones such as ecdysone or juvenile hormone are condition-dependent, and changes in hormone titers induce phenotypic changes (Suzuki & Nijhout, 2006; Bhardwaj et al., 2020; Yoon et al., 2023). For example, in the polyphenic butterfly *Bicyclus anynana*, temperature influences the timing and intensity of ecdysone pulses, which in turn trigger transcriptional changes across hundreds of loci (Tian & Monteiro, 2022). Thus, one possibility is that *pdm3* is regulated in part by a temperature-sensitive hormonal pathway.

Alternatively, temperature-dependent *pdm3* expression could reflect changes in chromatin state. Temperature changes can influence chromatin compaction, making regulatory sequences more or less accessible to transcription factors (TFs) (Kumar & Wigge, 2010; Cortijo et al., 2017). In *Drosophila* cells, temperature shifts increase or decrease chromatin accessibility at hundreds of enhancers and promoters, correlating with up- or downregulation of nearby genes, and reporter assays confirm that temperature-responsive DNA elements can directly confer temperature-dependent transcription (Bai et al., 2021). There is a well-documented pattern where many genes regulated by Polycomb-Group (PcG) chromatin remodeling factors show higher expression at lower temperatures, along with correlated changes in histone modifications (Fauvarque & Dura, 1993; Bantignies et al., 2003; Voigt & Froschauer, 2023). However, some PcG targets show the opposite change (Voigt & Froschauer, 2023). In particular, *bric à brac* (*bab*), another TF with a key role in regulating abdominal pigmentation in *Drosophila*, shows higher expression at higher temperatures despite being a PcG target (De Castro et al., 2018). Since *bab*, like *pdm3*, represses melanin synthesis (Kopp et al., 2000), increased *bab* expression is part of the mechanism leading to lighter pigmentation at high temperatures (Gibert et al., 2007; De Castro et al., 2018). *pdm3* itself does not appear to be a direct PcG target; such genes are more likely to increase expression with temperature, compared to PcG targets (Voigt & Froschauer, 2023). Finally, transcription is controlled by multi-level regulatory hierarchies that involve both activation and repression. If *pdm3* transcription is either activated by a TF such as *bab* that shows increased expression at higher temperature, or repressed by a typical PcG target gene that shows decreased expression at higher temperature, *pdm3* expression will increase with temperature, leading to the pattern hypothesized in our model (Fig. 8). The first step toward testing this model will be to test whether, in fact, *pdm3* expression responds to temperature in the manner we predict.

### Why could a linear change in *pdm3* level lead to a threshold phenotypic response?

If *pdm3* expression responds to temperature in a linear manner, then the threshold-like phenotypic transition must depend on the downstream target genes of *pdm3*. A sigmoidal transcriptional response to a linear change in the concentration of an upstream TF is modelled by the Hill equation (Hill, 1910; Bhaskaran et al., 2015; Bottani & Veitia, 2017), and can arise from multiple molecular mechanisms. A key contributor is cooperative TF binding: when multiple TF molecules bind synergistically, small increases in TF concentration at intermediate levels can sharply boost transcriptional output (Gregor et al., 2007; Frank, 2013). Threshold-like behaviors can emerge from the architecture of *cis*-regulatory elements (CREs) even without direct cooperativity: CREs that require a minimum number of TFs to bind simultaneously inherently produce nonlinear responses, as the probability of achieving an active state rises steeply with TF concentration (Sherman & Cohen, 2012). Chromatin accessibility can add a further layer of nonlinearity: TFs and cofactors must often remodel closed chromatin before productive binding can occur, and such chromatin transitions can behave as cooperative, switch-like events stabilized by histone modifications (Frank, 2013; Park et al., 2019). Higher-order cooperativity, involving not only TF-TF interactions but also recruitment of additional cofactors, amplifies the sharpness of the transcriptional response by stabilizing enhancer-promoter communication (Park et al., 2019). Finally, energy-dependent processes such as ATP-driven chromatin remodeling can further sharpen input-output relations by allowing irreversible transitions between regulatory states (Estrada et al. 2016). Any combination of these factors can steepen the sigmoid transcriptional curve, bringing it closer to a switch-like response.

The switch between black and yellow pigmentation in *Drosophila* requires coordinated changes in the expression of multiple genes involved in melanin synthesis. In particular, black pigmentation requires upregulation of the *yellow* (*y*) and *tan* (*t*) genes, concurrently with the downregulation of *ebony* (*e*) (Wittkopp et al., 2002; Ordway et al., 2014; Massey & Wittkopp, 2016; Gibert et al., 2017; Rebeiz & Williams, 2017; Liu et al., 2019). This suggests that *pdm3* may regulate these, and potentially other pigmentation genes, either directly or through an intermediate TF. Given the function of *pdm3* as a repressor of dark pigmentation, an obvious hypothesis is that it represses *y* and *t*, and upregulates *e*. Indeed, in *D. yakuba* and *D. santomea*, *pdm3* interacts epistatically with *y* in determining abdominal pigmentation, and may be acting in part by repressing that gene (Liu et al., 2019). In the simplest case, then, the threshold pigmentation phenotypes seen in the *montium* subgroup could be explained by the presence of multiple Pdm3 binding sites in the CREs of *pdm3* target genes, especially if their spacing facilitates cooperative binding of Pdm3. To test this hypothesis, it will be necessary to identify the direct downstream targets of *pdm3*.

### What could explain the differences between *pdm3* alleles and phenotypic thresholds?

In principle, evolution of dominance relationships between alleles could be due either to changes in the alleles themselves, or to the action of modifier genes (Fisher, 1928; Wright, 1934; Kacser & Burns, 1981; Gilchrist & Nijhout, 2001; Porter et al., 2017). These alternatives can be related to different features of the threshold model (Fig. 8). If the steepness of *pdm3* response to temperature differs between Light and Dark alleles and/or between species, the most likely explanation lies in the structure of these alleles. For example, the Light *pdm3* allele could have both higher expression overall and a steeper response to higher temperature (Fig. 8C) due to differences in the number of binding sites for upstream regulators between the Light and Dark alleles. If, however, the difference between species with constant dominance and those with dominance reversal is due primarily to the shifting threshold (Fig. 8D), these shifts are more likely to reflect the existence of second-site modifiers. These modifiers could be either the downstream targets of *pdm3* that evolve changes in the shape of their response to *pdm3* levels, or other TFs that regulate pigmentation genes in parallel with *pdm3* such that the binding of these TFs changes the response of the target genes to Pdm3.

Multiple instances have been described where different alleles of the same gene differ in their transcriptional response to temperature changes (Levine et al., 2011; Gibert et al., 2011; De Castro et al., 2018; Voigt & Froschauer, 2023), suggesting that this type of variation is quite common. At the same time, there is a strong indication that the differences between Light and Dark *pdm3* alleles are due to changes in their *cis*-regulatory elements. In *D. serrata*, another species of the *montium* subgroup, the Light and Dark alleles are distinguished by a non-coding structural variant in the first intron of *pdm3* (Yassin et al., 2016). This change results in an increased number of predicted binding sites for *Abdominal-B* (*Abd-B*) in the Dark *pdm3* allele. Abd-B is a HOX TF that controls the development of posterior abdominal segments, including the induction of dark pigmentation (Kopp et al., 2000; Jeong et al., 2006; Williams et al., 2008; Liu et al., 2019). Thus, *pdm3* could be a direct target of *Abd-B*, and the difference in the number of Abd-B binding sites in the CREs of *pdm3* could lead to differential regulation of Light vs Dark alleles in posterior abdominal segments, similar to the difference in Pdm3 expression observed between *D. yakuba* and *D. santomea* (Liu et al., 2019). The causative DNA sequences that distinguish Light and Dark *pdm3* alleles in other species remain to be identified, but it is likely that these differences are also regulatory in nature. Such differences could involve the gain and loss of binding sites for upstream TFs and chromatin remodeling factors or changes in the sensitivity to hormone signaling.

If interspecific differences in dominance relationships between Light and Dark *pdm3* alleles are due to second-site modifiers, one possibility is that the downstream target genes have different responses to *pdm3* in different species. Many examples of inter- and intraspecific differences in the activity of pigmentation gene CREs have been described (Jeong et al., 2006, 2008; Rebeiz et al. 2011; Johnson et al., 2015; Roeske et al., 2018; Méndez-González et al., 2023), and interactions between these CREs and species-specific *trans*-regulatory backgrounds play a key role in the evolution of color patterns (Ordway et al., 2014; Camino et al., 2015; Liu et al., 2019; Hughes et al., 2020). The non-additive interaction between *pdm3* and *y* in *D. yakuba* / *D. santomea* hybrids (Liu et al., 2019) suggests that changes in the CREs of *pdm3* target genes are a plausible mechanism for the evolution of dominance via changes in the threshold response to *pdm3* levels.

Alternatively, second-site modifiers of dominance could include other TFs that act in parallel with *pdm3* to regulate pigment synthesis genes. In *D. melanogaster*, temperature-dependent modulation of *bab* expression plays a key role in the thermal plasticity of abdominal pigmentation (De Castro et al., 2018). *bab* represses melanin synthesis genes such as *y* and *t*, in part by binding directly to the abdominal CRE of *y*, so that higher *bab* expression leads to lighter pigmentation (Gibert et al., 2016, 2017; Roeske et al., 2018; Hughes et al., 2020). Importantly, *bab* alleles differ in their response to temperature: an allele associated with lighter pigmentation shows a steeper increase in activity at higher temperatures, compared to an allele associated with darker pigmentation (De Castro et al., 2018). If *pdm3* and *bab* co-regulate the same target genes, and the response of *bab* alleles to temperature varies among *montium* subgroup species, the threshold between light and dark pigmentation, and thus the dominance relationship between *pdm3* alleles, could evolve via changes at the *bab* locus. A similar logic could potentially apply to any other TF or chromatin regulator acting in parallel with *pdm3*. For example, *trithorax* (*trx*), a histone methylase with a key role in chromatin remodeling, is expressed equally in both sexes but affects the activity of *tan* and *ebony* CREs in a sex-specific manner, suggesting that it shapes the response of these CREs to upstream TFs (Weinstein et al., 2023). Finally, a recent study in *D. melanogaster* has shown that genetic variation in the plasticity of pigmentation does not map to the same loci as variation in pigmentation itself (Lafuente et al 2024), suggesting that key molecular pathways remain to be identified. In summary, there is no shortage of hypotheses that could explain the shifting phenotypic threshold. Distinguishing between these hypotheses will require identifying the place of *pdm3* in the wider genetic network that controls abdominal pigmentation, as well as testing directly for the existence of second-site modifiers of dominance.

### Why do allelic dominance relationships differ between segments or sexes?

Intersegmental or intersexual differences in the dominance of *pdm3* alleles could stem either from the regulation of *pdm3* itself, or from the regulation of its downstream targets. The top-level regulators of pigmentation in posterior abdominal segments are *Abd-B*, which controls differences between segments, and the sex determination gene *doublesex* (*dsx*), which is responsible for the differences in pigmentation between males and females (Kopp et al., 2000). These genes act at multiple levels in the pigmentation pathway. *Abd-B* and *dsx* jointly regulate the expression of *bab*, leading to sex- and segment specific expression of that TF which in turn leads to sex- and segment specific pigmentation (Kopp et al., 2000; Williams at al., 2008; Rogers et al., 2013). If *pdm3* is also under the control of *Abd-B*, *pdm3* expression levels will differ between A5, A6, and A7, which, under the threshold model of dominance (Fig. 8), may lead to different dominance relationships between the Light and Dark *pdm3* alleles. However, *Abd-B* also controls the expression of *y* directly (Jeong et al., 2006), in addition to controlling it indirectly via *bab* (Roeske et al., 2018); other enzymes in the pigment synthesis pathway could also be regulated in a similar manner. Thus, another possibility is that *Abd-B, dsx,* and *bab* drive different expression levels of one or more *pdm3* target genes in different segments and in males vs females, which in turn shifts the threshold response to *pdm3* expression, leading to changes in dominance (Fig. 8D). These models are not mutually exclusive, and both could contribute to the differences in dominance relationships between segments and sexes.

## Conclusion

Interspecific differences in dominance relationships between *pdm3* alleles in the *D. montium* species subgroup point to the existence of a mechanistic connection between the evolution of dominance and phenotypic plasticity. However, the molecular nature of this connection is unknown. Although the threshold model offers a potential explanation for the difference between stable and condition-dependent dominance (Fig. 8), this model is purely descriptive and raises as many questions as it answers. Future work will be needed to understand the mechanism of the threshold response, the functional differences between *pdm3* alleles, and the relative roles of *pdm3* alleles and second-site modifiers in the evolution of dominance.

## Data availability

Raw data of phenotypic analyses and R scripts are available via Figshare: https://doi.org/10.6084/m9.figshare.28941818. Sequencing data have been deposited at DDBJ under accession numbers LC874378 - LC874448 (71 samples).

## Author contribution

Y.F. and A.K. designed the research; Y.F., A.P.G., J.L., and A.C. collected data; Y.F. analyzed data; M.W. and S.R. collected the strains used in the research; Y.F. and A.K. wrote the paper with feedback from M.W. and S.R.

## Acknowledgments

We thank Jean R. David, Brandon Cooper, Daniel Matute, and the Ehime and Cornell *Drosophila* species stock centers for providing *Drosophila* strains; Olga Barmina for sample collection; David Begun, Susan Lott, and Joanna Chiu for the use of their incubators; Benjamin Hopkins and Amir Yassin for advice and comments on the manuscript.

