## Supplemental Figures for "Evolution of dominance in a Mendelian trait is linked to the evolution of environmental plasticity"

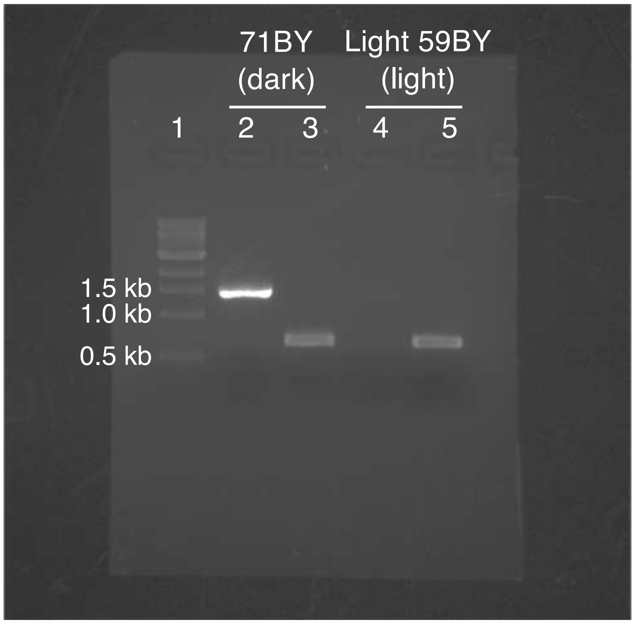


Figure. S1

The result of PCR on dark and light females of *D. rufa*. The regions homologous to the Dark and Light *pdm3* alleles of *D. serrata* were amplified. The digits from 1 to 5 indicate the numbers of lanes in the agarose gel. The sample in each lane is as follows. Lane 1: 1 kb DNA ladder from New England Biolabs. Lane 2: The PCR product amplified from the genomic DNA of a dark female from *71BY* strain with "Dark primer set" (forward: CCATACCATACAAGCGCCATCTAG, reverse: TTATCATGCATCAATGCGACAGCAAC). Lane 3: The PCR product amplified from the genomic DNA of a dark female from *71BY* strain with "Light primer set" (forward: CCATACCATACAAGCGCCATCTAG, reverse: AGCGACACAGATAACCAGATTTCGA). Lane 4: The PCR product amplified from the genomic DNA of a light female from *Light 59BY* strain with "Dark primer set". Lane 5: The PCR product amplified from the genomic DNA of a light female from *Light 59BY* strain with "Light primer set".

**(A)**

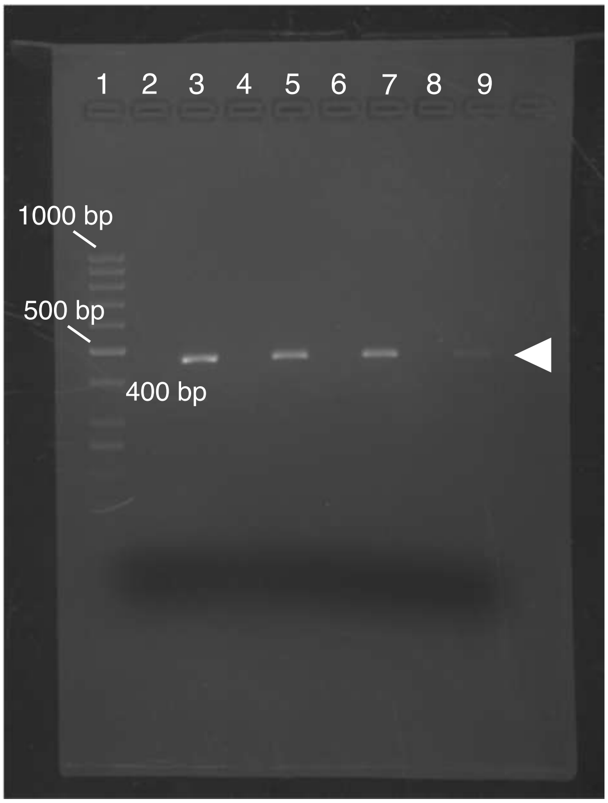


**(B)**

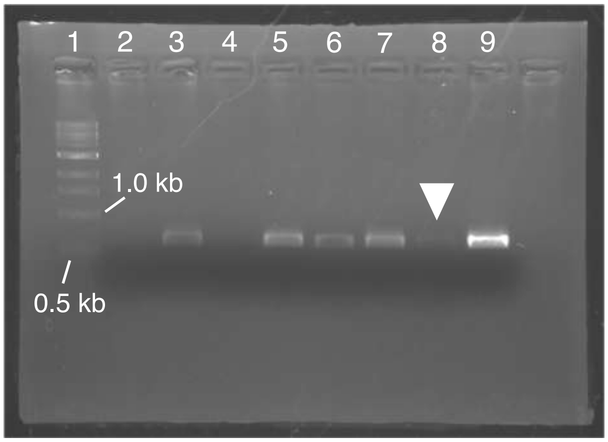

Figure. S2

The result of PCR on dark and light females of *D. jambulina*. (A) The 456 bp region containing sequences specific to the Light *pdm3* allele was amplified (the white arrowhead). The digits from 1 to 9 indicate the numbers of lanes in the agarose gel. The sample in each lane is as follows. Lane 1: GeneRuler 50bp DNA Ladder from Thermo Fisher Scientific. Lane 2: The PCR product amplified from the genomic DNA of a dark female from *Dark NHO* strain (a Dark homozygous strain, already genotyped with amplicon sequencing). Lane 3: The PCR product amplified from the genomic DNA of a light female from *58L* strain (a Light homozygous strain, already genotyped with amplicon sequencing). Lane 4: The PCR product amplified from the genomic DNA of a dark female from *60L* strain. Lane 5: The PCR product amplified from the genomic DNA of a light female from *Light 78L* strain. Lane 6: The PCR product amplified from the genomic DNA of a dark female from *103D* strain. Lane 7: The PCR product amplified from the genomic DNA of a light female from *71L* strain. Lane 8: The PCR product amplified from the genomic DNA of a dark female from *Dark 51+2* strain. Lane 9: The PCR product amplified from the genomic DNA of a light female from *74L* strain. (B) The 625 bp region containing SNPs specific to the Dark pdm3 allele was amplified. The digits from 1 to 9 indicate the numbers of lanes in the agarose gel. The sample in each lane is as follows. Lane 1: 1 kb DNA ladder from New England Biolabs. Lane 2: The PCR product amplified from the genomic DNA of a light female from *58L* strain (a Light homozygous strain, already genotyped with amplicon sequencing). Lane 3: The PCR product amplified from the genomic DNA of a dark female from *Dark NHO* strain (a Dark homozygous strain, already genotyped with amplicon sequencing). Lane 4: The PCR product amplified from the genomic DNA of a light female from *Light 78L* strain. Lane 5: The PCR product amplified from the genomic DNA of a dark female from *Dark 51+2* strain. Lane 6: The PCR product amplified from the genomic DNA of a light female from *71L* strain. Lane 7: The PCR product amplified from the genomic DNA of a dark female from *103D* strain. Lane 8: The PCR product amplified from the genomic DNA of a light female from *74L* strain. The band is indicated with the white arrowhead. Lane 9: The PCR product amplified from the genomic DNA of a dark female from *60L* strain.

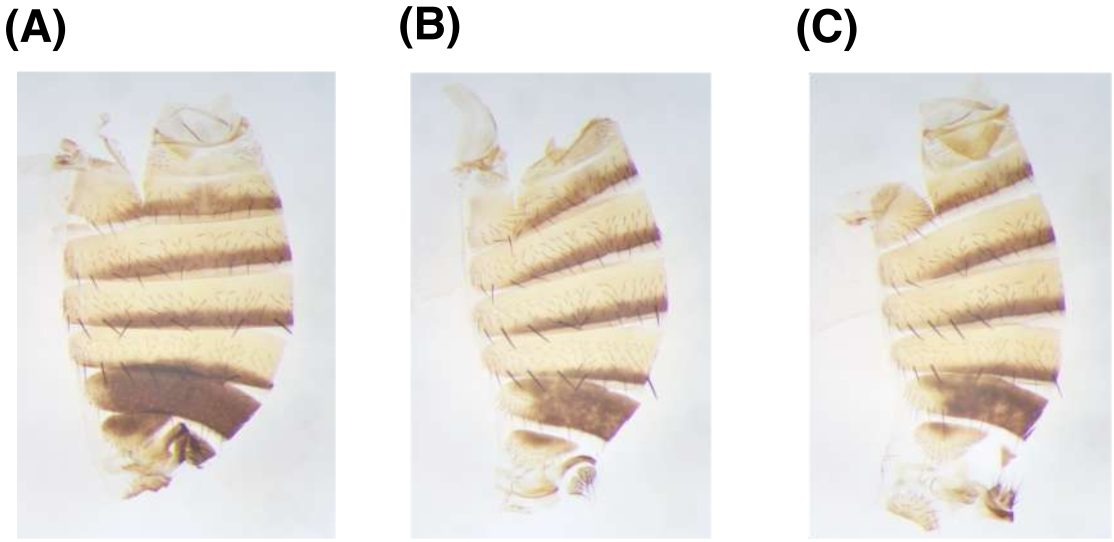


Figure. S3
Abdominal pigmentation of F1 females from the interspecific cross between *D. chauvacae* Dark l2 strain and *D. burlai* Light 11x strain. (A) An F1 female reared at 20 °C. The fully dark phenotype was observed. (B) An F1 female reared at 23 °C. Dominance of the Dark allele was incomplete. (C) An F1 female reared at 26 °C. Dominance of the Dark allele was incomplete.

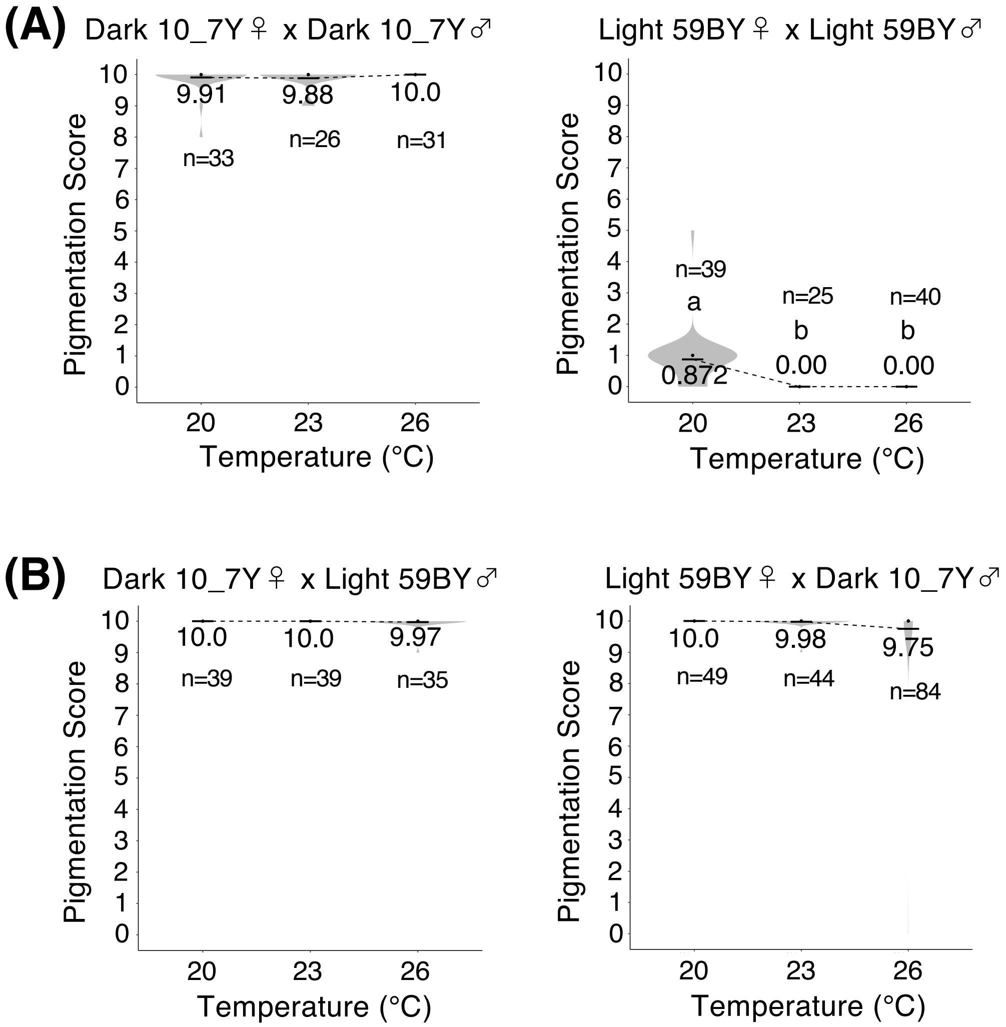


Figure. S4

The pigmentation scores for A7 segment of female abdomens of *D. rufa*, reared at 20 °C, 23 °C, and 26 °C. (A) The scores for Dark 10_7Y homozygotes (left) and Light 59BY homozygotes (right). There were significant differences between temperatures in Light 59BY homozygotes ($F = 38.41$, $p < 10^{-10}$, one-way ANOVA, degree of freedom = 2). (B) The scores for F1 hybrids from the crosses between females from Dark 10_7Y strain and males from Light 59BY strain (left), and between females from Light 59BY strain and males from Dark 10_7Y strain (right). There were no significant differences between temperatures in (one-way ANOVA, degree of freedom = 2). Black bars and black dots indicate mean values and median values for each. Mean values are written near the black bars. Different alphabets indicate significant differences ($p < 0.05$, Tukey's HSD test).

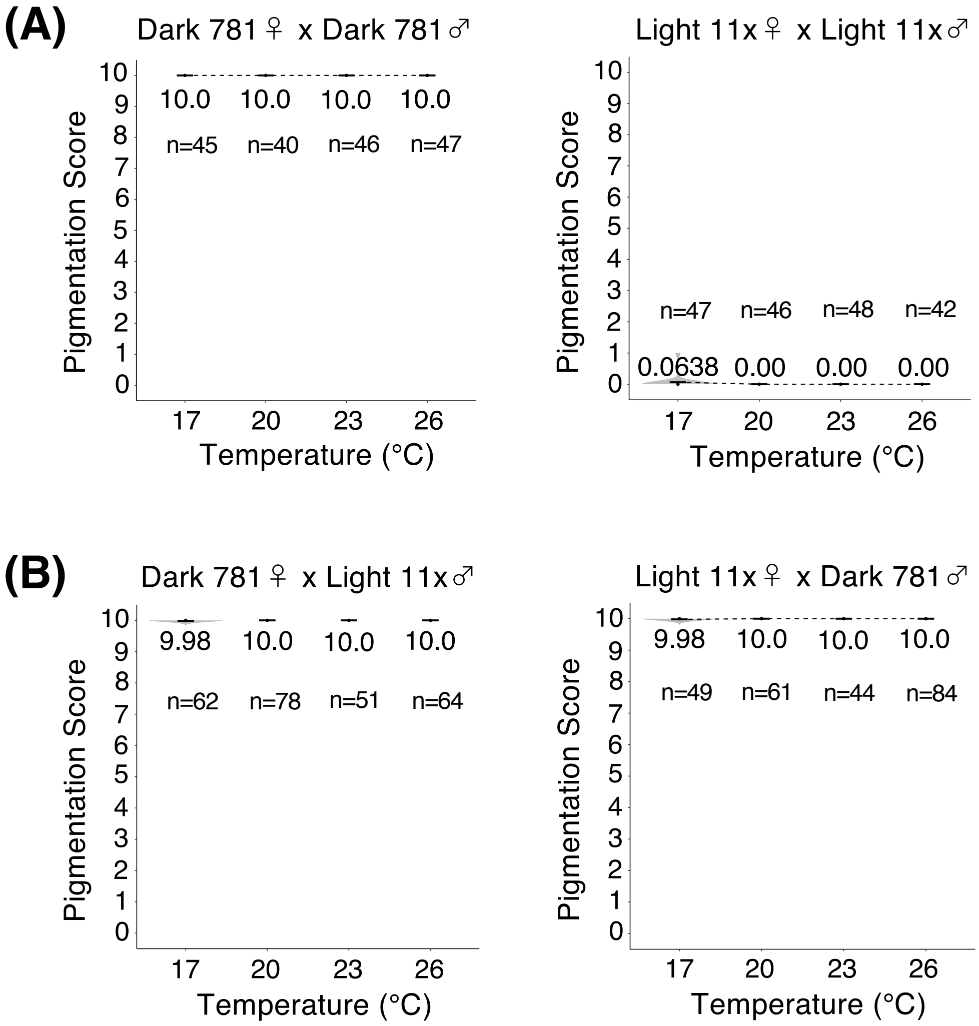


Figure. S5

The pigmentation scores for A7 segment of female abdomens of *D. burlai*, reared at 17 °C, 20 °C, 23 °C, and 26 °C. (A) The scores for Dark 781 homozygotes (left) and Light 11x homozygotes (right). In both categories, there were no significant differences between temperatures (one-way ANOVA, degree of freedom = 3). (B) The scores for F1 hybrids from the crosses between females from Dark 781 strain and males from Light 11x strain (left), and between females from Light 11x strain and males from Dark 781 strain (right). There were no significant differences between temperatures (one-way ANOVA, degree of freedom = 3). Black bars and black dots indicate mean values and median values for each. Mean values are written near the black bars. Different alphabets indicate significant differences ($p < 0.05$, Tukey's HSD test).

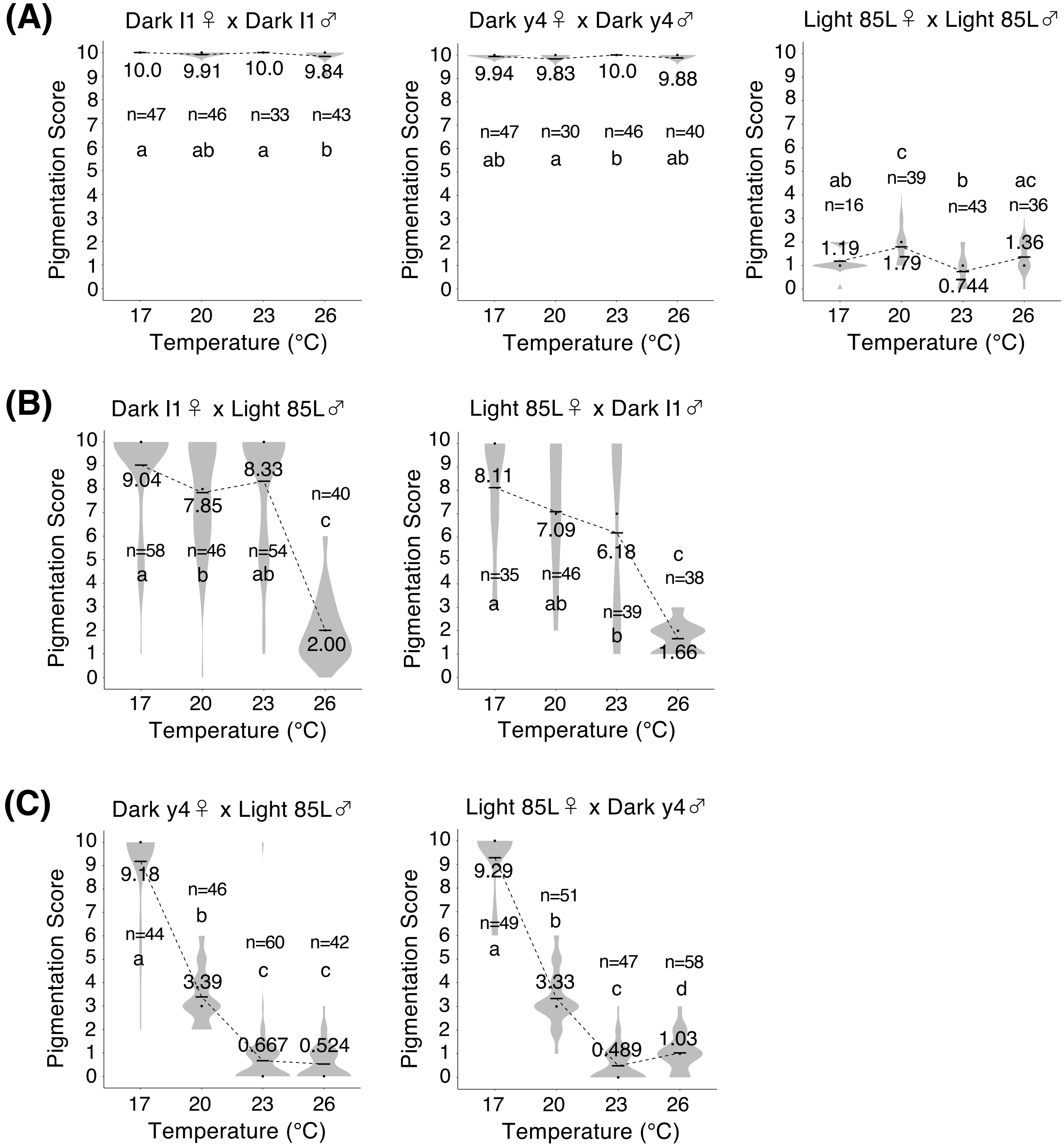

(A)
Dark I1♀ x Dark I1♂
10.0
9.91
10.0
9.84
n=47
n=46
n=33
n=43
a
ab
a
b
Dark y4♀ x Dark y4♂
9.94
9.83
10.0
9.88
n=47
n=30
n=46
n=40
ab
a
b
ab
Light 85L♀ x Light 85L♂
ab
c
b
ac
n=16
n=39
n=43
n=36
1.19
1.79
0.744
1.36
(B)
Dark I1♀ x Light 85L♂
9.04
7.85
8.33
2.00
n=58
n=46
n=54
n=40
a
b
ab
c
Light 85L♀ x Dark I1♂
8.11
7.09
6.18
1.66
n=35
n=46
n=39
n=38
a
ab
b
c
(C)
Dark y4♀ x Light 85L♂
9.18
3.39
0.667
0.524
n=44
n=46
n=60
n=42
a
b
c
c
Light 85L♀ x Dark y4♂
9.29
3.33
0.489
1.03
n=49
n=51
n=47
n=58
a
b
c
d
Pigmentation Score
0 1 2 3 4 5 6 7 8 9 10
Temperature (°C)
17 20 23 26


Figure. S6

Figure. S6

The pigmentation scores for A7 segment of female abdomens of *D.* cf. *bocqueti*, reared at 17 °C, 20 °C, 23 °C, and 26 °C. (A) The scores for homozygotes from Dark l1 strain (left), Dark y4 strain (middle), and Light 85L strain (right). There were significant differences between temperatures (one-way ANOVA, degree of freedom = 3). $F$ = 4.460, $p$ = 0.0049 for Dark l1 homozygotes. $F$ = 2.862, $p$ = 0.039 for Dark y4 homozygotes. $F$ = 14.41, $p < 10^{-7}$ for Light 85L homozygotes. (B) The scores for F1 hybrids from the crosses between females from Dark l1 strain and males from Light 85L strain (left), and between females from Light 85L strain and males from Dark l1 strain (right). There were significant differences between temperatures ($p < 10^{-10}$, one-way ANOVA, degree of freedom = 3). $F$ = 94.96 for hybrids between Dark l1 females and Light 85L males. $F$ = 42.93 for hybrids between Light 85l females and Dark l1 males. (C) The scores for F1 hybrids from the crosses between females from Dark y4 strain and males from Light 85L strain (left), and between females from Light 85L strain and males from Dark y4 strain (right). There were significant differences between temperatures ($p < 10^{-10}$, one-way ANOVA, degree of freedom = 3). $F$ = 381.7 for hybrids between Dark y4 females and Light 85L males. $F$ = 707.9 for hybrids between Light 85L females and Dark y4 males. Black bars and black dots indicate mean values and median values for each. Mean values are written near the black bars. Different alphabets indicate significant differences ($p < 0.05$, Tukey's HSD test).

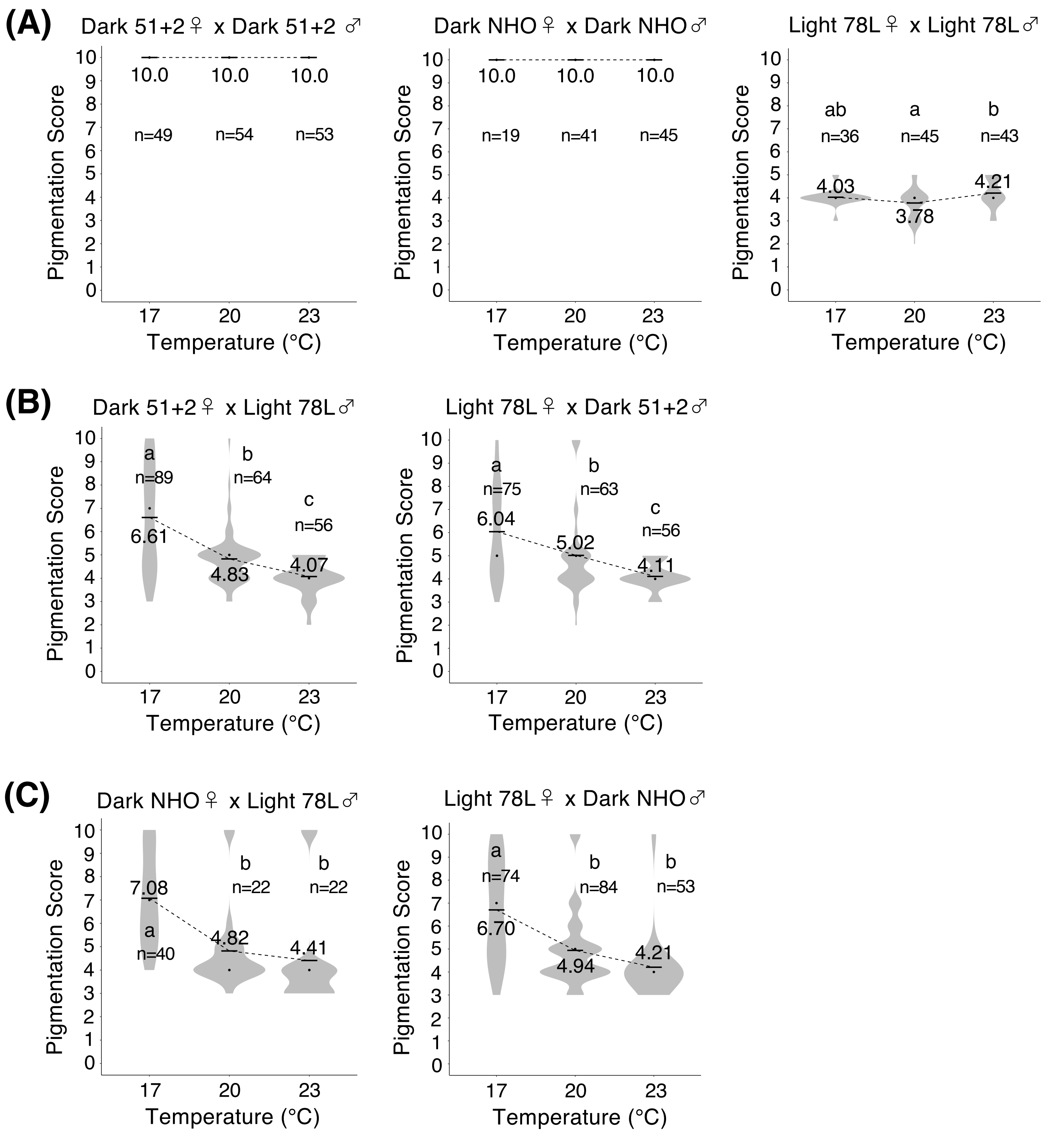

(A)
Dark 51+2♀ x Dark 51+2 ♂
10.0
10.0
10.0
n=49
n=54
n=53
Dark NHO♀ x Dark NHO♂
10.0
10.0
10.0
n=19
n=41
n=45
Light 78L♀ x Light 78L♂
ab
a
b
n=36
n=45
n=43
4.03
3.78
4.21
(B)
Dark 51+2♀ x Light 78L♂
a
n=89
b
n=64
c
n=56
6.61
4.83
4.07
Light 78L♀ x Dark 51+2♂
a
n=75
b
n=63
c
n=56
6.04
5.02
4.11
(C)
Dark NHO♀ x Light 78L♂
7.08
a
n=40
b
n=22
b
n=22
4.82
4.41
Light 78L♀ x Dark NHO♂
a
n=74
b
n=84
b
n=53
6.70
4.94
4.21
Pigmentation Score
Temperature (°C)
17
20
23


Figure. S7

Figure. S7

The pigmentation scores for A7 segments of female abdomens of *D. jambulina*, reared at 17 °C, 20 °C, and 23 °C. (A) The scores for homozygotes from Dark 51+2 strain (left), Dark NHO (middle), Light 78L strain (right). There was a significant difference between temperatures in Light 78L homozygotes ($F$ = 5.701, $p$ = 0.0043, one-way ANOVA, degree of freedom = 2). (B) The scores for F1 hybrids from the crosses between females from Dark 51+2 strain and males from Light 78L strain (left), and between females from Light 78L strain and males from Dark 51+2 strain (right). There were significant differences between temperatures (one-way ANOVA, degree of freedom = 2). $F$ = 47.73, $p < 10^{-10}$ for hybrids between Dark 51+2 females and Light 78L males. $F$ = 22.51, $p < 10^{-8}$ for hybrids between Light 78L females and Dark 51+2 males. (C) The scores for F1 hybrids from the crosses between females from Dark NHO strain and males from Light 78L strain (left), and between females from Light 78L strain and males from Dark NHO strain (right). There were significant differences between temperatures (one-way ANOVA, degree of freedom = 2). $F$ = 15.55, $p < 10^{-5}$ for hybrids between Dark NHO females and Light 78L males. $F$ = 33.28, $p < 10^{-10}$ for hybrids between Light 78L females and Dark NHO males. Black bars and black dots indicate mean values and median values for each. Mean values are written near the black bars. Different alphabets indicate significant differences ($p < 0.05$, Tukey's HSD test).

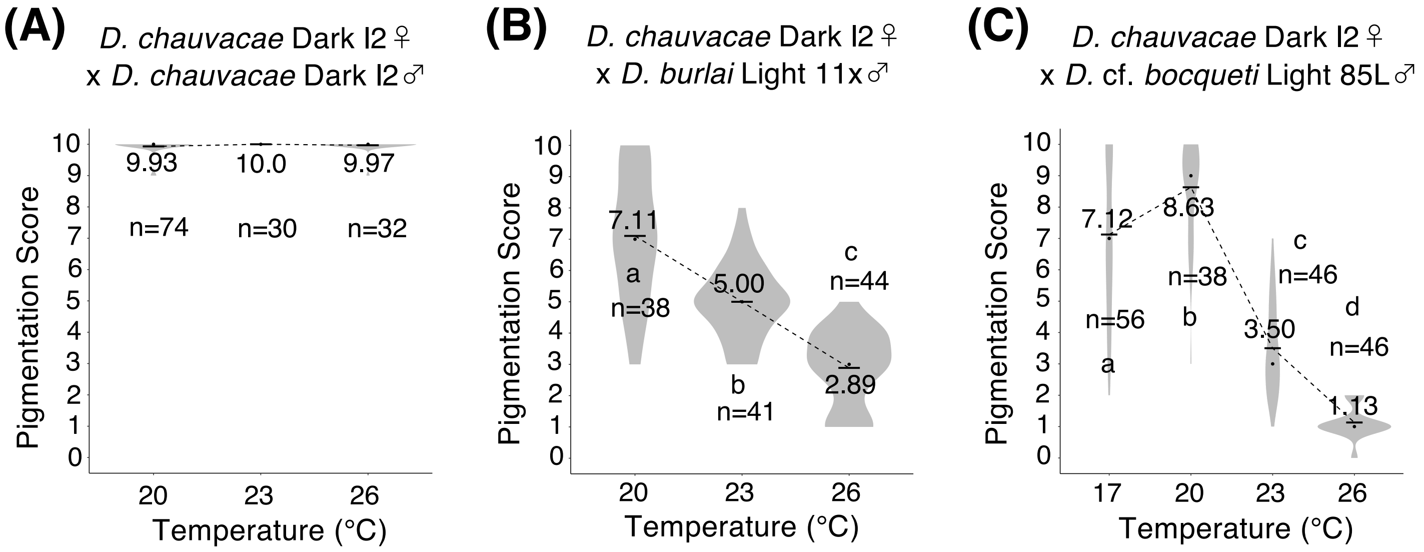


Figure. S8

The pigmentation scores for A7 segment of female abdomen of *D. chauvacae* and hybrids produced from interspecific crosses. (A) The score for Dark l2 homozygotes of *D. chauvacae*, reared at 20 °C, 23 °C, and 26 °C. There was no significant difference between temperatures (one-way ANOVA, degree of freedom = 2). (B) The pigmentation scores for F1 hybrids from the crosses between *D. chauvacae* females from Dark l2 strain and *D. burlai* males from Light 11x strain. The hybrid flies were reared at 20 °C, 23 °C, and 26 °C. There were significant differences between temperatures ($F = 77.92$, $p < 10^{-10}$, one-way ANOVA, degree of freedom = 2). (C) The pigmentation scores for F1 hybrids from the crosses between *D. chauvacae* females from Dark l2 strain and *D.* cf. *bocqueti* males from Light 85L strain. The hybrid flies were reared at 17 °C, 20 °C, 23 °C, and 26 °C. There were significant differences between temperatures ($F = 173.8$, $p < 10^{-10}$, one-way ANOVA, degree of freedom = 3). Black bars and black dots indicate mean values and median values for each. Mean values are written near the black bars. Different alphabets indicate significant differences ($p < 0.05$, Tukey's HSD test).

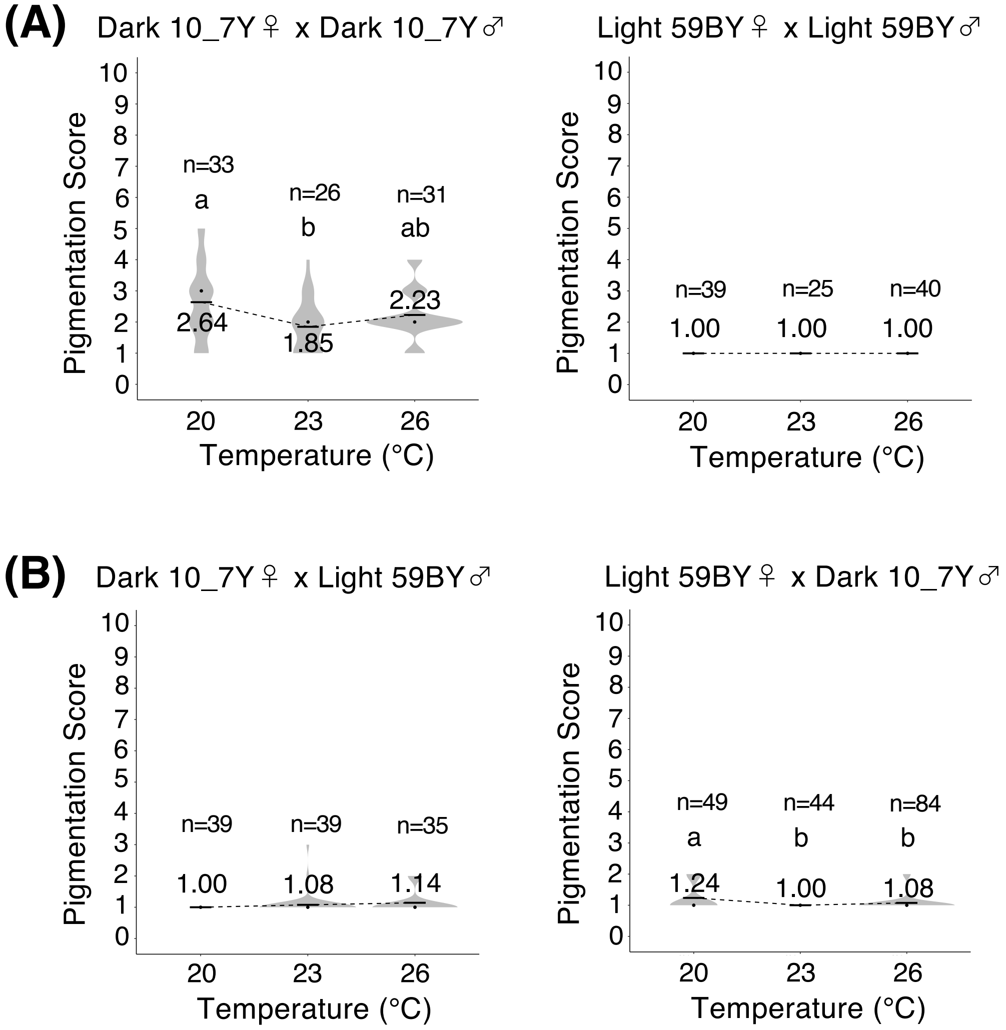


Figure. S9

The pigmentation scores for A5 segment of female abdomens of *D. rufa*, reared at 20 °C, 23 °C, and 26 °C. (A) The scores for Dark 10_7Y homozygotes (left) and Light 59BY homozygotes (right). There were significant differences between temperatures in Dark 10_7Y homozygotes ($F$ = 4.833, $p$ = 0.010, one-way ANOVA, degree of freedom = 2). (B) The scores for F1 hybrids from the crosses between females from Dark 10_7Y strain and males from Light 59BY strain (left), and between females from Light 59BY strain and males from Dark 10_7Y strain (right). There were significant differences between temperatures in hybrids between Light 59BY females and Dark 10_7Y males ($F$ = 6.879, $p$ = 0.0015, one-way ANOVA, degree of freedom = 2). Black bars and black dots indicate mean values and median values for each. Mean values are written near the black bars. Different alphabets indicate significant differences ($p$ < 0.05, Tukey's HSD test).

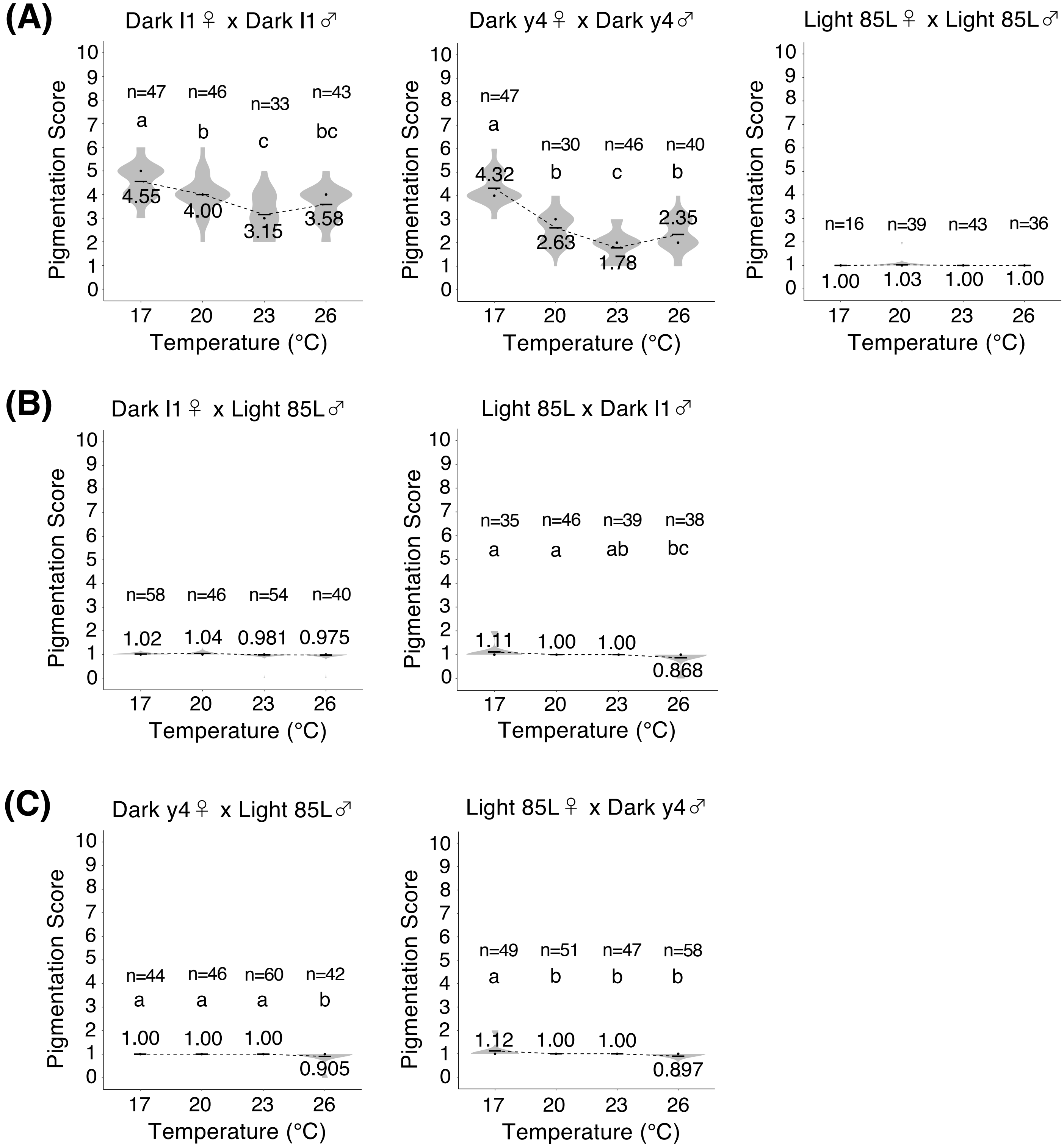

(A)
Dark I1♀ x Dark I1♂
n=47 n=46 n=33 n=43
a b c bc
4.55 4.00 3.15 3.58
Dark y4♀ x Dark y4♂
n=47 n=30 n=46 n=40
a b c b
4.32 2.63 1.78 2.35
Light 85L♀ x Light 85L♂
n=16 n=39 n=43 n=36
1.00 1.03 1.00 1.00
(B)
Dark I1♀ x Light 85L♂
n=58 n=46 n=54 n=40
1.02 1.04 0.981 0.975
Light 85L x Dark I1♂
n=35 n=46 n=39 n=38
a a ab bc
1.11 1.00 1.00 0.868
(C)
Dark y4♀ x Light 85L♂
n=44 n=46 n=60 n=42
a a a b
1.00 1.00 1.00 0.905
Light 85L♀ x Dark y4♂
n=49 n=51 n=47 n=58
a b b b
1.12 1.00 1.00 0.897
Pigmentation Score
Temperature (°C)
17 20 23 26


Figure. S10

Figure. S10

The pigmentation scores for A5 segment of female abdomens of *D.* cf. *bocqueti*, reared at 17 °C, 20 °C, 23 °C, and 26 °C. (A) The scores for homozygotes from Dark l1 strain (left), Dark y4 strain (middle), and Light 85L strain (right). There were significant differences between temperatures except Light 85L homozygotes ($p < 10^{-10}$, one-way ANOVA, degree of freedom = 3). $F = 21.99$ for Dark l1 homozygotes. $F = 55.00$ for Dark y4 homozygotes. (B) The scores for F1 hybrids from the crosses between females from Dark l1 strain and males from Light 85L strain (left), and between females from Light 85L strain and males from Dark l1 strain (right). There were significant differences between temperatures in hybrids between Light 85L females and Dark l1 males $F = 7.218$, ($p < 0.01$, one-way ANOVA, degree of freedom = 3). (C) The scores for F1 hybrids from the crosses between females from Dark y4 strain and males from Light 85L strain (left), and between females from Light 85L strain and males from Dark y4 strain (right). There were significant differences between temperatures ($p < 0.01$, one-way ANOVA, degree of freedom = 3). $F = 5.154$ for hybrids between Dark y4 females and Light 85L males. $F = 8.531$ for hybrids between Light 85L females and Dark y4 males. Black bars and black dots indicate mean values and median values for each. Mean values are written near the black bars. Different alphabets indicate significant differences ($p < 0.05$, Tukey's HSD test).

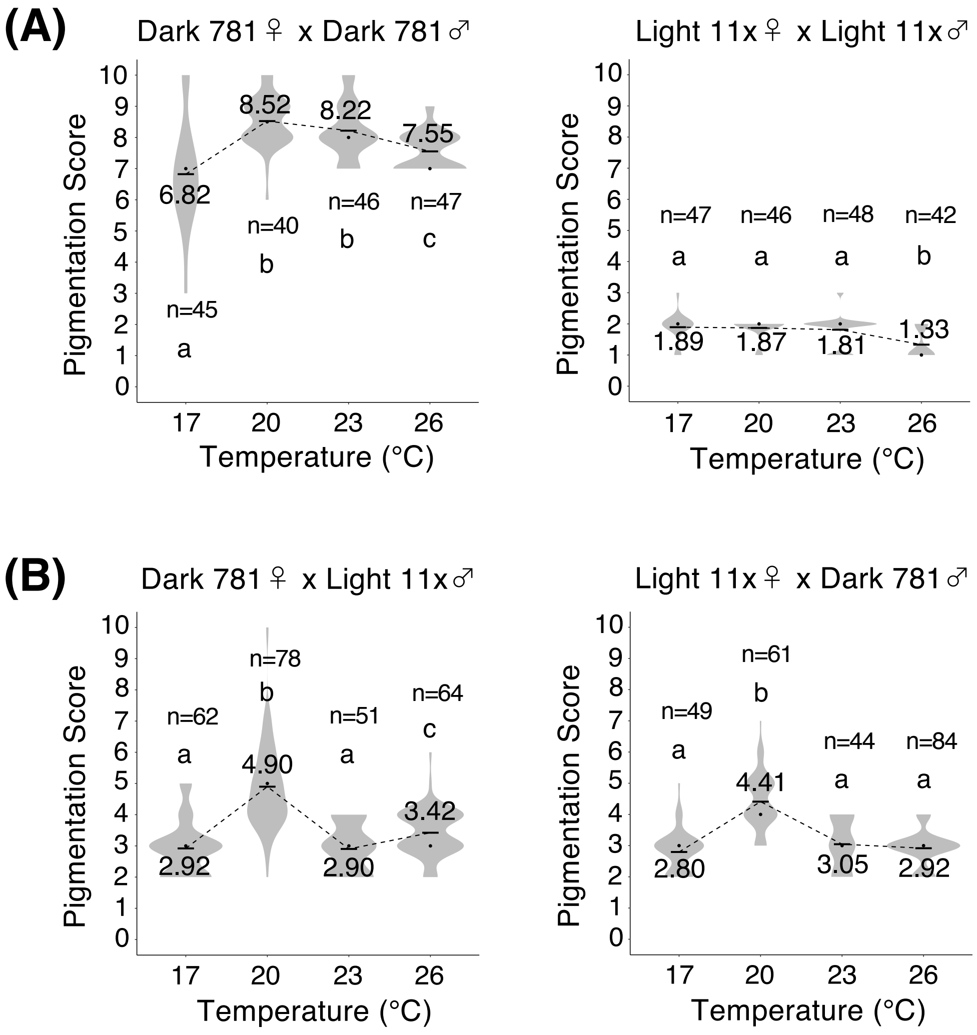


Figure. S11

The pigmentation scores for A5 segment of female abdomens of *D. burlai*, reared at 17 °C, 20 °C, 23 °C, and 26 °C. (A) The scores for Dark 781 homozygotes (left) and Light 11x homozygotes (right). There were significant differences between temperatures ($p < 10^{-7}$, one-way ANOVA, degree of freedom = 3). $F = 20.81$ for Dark 781 homozygotes. $F = 14.13$ for Light 11x homozygotes. (B) The scores for F1 hybrids from the crosses between females from Dark 781 strain and males from Light 11x strain (left), and between females from Light 11x strain and males from Dark 781 strain (right). There were significant differences between temperatures ($p < 10^{-10}$, one-way ANOVA, degree of freedom = 3). $F = 61.41$ for hybrids between Dark 781 females and Light 11x males. $F = 63.11$ for hybrids between Light 11x females and Dark 781 males. Black bars and black dots indicate mean values and median values for each. Mean values are written near the black bars. Different alphabets indicate significant differences ($p < 0.05$, Tukey's HSD test).

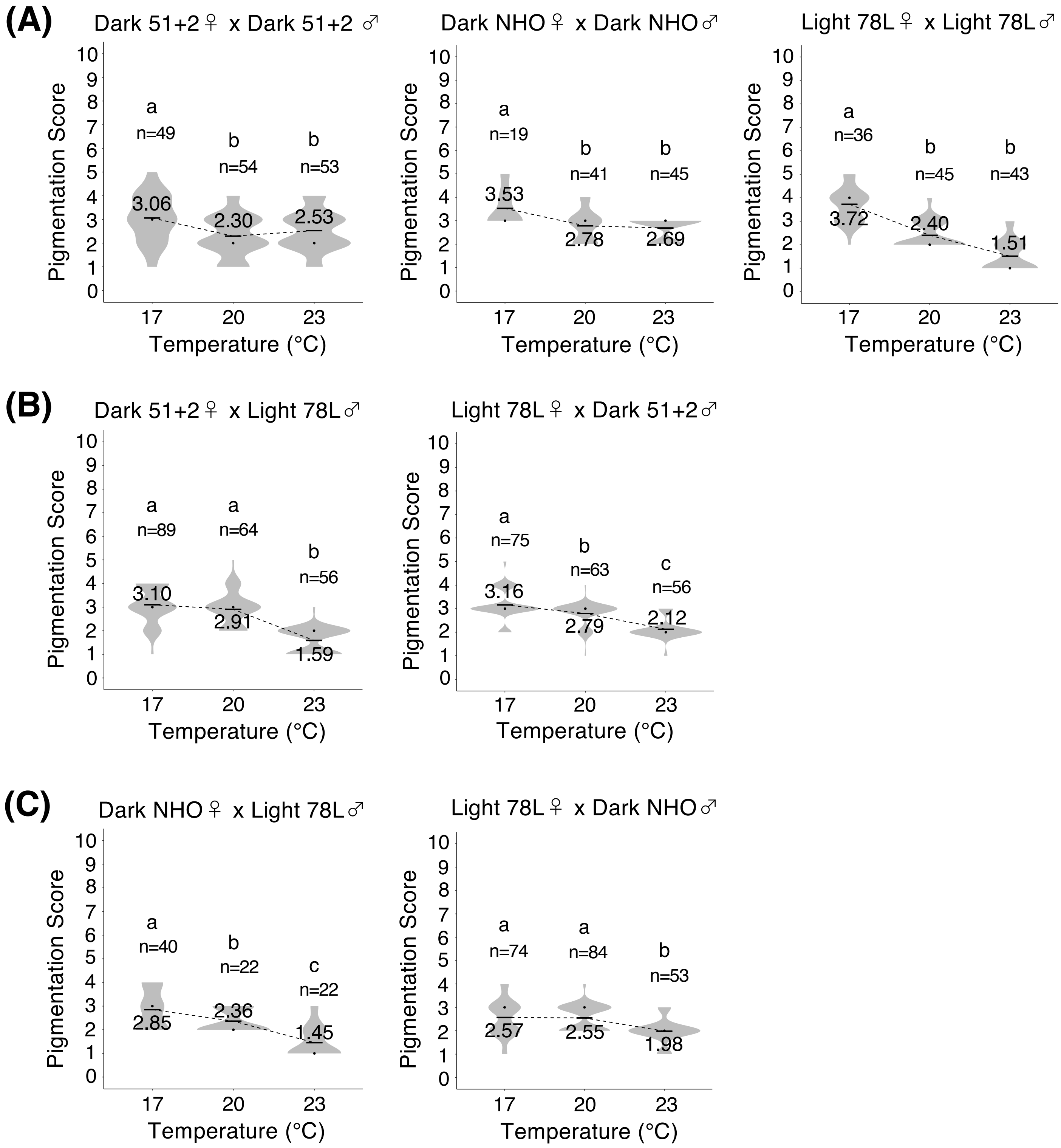

(A)
Dark 51+2♀ x Dark 51+2 ♂
Dark NHO♀ x Dark NHO♂
Light 78L♀ x Light 78L♂
(B)
Dark 51+2♀ x Light 78L♂
Light 78L♀ x Dark 51+2♂
(C)
Dark NHO♀ x Light 78L♂
Light 78L♀ x Dark NHO♂
Pigmentation Score
Temperature (°C)
17
20
23
a
b
c
n=49
n=54
n=53
3.06
2.30
2.53
n=19
n=41
n=45
3.53
2.78
2.69
n=36
n=45
n=43
3.72
2.40
1.51
n=89
n=64
n=56
3.10
2.91
1.59
n=75
n=63
n=56
3.16
2.79
2.12
n=40
n=22
n=22
2.85
2.36
1.45
n=74
n=84
n=53
2.57
2.55
1.98


Figure. S12

Figure. S12

The pigmentation scores for A5 segments of female abdomens of *D. jambulina*, reared at 17 °C, 20 °C, and 23 °C. (A) The scores for homozygotes from Dark 51+2 strain (left), Dark NHO (middle), Light 78L strain (right). There were significant differences between temperatures (one-way ANOVA, degree of freedom = 2). $F$ = 9.55, $p < 10^{-3}$ for Dark 51+2 homozygotes. $F$ = 12.92 $p < 10^{-5}$ for Dark NHO homozygotes. $F$ = 110.3, $p < 10^{-10}$ for Light 78L homozygotes. (B) The scores for F1 hybrids from the crosses between females from Dark 51+2 strain and males from Light 78L strain (left), and between females from Light 78L strain and males from Dark 51+2 strain (right). There were significant differences between temperatures ($p < 10^{-10}$, one-way ANOVA, degree of freedom = 2). $F$ = 88.66 for hybrids between Dark 51+2 females and Light 78L males. $F$ = 58.45 for hybrids between Light 78L females and Dark 51+2 males. (C) The scores for F1 hybrids from the crosses between females from Dark NHO strain and males from Light 78L strain (left), and between females from Light 78L strain and males from Dark nho strain (right). There were significant differences between temperatures (one-way ANOVA, degree of freedom = 2). $F$ = 26.88, $p < 10^{-8}$ for hybrids between Dark NHO females and Light 78L males. $F$ = 16.32, $p < 10^{-6}$ for hybrids between Light 78L females and Dark NHO males. Black bars and black dots indicate mean values and median values for each. Mean values are written near the black bars. Different alphabets indicate significant differences ($p < 0.05$, Tukey's HSD test).

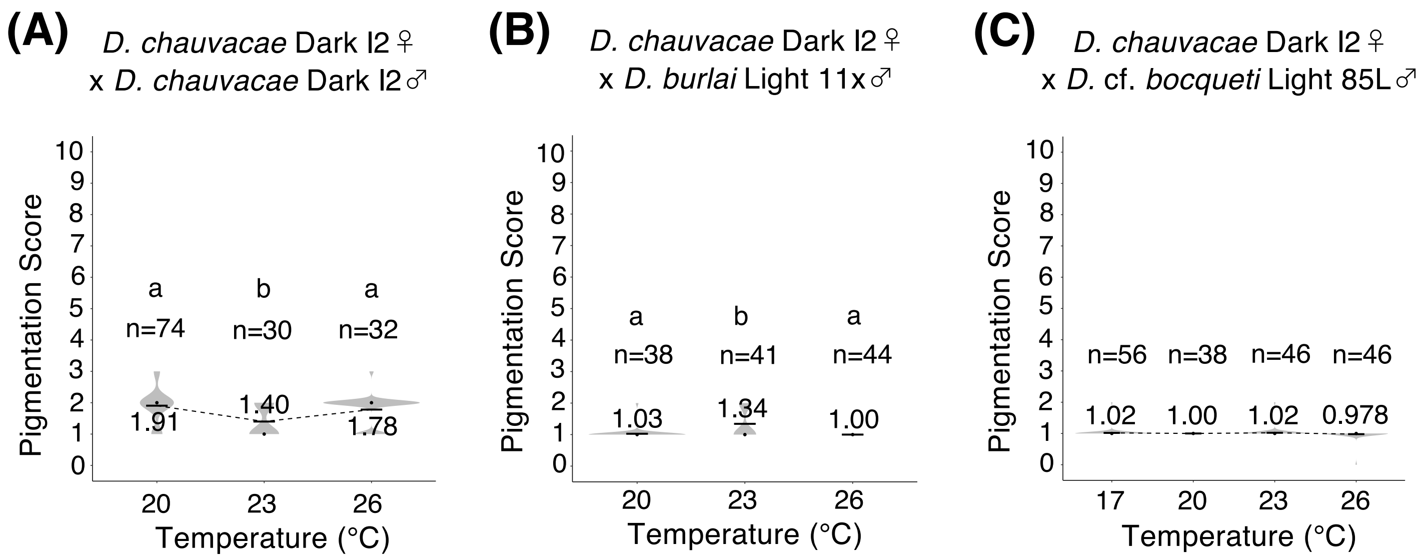


Figure. S13

The pigmentation scores for A5 segment of female abdomen of *D. chauvacae* and hybrids produced from interspecific crosses. (A) The score for Dark l2 homozygotes of *D. chauvacae*, reared at 20 °C, 23 °C, and 26 °C. There were significant differences between temperatures ($F = 9.315$, $p < 10^{-3}$, one-way ANOVA, degree of freedom = 2). (B) The pigmentation scores for F1 hybrids from the crosses between *D. chauvacae* females from Dark l2 strain and *D. burlai* males from Light 11x strain. The hybrid flies were reared at 20 °C, 23 °C, and 26 °C. There were significant differences between temperatures ($F = 17.53$, $p < 10^{-6}$, one-way ANOVA, degree of freedom = 2). (C) The pigmentation scores for F1 hybrids from the crosses between *D. chauvacae* females from Dark l2 strain and *D. bocqueti* males from Light 85L strain. The hybrid flies were reared at 17 °C, 20 °C, 23 °C, and 26 °C. There was no significant difference between temperatures (one-way ANOVA, degree of freedom = 3). Black bars and black dots indicate mean values and median values for each. Mean values are written near the black bars. Different alphabets indicate significant differences ($p < 0.05$, Tukey's HSD test).

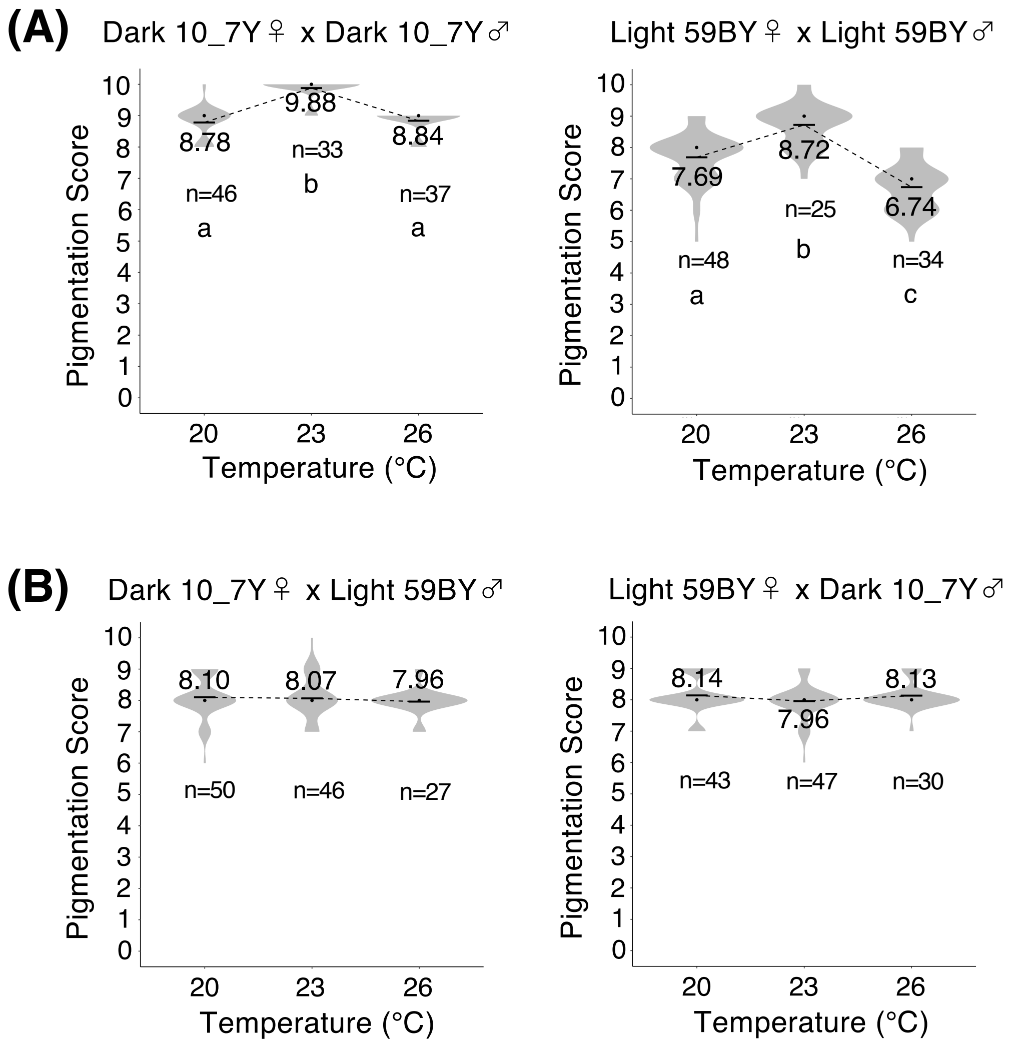


Figure. S14

The pigmentation scores for A6 segment of male abdomens of *D. rufa*, reared at 20 °C, 23 °C, and 26 °C. (A) The scores for Dark 10_7Y homozygotes (left) and Light 59BY homozygotes (right). In both categories, there were significant differences between temperatures ($p < 10^{-10}$, one-way ANOVA, degree of freedom = 2). $F = 75.38$ for Dark 10_7Y homozygotes, and $F = 49.49$ for Light 59BY homozygotes. (B) The scores for F1 hybrids from the crosses between females from Dark 10_7Y strain and males from Light 59BY strain (left), and between females from Light 59BY strain and males from Dark 10_7Y strain (right). There were no significant differences between temperatures (one-way ANOVA, degree of freedom = 2). Black bars and black dots indicate mean values and median values for each. Mean values are written near the black bars. Different alphabets indicate significant differences ($p < 0.05$, Tukey's HSD test).

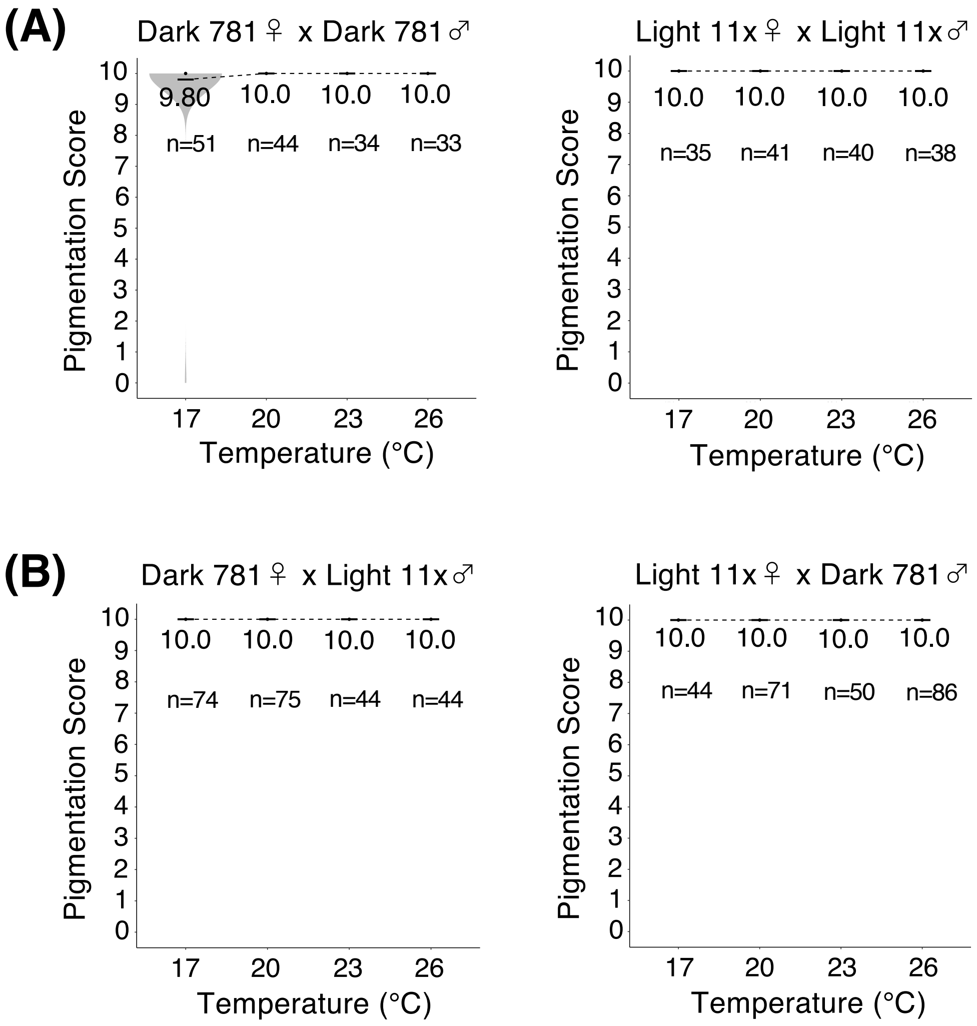


Figure. S15

The pigmentation scores for A6 segment of male abdomens of *D. burlai*, reared at 17 °C, 20 °C, 23 °C, and 26 °C. (A) The scores for Dark 781 homozygotes (left) and Light 11x homozygotes (right). In both categories, there were no significant differences between temperatures (one-way ANOVA, degree of freedom = 3). (B) The scores for F1 hybrids from the crosses between females from Dark 781 strain and males from Light 11x strain (left), and between females from Light 11x strain and males from Dark 781 strain (right). There were no significant differences between temperatures (one-way ANOVA, degree of freedom = 3). Black bars and black dots indicate mean values and median values for each. Mean values are written near the black bars. Different alphabets indicate significant differences ($p < 0.05$, Tukey's HSD test).

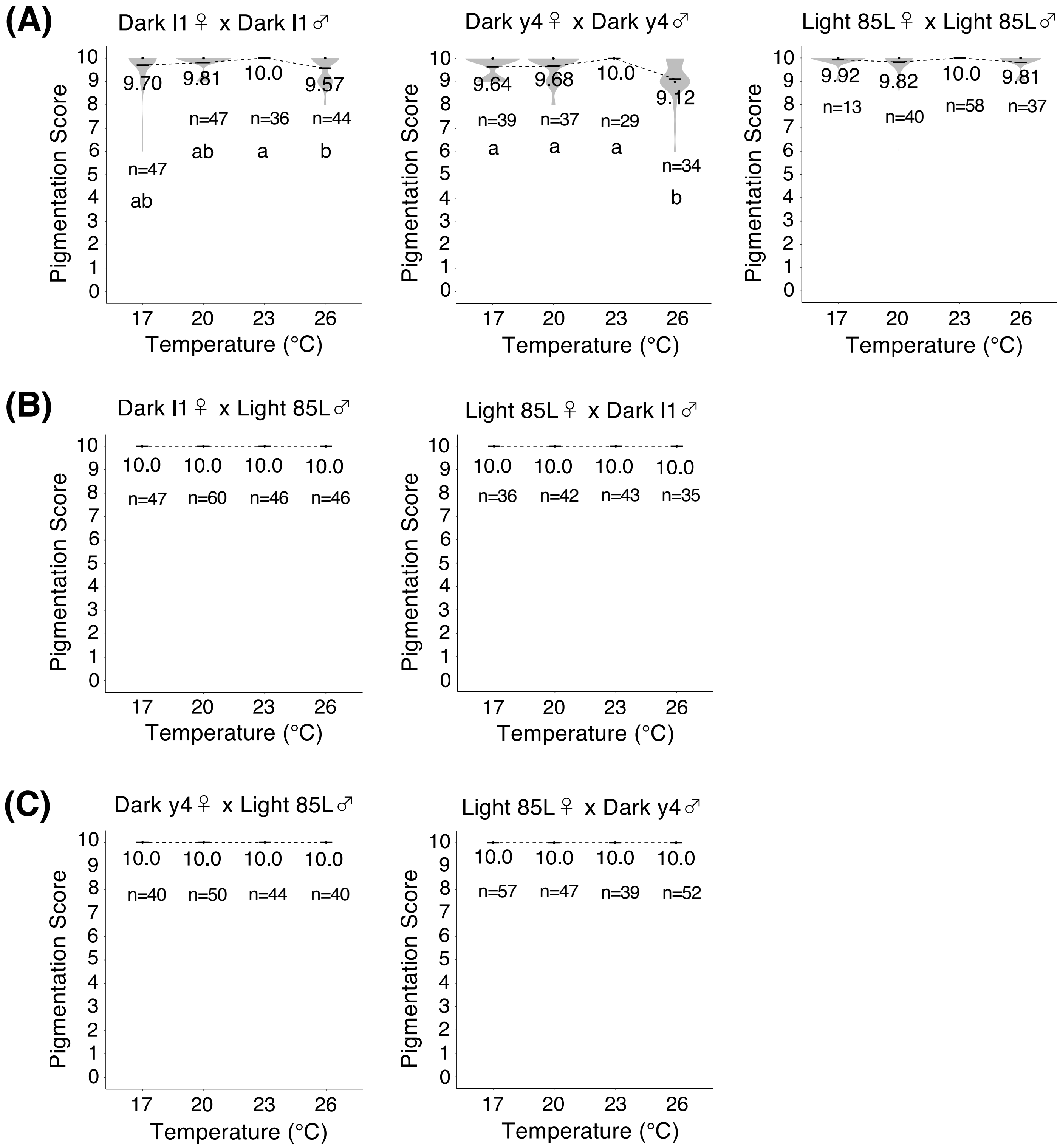
(A)
Dark I1♀ x Dark I1♂
Pigmentation Score
9.70
9.81
10.0
9.57
n=47
n=47
n=36
n=44
ab
ab
a
b
Temperature (°C)
Dark y4♀ x Dark y4♂
9.64
9.68
10.0
9.12
n=39
n=37
n=29
n=34
a
a
a
b
Light 85L♀ x Light 85L♂
9.92
9.82
10.0
9.81
n=13
n=40
n=58
n=37
(B)
Dark I1♀ x Light 85L♂
10.0
10.0
10.0
10.0
n=47
n=60
n=46
n=46
Light 85L♀ x Dark I1♂
10.0
10.0
10.0
10.0
n=36
n=42
n=43
n=35
(C)
Dark y4♀ x Light 85L♂
10.0
10.0
10.0
10.0
n=40
n=50
n=44
n=40
Light 85L♀ x Dark y4♂
10.0
10.0
10.0
10.0
n=57
n=47
n=39
n=52

Figure. S16

Figure. S16

The pigmentation scores for A6 segment of male abdomens of *D.* cf. *bocqueti*, reared at 17 °C, 20 °C, 23 °C, and 26 °C. (A) The scores for homozygotes from Dark l1 strain (left), Dark y4 strain (middle), and Light 85L strain (right). There were significant differences between temperatures except Light 85L homozygotes (one-way ANOVA, degree of freedom = 3). $F = 4.497$, $p = 0.0046$ for Dark l1 homozygotes. $F = 12.38$, $p < 10^{-6}$ for Dark y4 homozygotes. (B) The scores for F1 hybrids from the crosses between females from Dark l1 strain and males from Light 85L strain (left), and between females from Light 85L strain and males from Dark l1 strain (right). In both categories, there were no significant differences between temperatures (one-way ANOVA, degree of freedom = 3). (C) The scores for F1 hybrids from the crosses between females from Dark y4 strain and males from Light 85L strain (left), and between females from Light 85L strain and males from Dark y4 strain (right). There were no significant differences between temperatures (one-way ANOVA, degree of freedom = 3). Black bars and black dots indicate mean values and median values for each. Mean values are written near the black bars. Different alphabets indicate significant differences ($p < 0.05$, Tukey's HSD test).

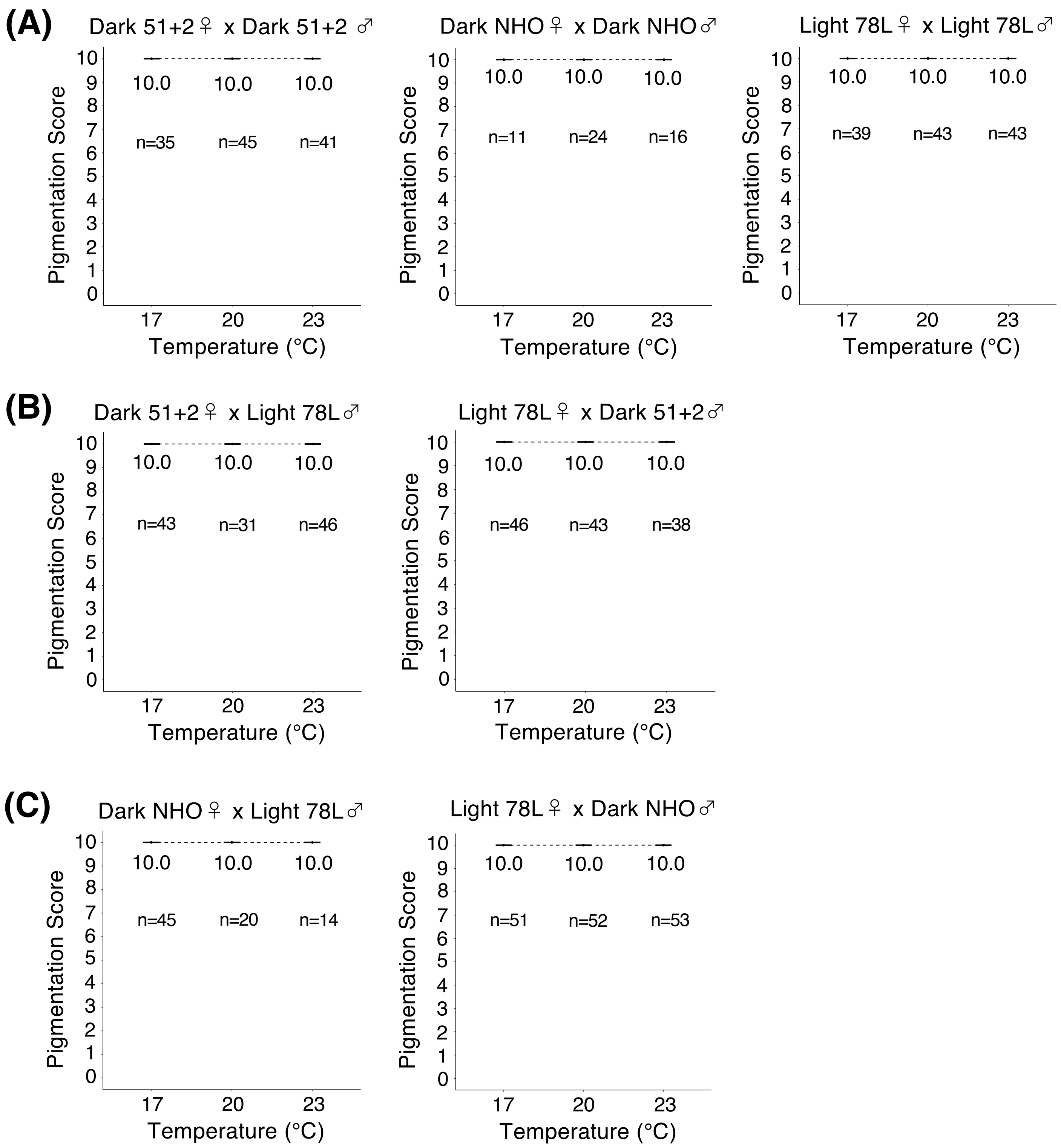
(A)
Dark 51+2♀ x Dark 51+2 ♂
Pigmentation Score
10
9
8
7
6
5
4
3
2
1
0
10.0
10.0
10.0
n=35
n=45
n=41
17
20
23
Temperature (°C)
Dark NHO♀ x Dark NHO♂
10.0
10.0
10.0
n=11
n=24
n=16
Light 78L♀ x Light 78L♂
10.0
10.0
10.0
n=39
n=43
n=43
(B)
Dark 51+2♀ x Light 78L♂
10.0
10.0
10.0
n=43
n=31
n=46
Light 78L♀ x Dark 51+2♂
10.0
10.0
10.0
n=46
n=43
n=38
(C)
Dark NHO♀ x Light 78L♂
10.0
10.0
10.0
n=45
n=20
n=14
Light 78L♀ x Dark NHO♂
10.0
10.0
10.0
n=51
n=52
n=53

Figure. S17

Figure. S17

The pigmentation scores for A6 segment of male abdomens of *D. jambulina*, reared at 17 °C, 20 °C, and 23 °C. (A) The scores for homozygotes from Dark 51+2 strain (left), Dark NHO (middle), Light 78L strain (right). There were no significant differences between temperatures (one-way ANOVA, degree of freedom = 2). (B) The scores for F1 hybrids from the crosses between females from Dark 51+2 strain and males from Light 78L strain (left), and between females from Light 78L strain and males from Dark 51+2 strain (right). There were no significant differences between temperatures (one-way ANOVA, degree of freedom = 2). (C) The scores for F1 hybrids from the crosses between females from Dark NHO strain and males from Light 78L strain (left), and between females from Light 78L strain and males from Dark NHO strain (right). There were no significant differences between temperatures (one-way ANOVA, degree of freedom = 2). Black bars and black dots indicate mean values and median values for each. Mean values are written near the black bars.

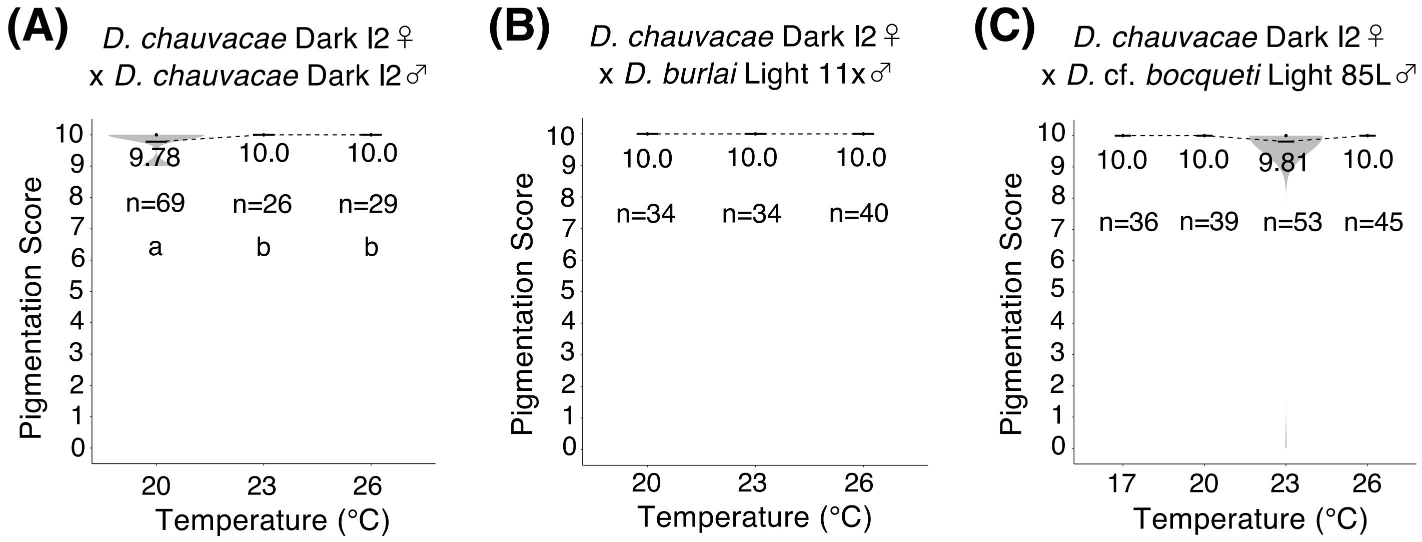


Figure. S18

The pigmentation scores for A6 segment of male abdomen of *D. chauvacae* and hybrids produced from interspecific crosses. (A) The score for Dark l2 homozygotes of *D. chauvacae*, reared at 20 °C, 23 °C, and 26 °C. There were significant differences between temperatures ($F = 7.454$, $p < 10^{-3}$, one-way ANOVA, degree of freedom = 2). (B) The pigmentation scores for F1 hybrids from the crosses between *D. chauvacae* females from Dark l2 strain and *D. burlai* males from Light 11x strain. The hybrid flies were reared at 20 °C, 23 °C, and 26 °C. There was no significant difference between temperatures (one-way ANOVA, degree of freedom = 2). (C) The pigmentation scores for F1 hybrids from the crosses between *D. chauvacae* females from Dark l2 strain and *D. bocqueti* males from Light 85L strain. The hybrid flies were reared at 17 °C, 20 °C, 23 °C, and 26 °C. There was no significant difference between temperatures (one-way ANOVA, degree of freedom = 3). Black bars and black dots indicate mean values and median values for each. Mean values are written near the black bars. Different alphabets indicate significant differences ($p < 0.05$, Tukey's HSD test).

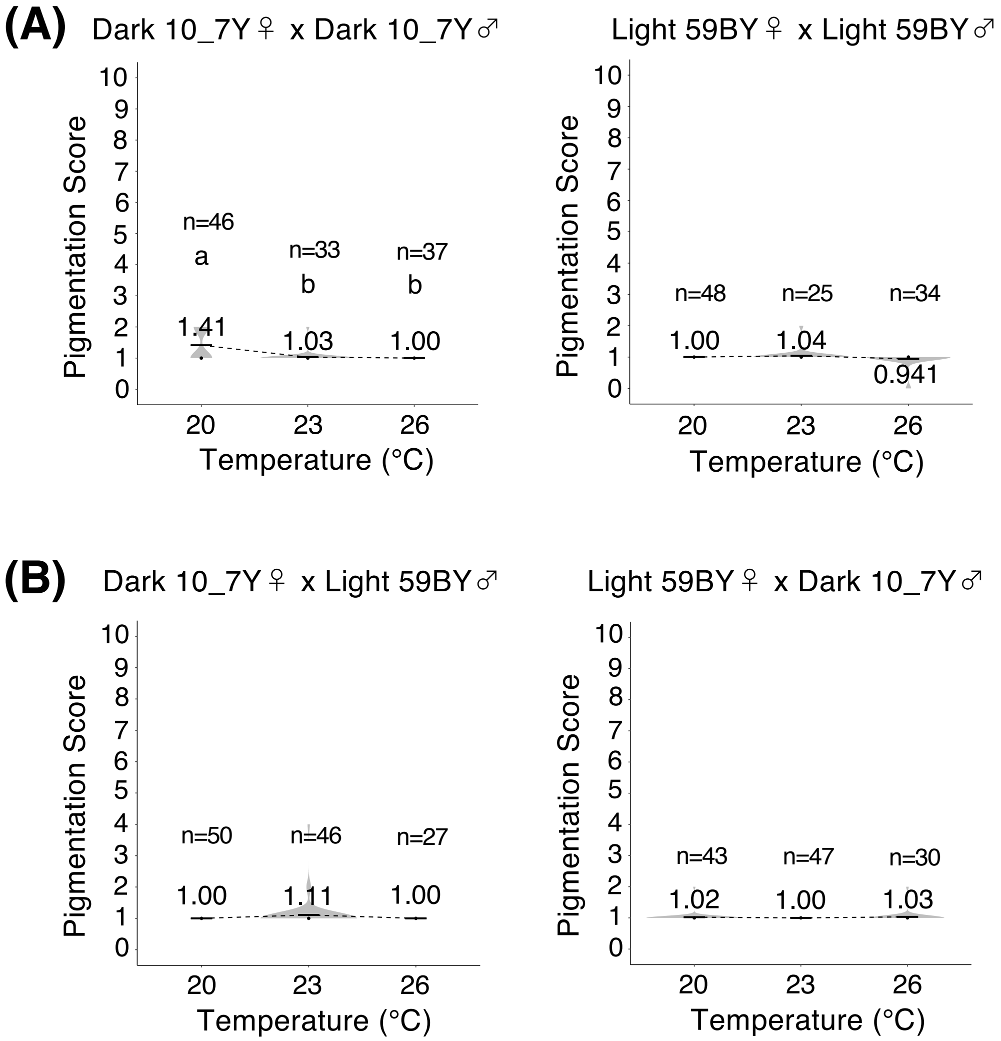


Figure. S19
The pigmentation scores for A5 segment of male abdomens of *D. rufa*, reared at 20 °C, 23 °C, and 26 °C. (A) The scores for Dark 10_7Y homozygotes (left) and Light 59BY homozygotes (right). There were significant differences between temperatures in Dark 10_7Y homozygotes ($F = 20.65$, $p < 10^{-7}$, one-way ANOVA, degree of freedom = 2). (B) The scores for F1 hybrids from the crosses between females from Dark 10_7Y strain and males from Light 59BY strain (left), and between females from Light 59BY strain and males from Dark 10_7Y strain (right). There were no significant differences between temperatures (one-way ANOVA, degree of freedom = 2). Black bars and black dots indicate mean values and median values for each. Mean values are written near the black bars. Different alphabets indicate significant differences ($p < 0.05$, Tukey's HSD test).

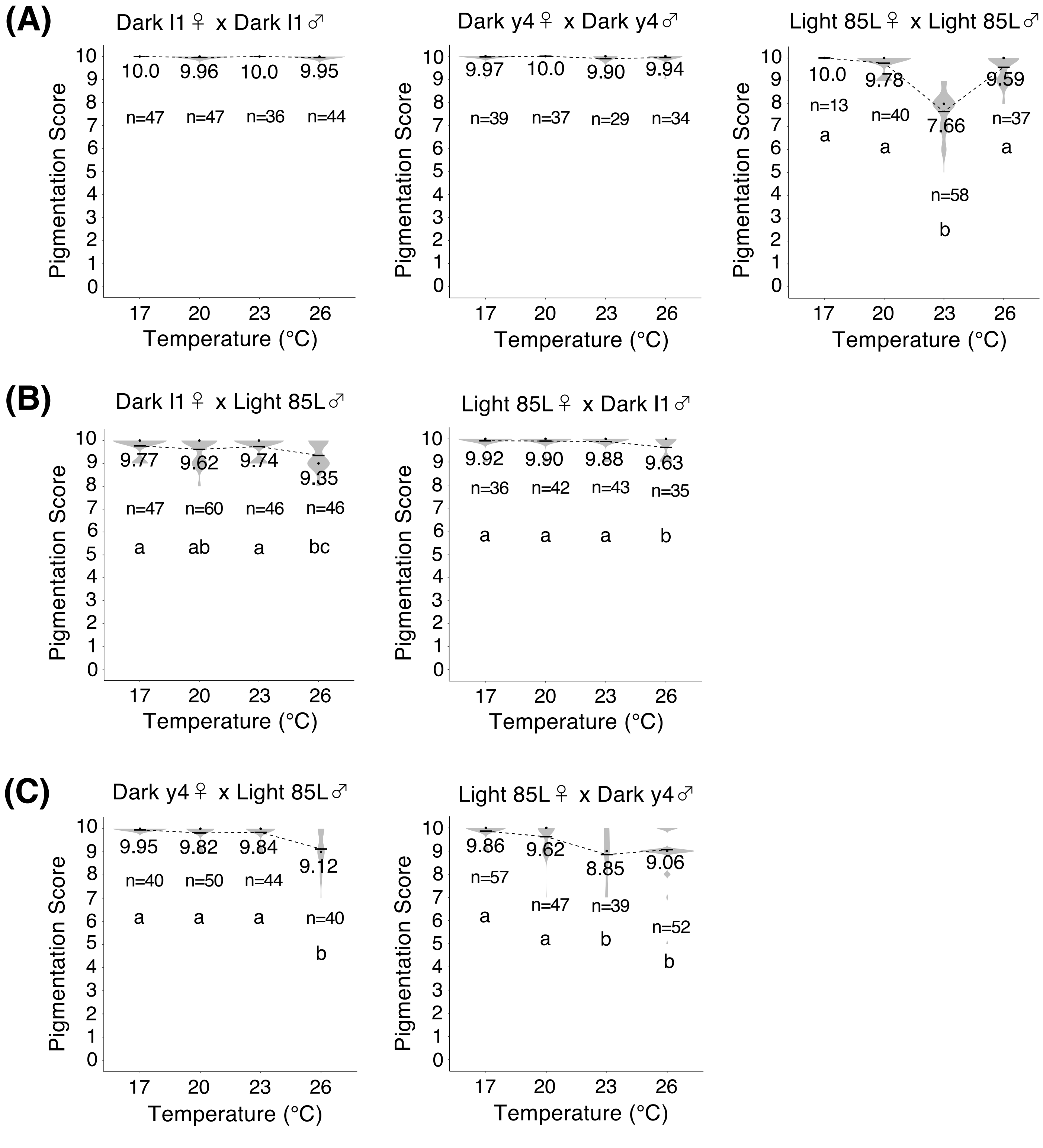
(A)
Dark I1♀ x Dark I1♂
10.0 9.96 10.0 9.95
n=47 n=47 n=36 n=44
Dark y4♀ x Dark y4♂
9.97 10.0 9.90 9.94
n=39 n=37 n=29 n=34
Light 85L♀ x Light 85L♂
10.0 9.78 7.66 9.59
n=13 n=40 n=58 n=37
a a b a
(B)
Dark I1♀ x Light 85L♂
9.77 9.62 9.74 9.35
n=47 n=60 n=46 n=46
a ab a bc
Light 85L♀ x Dark I1♂
9.92 9.90 9.88 9.63
n=36 n=42 n=43 n=35
a a a b
(C)
Dark y4♀ x Light 85L♂
9.95 9.82 9.84 9.12
n=40 n=50 n=44 n=40
a a a b
Light 85L♀ x Dark y4♂
9.86 9.62 8.85 9.06
n=57 n=47 n=39 n=52
a a b b
Pigmentation Score
0 1 2 3 4 5 6 7 8 9 10
Temperature (°C)
17 20 23 26

Figure. S20

Figure. S20

The pigmentation scores for A5 segment of male abdomens of *D.* cf. *bocqueti*, reared at 17 °C, 20 °C, 23 °C, and 26 °C. (A) The scores for homozygotes from Dark l1 strain (left), Dark y4 strain (middle), and Light 85L strain (right). There were significant differences between temperatures in Light 85L homozygotes ($F$ = 107.2, $p < 10^{-10}$, one-way ANOVA, degree of freedom = 3). (B) The scores for F1 hybrids from the crosses between females from Dark l1 strain and males from Light 85L strain (left), and between females from Light 85L strain and males from Dark l1 strain (right). There were significant differences between temperatures ($p < 0.01$, one-way ANOVA, degree of freedom = 3). $F$ = 5.884 for hybrids between Dark l1 females and Light 85L males. $F$ = 5.437 for hybrids between Light 85L females and Dark l1 males. (C) The scores for F1 hybrids from the crosses between females from Dark y4 strain and males from Light 85L strain (left), and between females from Light 85L strain and males from Dark y4 strain (right). There were significant differences between temperatures ($p < 10^{-10}$, one-way ANOVA, degree of freedom = 3). $F$ = 22.53 for hybrids between Dark y4 females and Light 85L males. $F$ = 19.04 for hybrids between Light 85L females and Dark y4 males. Black bars and black dots indicate mean values and median values for each. Mean values are written near the black bars. Different alphabets indicate significant differences ($p < 0.05$, Tukey's HSD test).

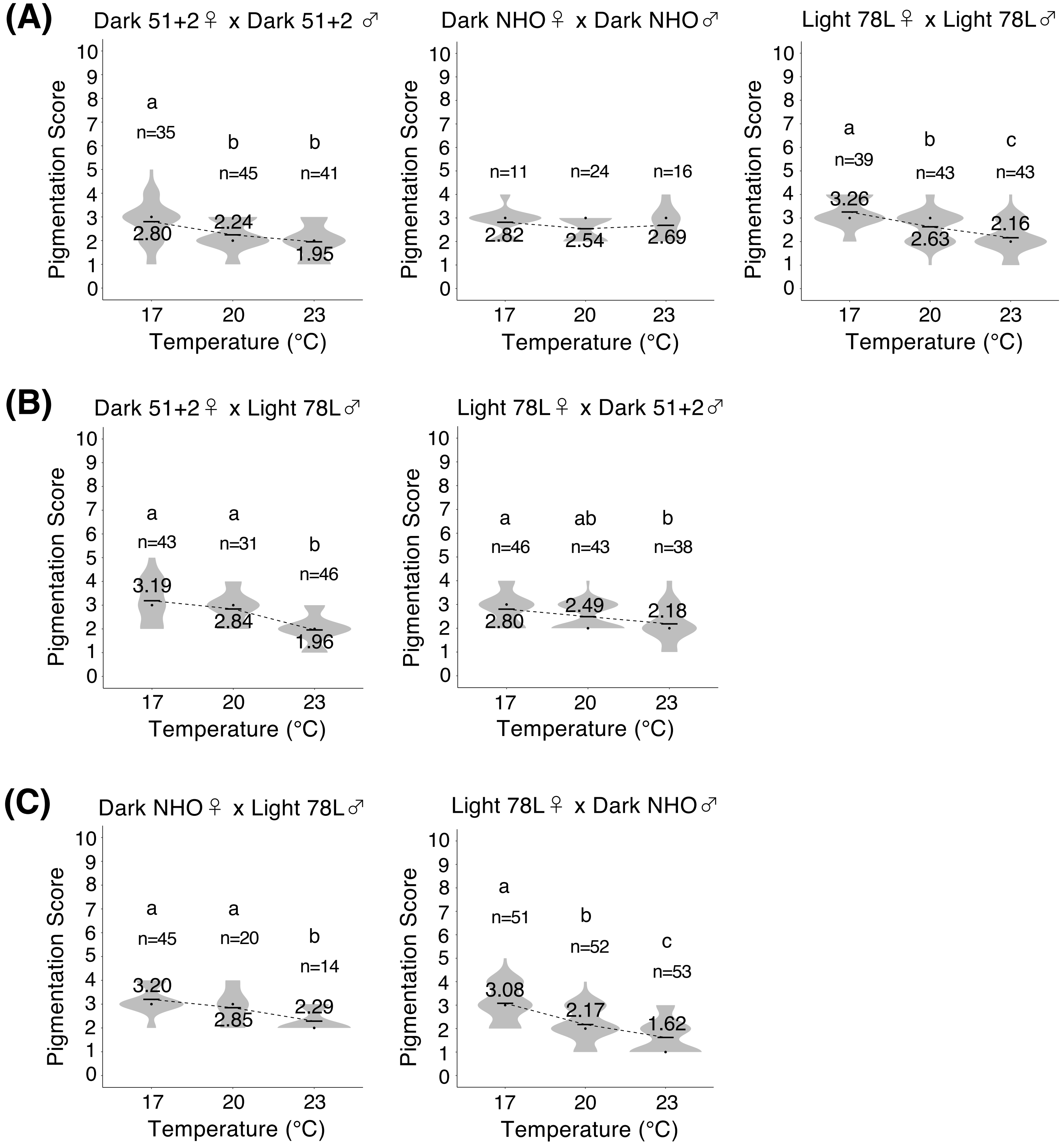

(A)
Dark 51+2♀ x Dark 51+2 ♂
Dark NHO♀ x Dark NHO♂
Light 78L♀ x Light 78L♂
(B)
Dark 51+2♀ x Light 78L♂
Light 78L♀ x Dark 51+2♂
(C)
Dark NHO♀ x Light 78L♂
Light 78L♀ x Dark NHO♂
Pigmentation Score
Temperature (°C)


Figure. S21

Figure. S21

The pigmentation scores for A5 segments of male abdomens of *D. jambulina*, reared at 17 °C, 20 °C, and 23 °C. (A) The scores for homozygotes from Dark 51+2 strain (left), Dark NHO (middle), Light 78L strain (right). There were significant differences between temperatures except Dark NHO homozygotes (one-way ANOVA, degree of freedom = 2). $F = 12.44$, $p < 10^{-4}$ for Dark 51+2 homozygotes. $F = 28.07$, $p < 10^{-10}$ for Light 78L homozygotes. (B) The scores for F1 hybrids from the crosses between females from Dark 51+2 strain and males from Light 78L strain (left), and between females from Light 78L strain and males from Dark 51+2 strain (right). There were significant differences between temperatures (one-way ANOVA, degree of freedom = 2). $F = 30.27$, $p < 10^{-10}$ for hybrids between Dark 51+2 females and Light 78L males. $F = 8.949$, $p < 10^{-3}$ for hybrids between Light 78L females and Dark 51+2 males. (C) The scores for F1 hybrids from the crosses between females from Dark NHO strain and males from Light 78L strain (left), and between females from Light 78L strain and males from Dark NHO strain (right). There were significant differences between temperatures (one-way ANOVA, degree of freedom = 2). $F = 12.21$, $p < 10^{-4}$ for hybrids between Dark NHO females and Light 78L males. $F = 16.32$, $p < 10^{-6}$ for hybrids between Light 78L females and Dark NHO males. Black bars and black dots indicate mean values and median values for each. Mean values are written near the black bars. Different alphabets indicate significant differences ($p < 0.05$, Tukey's HSD test).

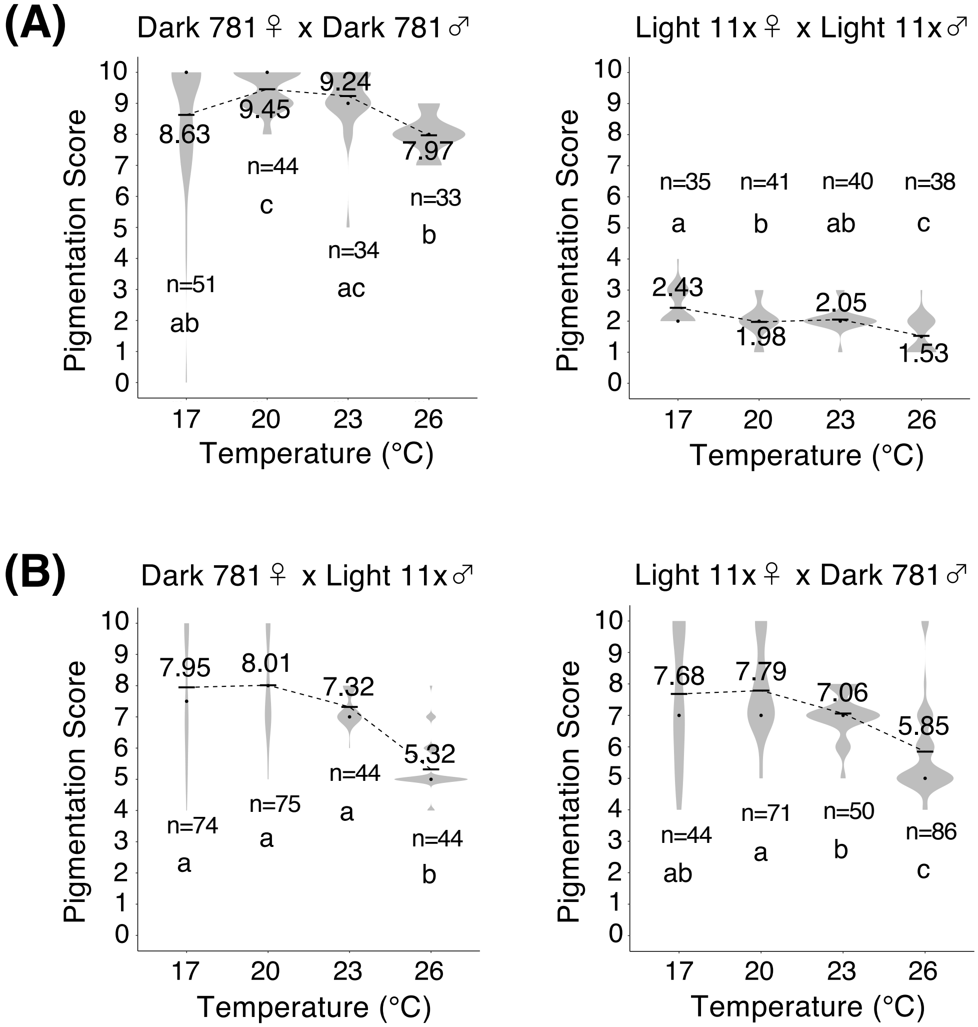


Figure. S22

The pigmentation scores for A5 segment of male abdomens of *D. burlai*, reared at 17 °C, 20 °C, 23 °C, and 26 °C. (A) The scores for Dark 781 homozygotes (left) and Light 11x homozygotes (right). There were significant differences between temperatures ($p < 10^{-5}$, one-way ANOVA, degree of freedom = 3). $F$ = 10.37 for Dark 781 homozygotes. $F$ = 18.39 for Light 11x homozygotes. (B) The scores for F1 hybrids from the crosses between females from Dark 781 strain and males from Light 11x strain (left), and between females from Light 11x strain and males from Dark 781 strain (right). There were significant differences between temperatures ($p < 10^{-10}$, one-way ANOVA, degree of freedom = 3). $F$ = 39.16 for hybrids between Dark 781 females and Light 11x males. $F$ = 27.96 for hybrids between Light 11x females and Dark 781 males. Black bars and black dots indicate mean values and median values for each. Mean values are written near the black bars. Different alphabets indicate significant differences ($p < 0.05$, Tukey's HSD test).

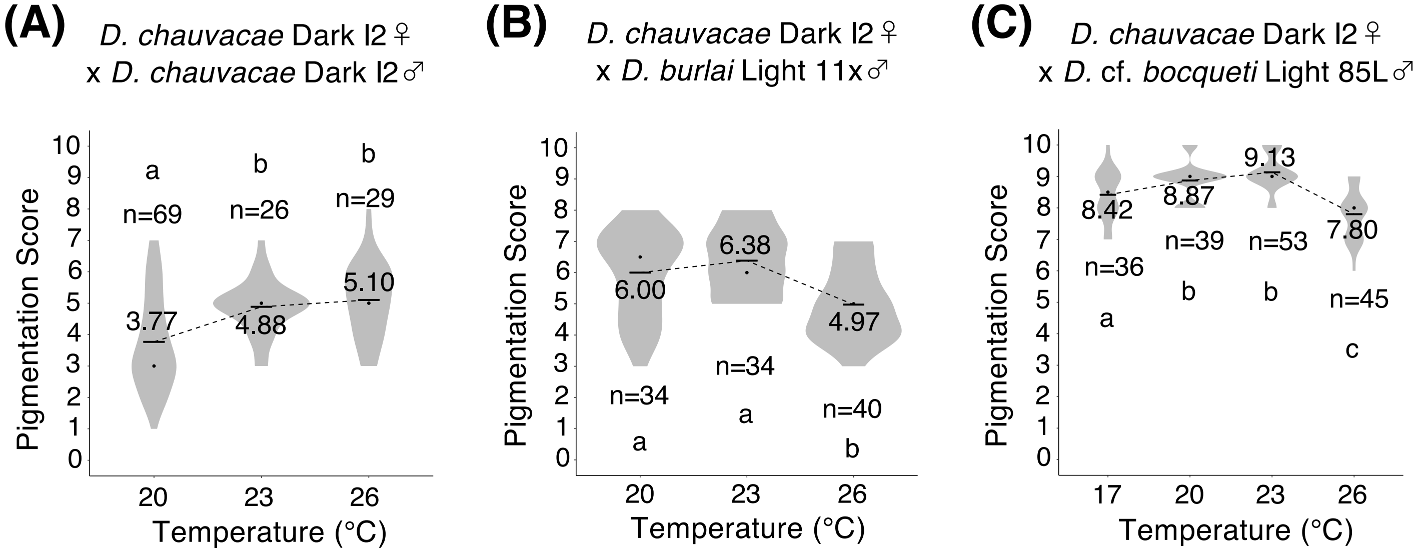


Figure. S23

The pigmentation scores for A5 segment of male abdomen of *D. chauvacae* and hybrids produced from interspecific crosses. (A) The score for Dark l2 homozygotes of *D. chauvacae*, reared at 20 °C, 23 °C, and 26 °C. There were significant differences between temperatures ($F$ = 13.21, $p < 10^{-7}$, one-way ANOVA, degree of freedom = 2). (B) The pigmentation scores for F1 hybrids from the crosses between *D. chauvacae* females from Dark l2 strain and *D. burlai* males from Light 11x strain. The hybrid flies were reared at 20 °C, 23 °C, and 26 °C. There were significant differences between temperatures ($F$ = 13.46, $p < 10^{-7}$, one-way ANOVA, degree of freedom = 2). (C) The pigmentation scores for F1 hybrids from the crosses between *D. chauvacae* females from Dark l2 strain and *D. bocqueti* males from Light 85L strain. The hybrid flies were reared at 17 °C, 20 °C, 23 °C, and 26 °C. There were significant differences between temperatures ($F$ = 31.92, $p < 10^{-10}$, one-way ANOVA, degree of freedom = 3). Black bars and black dots indicate mean values and median values for each. Mean values are written near the black bars. Different alphabets indicate significant differences ($p < 0.05$, Tukey's HSD test).
