## Supplemental Tables for "Evolution of dominance in a Mendelian trait is linked to the evolution of environmental plasticity"

**Table S1**

The results of diagnostic PCR on 43 isofemale strains of *D. rufa* sampled at Dogo Park in Matsuyama, Japan (Lat: 33.848338, Long: 132.786791). Dark 10_7Y, a Dark homozygous strain, and Light 59BY, a Light homozygous strain, were used as controls. A single female from each of the strains listed in the second column was genotyped; their phenotypes are indicated in the third column. The intronic region between 1st and 2nd exons of *pdm3* was targeted with two different sets of primers, “Dark”, (forward: CCATACCATACAAGCGCCATCTAG, reverse: TTATCATGCATCAATGCGACAGCAAC), which only amplifies the Dark *pdm3* allele, and “Light” (forward: same as “Dark”, reverse: AGCGACACAGATAACCAGATTTCGA), which only amplifies the Light *pdm3* allele. “+” in the “Dark primer set” column indicates that the 1.5 kb PCR product diagnostic of the Dark *pdm3* allele was amplified. “+” in “Light primer set” column indicates that the 656 bp PCR product diagnostic of the Light *pdm3* allele. “-” indicates that the corresponding PCR product was not observed.

| Individual No. | Strain | Phenotype | Dark primer set | Light primer set |
| --- | --- | --- | --- | --- |
| 1 | 17_7Y | dark | + | + |
| 2 | 6B | dark | + | - |
| 3 | 75BY | dark | + | + |
| 4 | 26B | dark | + | - |
| 5 | 17_28B | dark | + | - |
| 6 | 17_18B | dark | + | + |
| 7 | 36B | dark | + | - |
| 8 | 12B | dark | + | - |
| 9 | 17_40B | dark | + | + |
| 10 | 17_13Y | dark | + | + |
| 11 | 35B | dark | + | - |
| 12 | 1B | dark | + | - |
| 13 | 51BY | dark | + | + |
| 14 | 64BY | dark | + | - |
| 15 | 71BY | dark | + | + |
| 16 | 20B | dark | + | - |
| 17 | 15B | dark | + | - |
| 18 | 33B | dark | + | - |
| 19 | Dark 10_7Y | dark | + | - |
| 20 | 27B | dark | + | + |
| 21 | 24B | dark | + | - |
| 22 | Light 59BY | light | - | + |
| 23 | 3B | light | - | + |
| 24 | 58BY | light | - | + |
| 25 | 10_13_B | light | - | + |
| 26 | 17_22B | light | - | + |
| 27 | 17_33B | light | - | + |
| 28 | 50BY | light | - | + |
| 29 | 17_4Y | light | - | + |
| 30 | 73BY | light | - | + |
| 31 | 17B | light | - | + |
| 32 | 63BY | light | - | + |
| 33 | 79BY | light | - | + |
| 34 | 10_15_B | light | - | + |
| 35 | 17_14Y | light | - | + |
| 36 | 41Y | light | - | + |
| 37 | 54BY | light | - | + |
| 38 | 69BY | light | - | + |
| 39 | 40Y | light | - | + |
| 40 | 17_16B | light | - | + |
| 41 | 3B | light | - | + |
| 42 | 5BY | light | - | + |
| 43 | 17_9Y | light | - | + |

**Table S2**

Genotyping in *D. jambulina*. A single female from each of the isofemale strains listed in the second column was genotyped. Other than the Dark NHO strain, all strains were collected at the Chandigarh University campus. The second column indicates whether the isofemale strain was monomorphic (“mono dark” or “mono light”) or polymorphic (“poly”). The phenotypes of the genotyped individuals are indicated in the third column. For collecting dark virgin females, we selected individuals with the darkest phenotype among flies reared at 25 °C. For light virgin females, we selected individuals with the lightest phenotype among flies reared at 17 °C. The fourth column indicates the method of genotyping.

| Individual No. | Strain | Phenotype | Method |
| --- | --- | --- | --- |
| 1 | 14D (poly) | dark | amplicon sequencing |
| 2 | 16D (poly) | dark | amplicon sequencing |
| 3 | 29D (poly) | dark | amplicon sequencing |
| 4 | 54L (poly) | dark | amplicon sequencing |
| 5 | 51L (poly) | dark | amplicon sequencing |
| 6 | 27D (poly) | dark | amplicon sequencing |
| 7 | 59L (poly) | dark | amplicon sequencing |
| 8 | 74L (poly) | dark | amplicon sequencing |
| 9 | 56L (poly) | dark | amplicon sequencing |
| 10 | 76L (poly) | dark | amplicon sequencing |
| 11 | 63L (poly) | dark | amplicon sequencing |
| 12 | Dark NHO (mono dark) | dark | amplicon sequencing |
| 13 | 104D (poly) | dark | amplicon sequencing |
| 14 | 60L (poly) | dark | diagnostic PCR |
| 15 | 103D (mono dark) | dark | diagnostic PCR |
| 16 | Dark 51+2 (mono dark) | dark | diagnostic PCR |
| 17 | 70L (poly) | light | amplicon sequencing |
| 18 | 56L (poly) | light | amplicon sequencing |
| 19 | 76L (poly) | light | amplicon sequencing |
| 20 | 51L (poly) | light | amplicon sequencing |
| 21 | 24D (poly) | light | amplicon sequencing |
| 22 | 16D (poly) | light | amplicon sequencing |
| 23 | 58L (mono light) | light | amplicon sequencing |
| 24 | 14D (poly) | light | amplicon sequencing |
| 25 | 54L (poly) | light | amplicon sequencing |
| 26 | 59L (poly) | light | amplicon sequencing |
| 27 | 32D (poly) | light | amplicon sequencing |
| 28 | 66L (poly) | light | amplicon sequencing |
| 29 | 9D (poly) | light | amplicon sequencing |
| 30 | Light 78L (mono light) | light | diagnostic PCR |
| 31 | 71L (poly) | light | diagnostic PCR |
| 32 | 74L (poly) | light | diagnostic PCR |

**Table S3**

The combination of crosses and rearing temperatures. Females from the strains written in “Females” column were crossed with males from the strains written in the “Males” column on the same row. The rearing temperatures are indicated in “Temperature (°C)” column. Some species were not viable at all temperatures.

| Species | Females | Males | Temperatures (°C) |
| --- | --- | --- | --- |
| *D. rufa* | Dark 10_7Y | Dark 10_7Y | 20, 23, 26 |
|  | Light 59BY | Light 59BY | 20, 23, 26 |
|  | Dark 10_7Y | Light 59BY | 20, 23, 26 |
|  | Light 59BY | Dark 10_7Y | 20, 23, 26 |
| *D. burlai* | Dark 781 | Dark 781 | 17, 20, 23, 26 |
|  | Light 11x | Light 11x | 17, 20, 23, 26 |
|  | Dark 781 | Light 11x | 17, 20, 23, 26 |
|  | Light 11x | Dark 781 | 17, 20, 23, 26 |
| *D.* cf. *bocqueti* | Dark l1 | Dark l1 | 17, 20, 23, 26 |
|  | Dark y4 | Dark y4 | 17, 20, 23, 26 |
|  | Light 85L | Light 85L | 17, 20, 23, 26 |
|  | Dark l1 | Light 85L | 17, 20, 23, 26 |
|  | Light 85L | Dark l1 | 17, 20, 23, 26 |
|  | Dark y4 | Light 85L | 17, 20, 23, 26 |
|  | Light 85L | Dark y4 | 17, 20, 23, 26 |
| *D. jambulina* | Dark 51+2 | Dark 51+2 | 17, 20, 23 |
|  | Dark NHO | Dark NHO | 17, 20, 23 |
|  | Light 78L | Light 78L | 17, 20, 23 |
|  | Dark 51+2 | Light 78L | 17, 20, 23 |
|  | Light 78L | Dark 51+2 | 17, 20, 23 |
|  | Dark NHO | Light 78L | 17, 20, 23 |
|  | Light 78L | Dark NHO | 17, 20, 23 |
| *D. chauvacae* | Dark l2 | Dark l2 | 20, 23, 26 |
| *D. chauvacae* x  *D. burlai* | Dark l2 | Light 11x | 20, 23, 26 |
| *D. chauvacae* x  *D.* cf. *bocqueti* | Dark l2 | Light 85L | 17, 20, 23, 26 |

**Table S4**

Genotyping results in *D. jambulina*. A fragment of the 1st intron of *pdm3* was amplified and sequenced from light and dark virgin females sampled from 19 strains. For genotyping, dark virgin females from 13 different strains and light virgin females from 13 different strains were used. The alleles and positions of indels and SNPs within this amplicon are indicated. Nucleotides in black letters indicate insertions or SNPs, “-” indicates deletions, and gray letters indicate invariant nucleotides flanking the indels. The nucleotide 3 bp upstream of the 8-bp indel is defined as Position 1. All indels and SNPs are completely fixed between light and dark females.

|  | Position | | | | | | | | | | | | | | | | |
| --- | --- | --- | --- | --- | --- | --- | --- | --- | --- | --- | --- | --- | --- | --- | --- | --- | --- |
|  | 1 | 2 | 3 | 4 | 5 | 6 | 7 | 8 | 9 | 10 | 11 | 12 | 13 | 14 | 15 | 16 | 17 |
| dark females  (13 strains) | T | C | A | - | - | - | - | - | - | - | - | T | T | A | T | A | A |
| light females  (13 strains) | T | C | A | G | G | A | C | T | A | G | A | T | T | A | T | A | T |

|  | Position | | | | | | | | | | | | | | |
| --- | --- | --- | --- | --- | --- | --- | --- | --- | --- | --- | --- | --- | --- | --- | --- |
|  | 30 | 34 | 35 | 55 | 59 | 66 | 67 | 77 | 92 | 97 | 120 | 141 | 145 | 154 | 162 |
| dark females  (13 strains) | C | C | T | T | T | A | C | - | G | A | G | A | - | T | G |
| light females  (13 strains) | G | T | G | C | C | C | T | T | T | T | A | G | G | C | C |

**Table S5**

The results of selective genotyping in the *D.* cf. *bocqueti* introgression line. A 770 bp PCR amplicon around Exon 11 of *pdm3* contains seven single nucleotide polymorphisms (SNPs) that distinguish the dark parental strain Dark l1 and the light parental strain Light 85L. SNP alleles and positions relative to the 5’ end of the amplicon are indicated in the “Position” columns. All dark females from the introgression line have the same sequence as the Dark l1 parental strain, and light introgression females have the same sequence as the Light 85L strain.

| Individuals | Position | | | | | | |
| --- | --- | --- | --- | --- | --- | --- | --- |
|  | 182 | 190 | 203 | 219 | 276 | 407 | 427 |
| Dark l1 females (n = 4) | C | T | T | C | C | G | T |
| Light 85L females (n = 4) | G | C | G | T | T | A | C |
| Dark intro females (n = 19) | C | T | T | C | C | G | T |
| Light intro females (n = 18) | G | C | G | T | T | A | C |

**Table S6**

The results of two-way ANOVA on A6 pigmentation scores in heterozygous females of *D.* cf. *bocqueti*. Four “Genotypes” were used in the analyses: F1 females from the crosses between Light 85L females and Dark l1 males, between Dark l1 females and Light 85L males, between Light 85L females and Dark y4 males, and between Dark y4 females and Light 85L males. Two “Genotypes” (same combination of strains, but different direction of crosses) were compared in each contrast. “G x E” indicates an interaction between “Genotype” and “Temperature”. Sum Sq: sum of squares, Df: degrees of freedom, ***: *p* < 10^-10^. NS: not significant.

| Effect | Sum Sq | Df | *F* value | *p* value |
| --- | --- | --- | --- | --- |
| **Comparison of “Light 85L ♀ x Dark l1 ♂︎” & “Dark l1 ♀ x Light 85L ♂︎”** | | | | |
| Genotype | 1 | 1 | 0.52 | NS |
| Temperature | 3536 | 3 | 785.73 | *** |
| G x E | 2 | 3 | 0.52 | NS |
| Residuals | 522 | 348 |  |  |
| **Comparison of “Light 85L ♀ x Dark y4 ♂︎” & “Dark y4 ♀ x Light 85L ♂︎”** | | | | |
| Genotype | 3 | 1 | 2.18 | NS |
| Temperature | 5456 | 3 | 1247.39 | *** |
| G x E | 1 | 3 | 0.12 | NS |
| Residuals | 567 | 389 |  |  |

**Table S7**

The results of two-way ANOVA on A6 pigmentation scores in heterozygous females of *D. jambulina*. Four “Genotypes” were used in the analyses: F1 females from the crosses between Light 78L females and Dark 51+2 males, between Dark 51+2 females and Light 78L males, between Light 78L females and Dark NHO males, and between Dark NHO females and Light 78L males. Two “Genotypes” (same combination of strains, but different direction of crosses) were compared in each contrast. “G x E” indicates an interaction between “Genotype” and “Temperature”. Sum Sq: sum of squares, Df: degrees of freedom, ***: *p* <10^-10^, NS: not significant.

| Effect | Sum Sq | Df | *F* value | *p* value |
| --- | --- | --- | --- | --- |
| **Comparison of “Light 78L ♀ x Dark 51+2 ♂︎” & “Dark 51+2 ♀ x Light 78L ♂︎”** | | | | |
| Genotype | 3.4 | 1 | 0.59 | NS |
| Temperature | 1851.5 | 2 | 161.70 | *** |
| G x E | 13.2 | 2 | 1.15 | NS |
| Residuals | 2272.8 | 397 |  |  |
| **Comparison of “Light 78L ♀ x Dark NHO ♂︎” & “Dark NHO ♀ x Light 78L ♂︎”** | | | | |
| Genotype | 2.0 | 1 | 0.31 | NS |
| Temperature | 766.9 | 2 | 58.90 | *** |
| G x E | 12.5 | 2 | 0.96 | NS |
| Residuals | 1881.3 | 289 |  |  |
